# Large-scale single-cell phylogenetic mapping of clonal evolution in the human aging esophagus

**DOI:** 10.1101/2025.10.11.681805

**Authors:** Tamara Prieto, Dennis J. Yuan, John Zinno, Clayton Hughes, Nicholas Midler, Sheng Kao, Jani Huuhtanen, Ramya Raviram, Foteini Fotopoulou, Neil Ruthen, Srinivas Rajagopalan, Joshua S. Schiffman, Andrew R. D’Avino, Sang-Ho Yoon, Jesus Sotelo, Nathaniel D. Omans, Noelle Wheeler, Alejandro Garces, Barun Pradhan, Alexandre Pellan Cheng, Nicolas Robine, Catherine Potenski, Katharine Godfrey, Nobuyuki Kakiuchi, Akira Yokoyama, Seishi Ogawa, Julian Abrams, Ivan Raimondi, Dan A. Landau

## Abstract

The human somatic genome evolves throughout our lifespan, producing mosaic individuals comprising clones harboring different mutations across tissues. While clonal expansions in the hematopoietic system have been extensively characterized and reported to be nearly ubiquitous, clonal mosaicism (CM) has more recently also been described across multiple solid tissues. However, outstanding questions remain about the parameters and processes of human somatic evolution in non-cancerous solid human tissues, including when clones arise, how they evolve over time, and what mechanisms lead to their expansion. Questions of timing and clonal dynamics can be addressed through phylogenetic reconstruction, which serves as a ‘temporal microscope’, while uncovering the mechanisms of expansion necessitates simultaneous phenotypic profiling. To address this gap, here we develop Single-cell Miniaturized Automated Reverse Transcription and Primary Template-directed Amplification (SMART-PTA) for joint single-cell whole-genome and whole-transcriptome sequencing for large scale and cost efficient interrogation of solid tissue CM. We established a workflow that generates hundreds of matched single-cell whole genome and transcriptome libraries within a week. We profiled phenotypically normal esophagus tissue from four aged donors and used somatic variants to build high-resolution single-cell lineages from >2,700 cells with accompanying transcriptomic information, reconstructing >70 years of somatic evolution. T cell expansions identified from T cell receptor (TCR) sequences validated the clonal structure of the single-nucleotide variant (SNV)-based phylogenies and phylogenetic cross-correlation analysis showed that epithelial cells had higher degrees of shared ancestry by spatial location compared to immune cells. Mapping mutation signatures to the phylogenetic tree revealed the emergence of tobacco/alcohol exposure-related signatures later in life, consistent with the donors’ exposure histories. We identified variants in driver genes that were previously reported in the phenotypically normal esophagus, detecting clonal expansions harboring mutations in genes including *TP53* and *FAT1*. We mapped the evolution of clones with both monoallelic as well as biallelic *TP53* loss, including a clone associated with high expression of cell cycling genes and higher chromosome instability. Leveraging the matched transcriptome data, we uncovered cell type biases in mutant clones, with a higher proportion of *TP53* or *FAT1*-mutant cells in an earlier basal epithelial cell state compared to wild-type cells. We further observed copy-neutral loss of heterozygosity (CNLOH) events on chromosome 9q that spanned the *NOTCH1* locus in up to ∼35% of epithelial cells. Mapping CNLOH events to the phylogenetic tree revealed a striking pattern in which CNLOH was separately acquired many times, reflecting convergent evolution. Cells with CNLOH events were biased towards the earlier basal epithelial state, suggestive of a selective advantage that leads to prevalent recurrence of chr9q CNLOH. Together, we demonstrate that SMART-PTA is an efficient, scalable approach for single-cell whole-genome and whole-transcriptome profiling to build phenotypically annotated single-cell phylogenies with enough throughput and power for application to normal tissue somatic evolution. Moreover, we reconstruct the evolutionary history of the esophageal epithelium at high scale and resolution, providing a window into the dynamics and processes that shape clonal expansions in phenotypically normal tissues throughout a lifespan.

## Introduction

Somatic mosaicism is an exciting, novel frontier in the study of the human genome. Somatic mutations accumulate throughout our lifetime, with clonal evolution leading to significant mosaicism across tissues in physiological aging and disease^1,2^. For example, the phenotypically normal esophagus was shown to be replaced with somatically mutated clones with aging and environmental exposures^3,4^. While recent bulk sequencing methods based on duplex sequencing can now detect very rare somatic mutations and resolve the mutation landscape of some small clones^5^, they do not always allow for complete clonal deconvolution^6^. Single-cell whole-genome sequencing (scWGS) has the potential to deliver the highest resolution somatic profiles and directly identify sets of mutations present in cell clones of any size, without relying on inferences based on mutation frequency^7,8^, providing a new window into clonal evolution and human somatic mosaicism across tissues.

Excitingly, such clonal evolution can be defined through cell lineage tracing technologies, which provide a ‘temporal microscope’ to chart the evolution of somatic cells within tissues or organisms, allowing to infer back in time how cell populations and their phenotypes evolve. Such analysis calls for approaches that map phylogenetic relationships between cells by leveraging molecular barcodes, which can be exogenous (synthetic) or endogenous (intrinsic), allowing to define how tissues evolve over time^9–11^. In humans, only somatic endogenous variation can be leveraged to build cell lineages, where mutations act as intrinsic barcodes for phylogenetic reconstruction, in order to trace the history of mitotic relationships throughout time. This lineage tracing approach allows for phylodynamic analysis of the clonal landscape of somatic tissues directly from human samples, enabling the determination of when variants arose during an individual’s life and to what degree these variants promote clonal expansion. In high-aneuploid cancer cells, shallow scWGS has been used to profile copy-number variants (CNVs) to build tumor phylogenies^7,8,12,13^. In normal, copy-number stable tissues, CNV events are not frequent enough to result in high-resolution phylogenies, restricting analysis to only those rare cells that harbor aneuploidy^14^. As an alternative, lineage tracing using single-cell somatic single-nucleotide variants (SNVs) presents an advantageous approach in normal tissue. However, SNV lineage tracing has been historically challenging due to high rates of allelic dropout and base substitution errors occurring during exponential amplification intrinsic to PCR-based, multiple displacement amplification (MDA)-based and hybrid scWGS approaches^15–17^. These limitations commonly challenge the detection of shared informative genomic differences among cells, needed for phylogenetic reconstruction. In addition to high error and dropout rates, the cost of high-depth scWGS has limited application to ∼10s of cells per patient in most studies^16–18^.

To circumvent these amplification biases, WGS of single-cell derived colonies of primary cells^10,11^ has been used to obtain high-quality single cell-like data. However, this approach is limited in analysis of somatic evolution in solid tissues, where live cells are not readily available for in vitro expansion. Additionally, clonal expansion followed by WGS generally lacks opportunity to directly assess matching single-cell phenotypes from the primary cells, which can be annotated on phylogenies to simultaneously analyze cell ancestry with current cell state^19^ and map somatic genotype to clonal phenotype.

Recently, primary template-directed amplification (PTA) has enabled scWGS with lower allelic dropout and reduced nucleotide substitution rates^20^ due to its non-exponential nature, and has been successfully applied to characterize genetic diversity in primary human cells^19,21–23^. While powerful, the inherent low throughput of this method prevents large-scale application to the analysis of somatic variation in normal solid human tissues in which, unlike blood, clones are spatially restricted and multiple cells from the same clone are only infrequently sampled using standard size biopsies and low sampling rates. Thus, analysis of the mechanisms of solid tissue somatic evolution that occurs over decades directly from primary samples requires direct scWGS at scale, calling for novel single-cell workflows specifically conceptualized for phylogenetic reconstruction that enable analysis of large cell numbers and ability to co-capture cell phenotypes.

To address this gap, we developed SMART-PTA (<u>S</u>ingle-cell <u>M</u>iniaturized <u>A</u>utomated <u>R</u>everse <u>T</u>ranscription and <u>P</u>rimary <u>T</u>emplate-directed <u>A</u>mplification), enabling high-throughput phylogenetic reconstruction of solid tissues, such as the phenotypically normal esophagus, at scale. By miniaturizing and automating the standard mRNA+PTA workflow, we were able to generate hundreds of single-cell whole genome and transcriptome libraries in a single run. We obtained high-quality SMART-PTA libraries with coverage and detection metrics comparable to or exceeding those of standard-PTA+RNA. To further increase throughput, enabling phylogenetic applications at scale, we adapted scWGS library preparation to be compatible with the mostly natural sequencing-by-synthesis technology from Ultima Genomics^24^, achieving genome sequencing with lower costs (0.8 USD/Gb). To assess precision and accuracy of our workflow, we sampled single cells from an *in vitro* experimental clonal tree, recapitulating ground-truth lineage relationships.

To demonstrate the power of this approach for application to human solid tissue, we applied this workflow to investigate somatic mosaicism in the phenotypically normal esophagus from elderly donors (>76 years of age), a tissue in which somatic clones dramatically increase with aging and environmental exposures^3,4^. Using this method, we generated matched single-cell whole genomes and full-length transcriptomes from 2,783 cells. We built high-resolution single-cell phylogenies from somatic SNVs and annotated each cell with gene expression and cell type information. Using these large phylogenies, we traced lineage relationships and timed the most recent common ancestor between epithelial and immune cells to early development, with phylogenetic reconstruction accurately reflecting the distinct origins of epithelial versus tissue-resident immune cells. Mutation signature analysis along the phylogeny identified dynamic patterns of exposure-related signatures. We identified driver mutations and observed increased proliferation of mutant clones with biallelic *TP53* loss. Clones harboring loss-of-function driver variants in *TP53* or *FAT1* displayed earlier differentiation biases, with significantly higher proportions of cells in the basal epithelial state compared to wild-type cells. Finally, we detected prevalent, recurrent copy neutral loss of heterozygosity (CNLOH) events on chromosome 9q containing the *NOTCH1* gene. Notably, multiple CNLOH events arose independently, affecting both maternal and paternal alleles, and skewing cells towards the earlier basal epithelial state. In summary, we generated large, phenotypically annotated single-cell phylogenies from primary human esophagus tissue to dissect clonal origins, growth patterns and phenotypic impact in somatic evolution.

## Results

### SMART-PTA generates high-quality whole-genome + full-length RNA sequencing from single cells at high throughput

To investigate the mechanisms driving somatic evolution in normal human solid tissues across the lifespan, phylogenetic approaches applied to single-cell genomic data have emerged as powerful tools, offering critical insights into developmental history and clonal dynamics^10,11,25^. In order to dissect phenotypic changes dictated by genetic mosaicism, phylogenetic information needs to be coupled with matched transcriptome profiles. While methods that enable joint genome and transcriptome profiling in single cells are increasingly being adopted by the scientific community, they typically operate at low throughput, limited to dozen cells per donor^26–30^, making them prone to severe sampling bias and limited for comprehensive clonal reconstruction. To reduce the uncertainty of clonal time inferences over decades, we hypothesized that somatic phylogenetic studies would benefit from large-scale whole-genome sequencing of hundreds of primary human cells. Furthermore, the statistical power of analytical approaches that integrate phylogenetic inference with models of population dynamics (Phylodynamics) and the study of phenotypic variation across a phylogenetic tree (Comparative Phylogenetics), increases with sample size. To address this need, we developed the SMART-PTA workflow, by integrating advances in whole-genome amplification, library preparation and sequencing.

First, to ensure that scWGS with PTA is able to generate accurate phylogenies that faithfully reconstruct chronological time, we assessed whether we could robustly recapitulate the same evolutionary relationships and clonal structure using an *in vitro* evolutionary experiment where single-cell whole genomes were amplified with PTA. To ensure the reliability and compatibility of phylogenetic inference derived from PTA single-cell sequencing, we generated an *in vitro* clonal evolution model by iteratively isolating and expanding single-cell-derived clones of DLD-1 cells over 123 days (**Methods**). Over this time, the inherent genetic instability of this cell line leads to the accumulation of spontaneous mutations, resulting in subclonal diversification, mimicking the natural process of clonal evolution. At defined time points during the *in vitro* expansion, single cells were isolated to generate a ground truth phylogenetic tree with a known temporal scale (**Fig. 1A**). Somatic SNVs were then used to reconstruct a phylogenetic tree of the 16 sequenced DLD-1 single cells. The phylogenetic topology obtained based on patterns of shared and unique SNVs between clones matched the *in vitro* experimental design with 100% accuracy (**Fig. 1B, Methods**). As we are not only interested in the topology or relationships of the cells but also in the of time origin of the different clones, we compared somatic mutation burden of independent clones across experimental time points, obtaining high correlation between evolutionary rates (in SNV counts) and sampling time (Pearson’s R = 0.78, *P* = 3.6 × 10^-4^). Together, these analyses provide confidence that PTA-based scWGS can be used for phylogeny building from SNVs.

**Figure 1:**
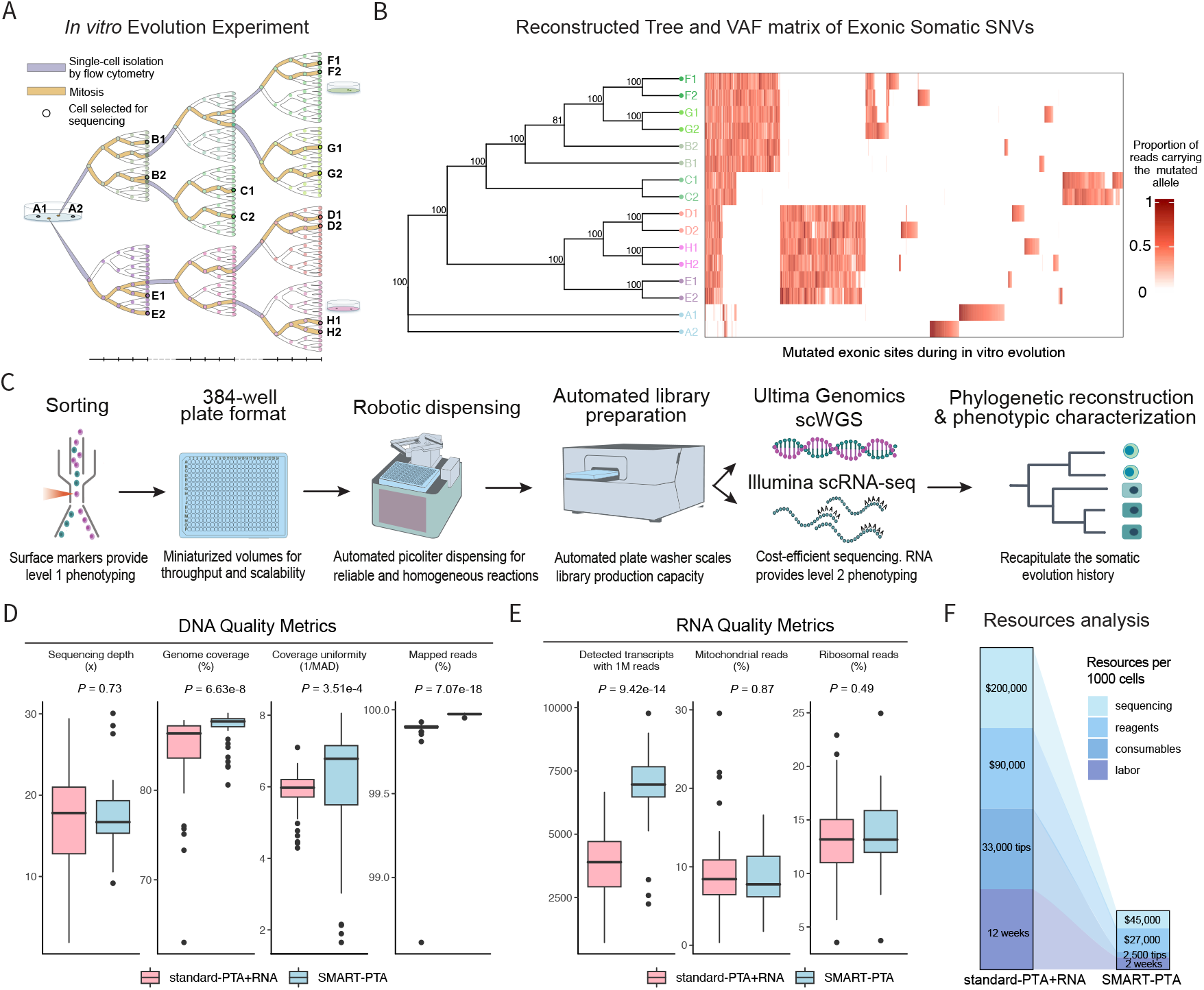
Scalable joint single-cell capture of whole genomes and full-length transcriptomes. **A**, *In vitro* evolution experiment using DLD-1 cells that were cultured and split over four months. We isolated 16 single cells (indicated with bold circles and labeled) across timepoints and performed single-cell whole-genome sequencing using mostly natural sequencing-by-synthesis chemistry from Ultima Genomics (>15× coverage depth). **B**, Phylogenetic reconstruction based on single-cell whole-genome amplification with UG reflects the topology of the *in vitro* experiment in (A). Variant allele frequency (VAF) of exonic somatic variants is indicated, displayed as the proportion of reads carrying the mutated allele per cell. Bootstrap support is annotated in the internal nodes of the tree. **C**, Schematic for single-cell miniaturized automated reverse transcription and Primary Template-directed Amplification (SMART-PTA). First, cell populations of interest are isolated with flow cytometry using cell surface markers to provide level 1 phenotyping. Single cells are sorted into 384-well plates, enabling miniaturized reaction volumes for increased throughput and scalability. Reagents are dispensed in picoliter volumes using robotics to allow for consistency and homogeneous reactions. Library preparation uses automated plate washing to scale production capacity. Ultima Genomics provides cost-efficient single-cell whole-genome sequencing, and Illumina single-cell RNA-seq provides level 2 phenotyping to assign cells states. Finally, phylogenetic reconstruction and phenotypic characterization analysis recapitulates the somatic evolution history of the analyzed cells. **D**, Quality metrics for whole-genome amplification libraries for standard-PTA versus SMART-PTA calculated for 50 cells processed with each protocol, comparing sequencing depth, genomic coverage (percentage), genomic coverage uniformity [1/median absolute deviation (MAD)] and percentages of mapped reads. *P* values from two-sided Wilcoxon test. Box plots represent the median, bottom and upper quartiles; whiskers correspond to 1.5 times the interquartile range. **E**, Quality metrics for transcriptome libraries for standard versus SMART-PTA calculated for 50 cells processed with each protocol, comparing detected transcripts in cells with >1 million reads, percentage of mitochondrial reads and percentage of ribosomal RNA reads. *P* values from two-sided Wilcoxon test. Box plots represent the median, bottom and upper quartiles; whiskers correspond to 1.5 times the interquartile range. **F**, Comparison of cost (for sequencing, reagents and consumables) and time expenditures (labor) required to profile 1,000 cells using standard-PTA+RNA versus SMART-PTA. Cost estimates reflect pre-2025 prices.

Second, to further develop a framework that enables broader, more cost-effective adoption of large-scale single-cell phylodynamic studies, we also reasoned that we could leverage the recent high-throughput sequencing technology developed by Ultima Genomics (UG). By using mostly natural sequencing-by-synthesis chemistry, the UG platform reduces sequencing costs to 0.8 USD per Gb^24^. Hypothesizing that sequencing at 15× depth could further reduce sequencing costs by half while preserving phylogenetic accuracy, we implemented a joint calling strategy. Although lower coverage may decrease detection of cell-unique variants, which mainly affects estimates of external branch lengths, joint calling of two or more cells provides standard coverage for detection of shared somatic mutations that define tree topology. Thus, at 15× we can achieve sensitivity for internal branches comparable to standard 30× depth, while external branch lengths can be corrected using per-cell recall estimates (**Methods**). Moreover, we have shown that recall in standard-PTA libraries does not fully plateau with increasing sequencing depth but approaches saturation, with only a modest gain when increasing from 15× to 30×^31^. To assess single-cell SNV calling with UG, we sorted and amplified two cells from each of the seven clones and the parental line from the *in vitro* experiment with standard-PTA, generating both Illumina and UG libraries (**Supplementary Fig. 1A**). The analysis of UG libraries identified 52,287.25 ± 3,188.25 (mean ± standard deviation) somatic SNVs per cell. Of these, >95% of somatic SNVs per cell were also observed in matching Illumina libraries (**Supplementary Fig. 1B**), while <5% somatic SNVs per cell were found to be UG specific. To further characterize UG-specific mutations, we assessed their trinucleotide mutation profile, revealing almost identical 96-context proportions (cosine similarity = 0.99) to the shared SNVs (**Supplementary Fig. 1C**). Mutational signature analysis showed that SNVs exhibited strong contributions from clock-like (SBS5) and microsatellite instability-associated signatures (SBS6, SBS14, SBS15, SBS20, SBS21; COSMIC V3; **Supplementary Fig. 1D**), consistent with the known mutational landscape of this cell line. This supports the interpretation that both shared SNVs and those uniquely detected by UG likely reflect true positive variants. Together, these results demonstrate that UG can be incorporated as a low-cost sequencing option into our single-cell workflow and that lowering sequencing depth per cell can aid scalability while maintaining phylogenetic accuracy.

Finally, while sequencing with UG and implementing joint calling effectively reduced the cost of generating single-cell whole genomes, the expense (in terms of reagents and experimental consumables) and the time required to produce whole genomes plus transcriptomes at scale from hundreds to thousands of single cells remain prohibitive. To address this challenge, we developed the <u>S</u>ingle-cell <u>M</u>iniaturized <u>A</u>utomated <u>R</u>everse <u>T</u>ranscription and <u>P</u>rimary <u>T</u>emplate-directed <u>A</u>mplification (SMART-PTA) workflow for high-throughput, cost-efficient single-cell whole-genome plus transcriptome sequencing using UG. SMART-PTA incorporates automated volume dispensing that is accurate to picoliter volumes for improved precision and throughput of reagent dispensing and mixing, together with automated plate washing for increased library preparation efficiency in 384-well plate formats (**Methods**) for single-cell whole-genome and transcriptome amplification (**Fig. 1C**). The transition from manual standard-PTA+RNA^29^ workflow to miniaturized robotic automated format significantly increases speed and the number of single cells that can be processed in a single run of the workflow. We further reduced reagent volumes allowing us to decrease reaction costs during whole-genome and cDNA amplification steps.

To ensure that the miniaturization process did not affect the sequencing quality, we prepared whole-genome sequencing libraries, using either standard-PTA+RNA or SMART-PTA, from 50 human esophageal epithelial primary cells with each protocol. Human esophagus tissue was obtained by endoscopic sampling of individuals affected by gastroesophageal reflux disease (GERD; donor details in **Supplementary Table 1**). The collected tissue was enzymatically dissociated to obtain a single-cell suspension (**Methods**). Epithelial (EPCAM+) and immune (CD45+) single cells were sorted into multiwell plates by fluorescence-activated cell sorting (FACS) to enable single-cell whole-genome and cDNA amplification.

The matched single-cell whole-genome and transcriptome amplification libraries were sequenced and analyzed in order to compare quality metrics. Each individual genomic library (n = 100 single cells total) was sequenced aiming for 15× coverage depth, resulting in similar yield for both protocols (median ± median absolute deviation (MAD) of

17.81 ± 4.90 for standard-PTA+RNA and 16.69 ± 2.93 for SMART-PTA). SMART-PTA outperformed standard-PTA+RNA in terms of DNA quality metrics (**Fig. 1D**), with comparable genome coverage (87.70 ± 0.52 for SMART-PTA vs 86.35 ± 1.41 for standard-PTA+RNA), coverage uniformity (0.85 ± 0.02 for SMART-PTA vs 0.83 ± 0.01 for standard-PTA+RNA) and mappability (99.97 ± 0.001 for SMART-PTA vs 99.89 ± 0.01% for standard-PTA+RNA). The analysis of the matched transcriptome libraries revealed a higher mean number of transcripts per cell using SMART-PTA (n = 6,893 transcripts) versus standard-PTA+RNA (n = 3,797 transcripts), while percentages of mitochondrial and ribosomal RNA reads from the two approaches were not significantly different (**Fig. 1E**). SMART-PTA also showed higher RNA gene body coverage compared to standard-PTA+RNA (**Supplementary Fig. 1E**). Taken together, these data demonstrate that the introduction of automated robotic library preparation and the miniaturization of reaction volumes do not affect the performance of the PTA+RNA protocol, enabling scalable processing of large numbers of single cells.

Collectively, our *in vitro* evolutionary experiments demonstrate the accuracy and specificity of UG sequencing of PTA-amplified single cells, supporting its application in building large-scale single-cell phylogenies from SNV data. Furthermore, we developed an integrated multiomic single-cell sequencing solution, addressing key barriers to scaled production of large single-cell phylogenies, including the high costs of assay reagents and sequencing, as well as long processing times. Through miniaturization and automation of the workflow, we reduced reagent costs per cell by more than 75%; we further decreased processing time for 1,000 single-cell whole genomes from three months to two weeks and significantly reduced consumable usages (as an illustrative example, pipette tips used decreased from 33,000 to 2,500 per 1,000 single-cell whole genomes). Thus, our automated workflow combined with cost-efficient sequencing delivers matched single-cell whole genomes and transcriptomes with reduced labor, cost and reagent use compared with standard processing (**Fig. 1F**), providing an avenue for high-throughput analysis of a high number of cells from human samples and enabling phylogenetic reconstruction of complex somatic evolution dynamics.

### Large-scale single-cell multiomic sequencing of normal human esophagus

To investigate clonal architecture and phylogenetic relationships at the single-cell level in the aging human esophagus, we generated single-cell whole-genome and transcriptome libraries of biopsies from four elderly donors (ages 76-79; **Supplementary Table 1**). For eso01, three adjacent biopsies from the middle thoracic region (approximately 30 cm from the incisors) were sampled and processed. For eso02-eso04, multi-regional sampling was conducted at four anatomically distinct locations, spanning a distance of 20 to 40 cm, measured from the incisors, in order to capture spatial heterogeneity along the proximal-distal axis (**Fig. 2A**). All the patients underwent endoscopy due to GERD, achalasia and/or for surveillance or evaluation of gastric intestinal metaplasia and related upper gastrointestinal symptoms (**Supplementary Table 1**). The donors had different levels of exposure to risk factors, including tobacco and alcohol, that are associated with esophageal mosaicism (**Fig. 2B**). Each tissue punch biopsy was dissociated using an established enzymatic digestion protocol optimized to preserve whole cell integrity (**Methods**), and single EPCAM+ epithelial and CD45+ immune cells were sorted. We generated high-depth whole-genome data (average 14.8×) from a total of 2,783 single cells across four donors (61 cells from one donor with standard-PTA+RNA), three with SMART-PTA (2,722 cells total); **Fig. 2B**), attaining high breadth of whole-genome coverage (83.8%) and reaching a median mappable genome coverage (excluding repeat-masked regions) higher than 90% across all patients (**Supplementary Table 2**). For subsequent analyses throughout the study, we implemented specific quality metrics and filtering stringencies, appropriate for each type of analysis, to ensure the robustness of the results (**Supplementary Fig. 2A, Methods**).

**Figure 2:**
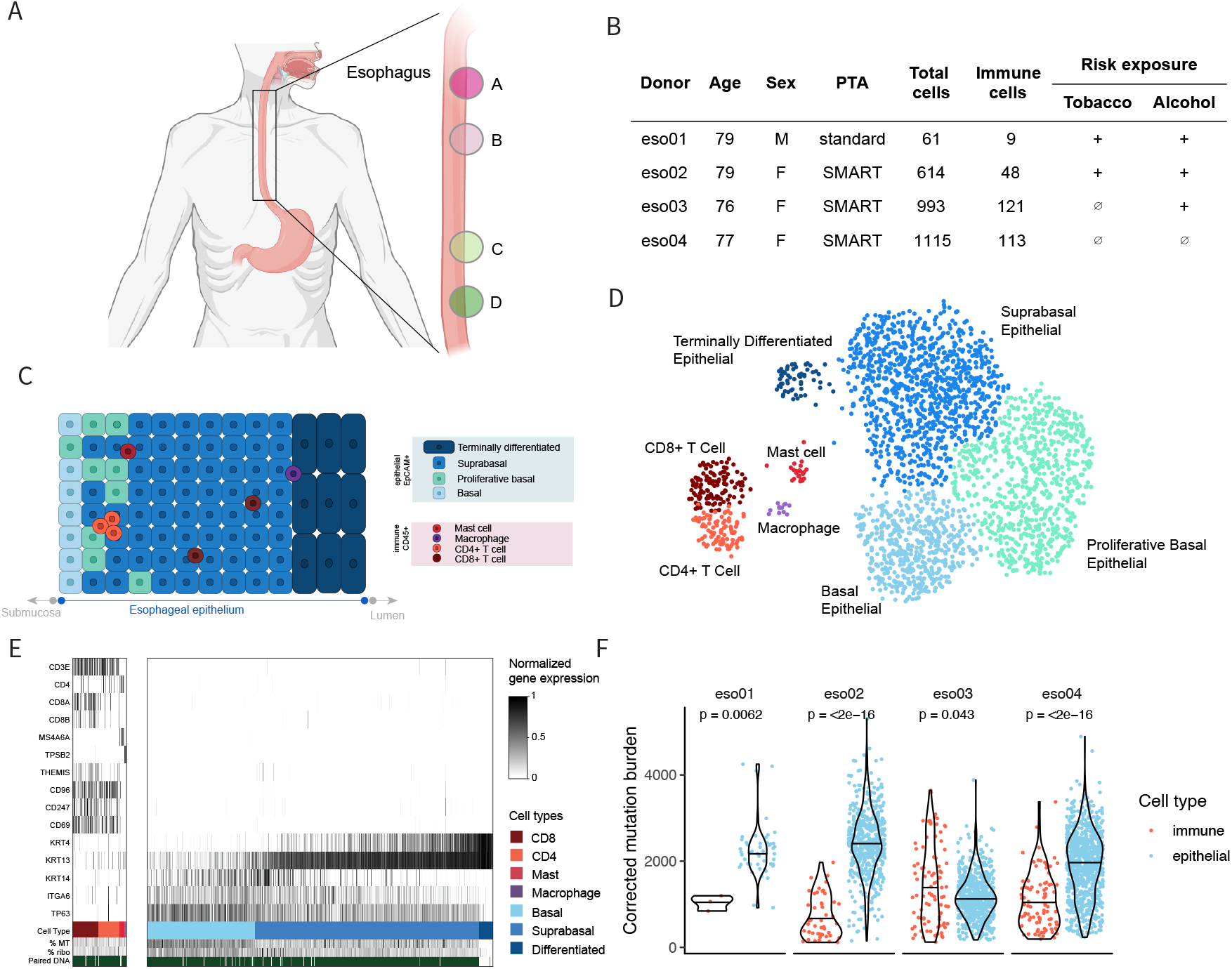
Large-scale single-cell genome and transcriptome sequencing of normal human esophagus. **A**, Schematic of the sampling strategy used to obtain tissue biopsy of the esophagus across four regions for three donors (eso02-eso04) processed and sequenced with SMART-PTA. **B**, Cohort description with donor, sequencing and risk exposure information (n = 2,783 cells). Detailed donor and sampling information can be found in Supplementary Table 1. **C**, Illustration of the cellular makeup of the esophagus, including epithelial and immune cell types, along with illustration of the stratification of the epithelial cells in esophageal layers. **D**, Uniform manifold approximation projection (UMAP) of 2,491 single cells with matched DNA and RNA libraries (∼10% of the total 2,783 cells were filtered out due to low RNA coverage; Supplementary Fig. 2A). Cells are colored by annotated cell type, including immune cells and epithelial cells, recapitulating the cells constituting the esophagus as illustrated in panel (C). **E**, Expression of marker genes underlying cell type assignment across single cells for immune cells (left) and epithelial cells (right). Percentage of mitochondrial (MT) reads, percentage of ribosomal (ribo) RNA reads and matched DNA sequencing reads are indicated. CD4 = CD4+ T cells, CD8 = CD8+ T cells. **F**, Mutation burden (estimated number of SNVs) derived from high-quality single-cell whole-genome sequencing of esophagus samples (this analysis summarizes 2,395/2,783 cells, excluding cells with potential contamination evidenced by >25% of cell-unique SNVs overlapping with dbSNP and those with low genome breadth and long external branch lengths; Methods) from four donors (eso01, n = 52 epithelial cells, n = 3 immune cells; eso02, n = 560 epithelial cells, n = 48 immune cells; eso03, n = 666 epithelial cells, n = 92 immune cells; eso04, n = 886 epithelial cells, n = 88 immune cells). Two-sided Wilcoxon test. To address variation in genome coverage, single-cell amplification and sequencing allelic imbalance was calculated using heterozygous germline SNPs and leveraged to correct the number of unique single-cell SNVs, correcting mutation burden estimates accordingly (Methods).

The human esophageal epithelium is composed of stratified layers of cells at distinct stages of differentiation (**Fig. 2C**). The basal layer, resting above the submucosa, harbors long-lived, quiescent basal stem cells. Just above, the epibasal layer contains proliferative basal cells that divide to supply the suprabasal layers^32^. These suprabasal cells undergo progressive differentiation as they migrate toward the lumen, ultimately forming a terminally differentiated cell barrier that is continuously shed to maintain epithelial homeostasis^33,34^. To achieve high resolution phenotypic characterization of the esophageal tissue that captures these layers, we used matched single-cell transcriptomes and performed unbiased clustering and annotation based on gene expression profiles. Analysis of the single-cell RNA libraries yielded a high-quality dataset, with a mean of over 5,300 genes detected per cell (**Supplementary Fig. 2B**; **Supplementary Table 2**). Additionally, the cells exhibited low mitochondrial gene content and minimal ribosomal RNA contamination (**Supplementary Fig. 2C, 2D**), consistent with high cellular integrity and efficient transcript capture. This enabled reliable identification and classification of the major cell types comprising the esophageal cellular landscape.

We leveraged the single-cell transcriptome data and canonical marker gene expression (**Supplementary Table 3**) to perform unsupervised clustering and assign cell type identities (**Fig. 2D, Supplementary Fig. 2E, 2F**). We were able to resolve the major epithelial and immune compartments, including distinct immune cell subsets (CD8+ T cells, CD4+ T cells, mast cells and macrophages) and epithelial subpopulations (basal, proliferative basal, suprabasal and terminally differentiated) (**Fig. 2E)**. We further leveraged the depth of our transcriptomic data to computationally infer cell cycle phase across cells (**Supplementary Fig. 2G**) as well as epithelial differentiation scores specifically within the EPCAM+ compartment (**Supplementary Fig. 2H**; **Methods**). To reconstruct the epithelial differentiation trajectory, we performed pseudotime analysis showing sequential expression of established differentiation markers, particularly lineage-specific keratins^32^, and a progression from basal to terminally differentiated states (**Supplementary Fig. 2I**; **Methods**), recapitulating known stages of squamous epithelial maturation^35^.

We integrated the matched genome-wide somatic variant profiles and transcriptomes to perform a phenotype-guided genomic characterization of both epithelial and immune lineages. This analysis revealed substantial inter- and intra-donor variability in somatic mutation load across both lineages (**Fig. 2F**). We found that for epithelial cells, the median number of mutations per cell ranged from 1,115-2,401, and for immune cells, ranged from 542-1,237. This difference could potentially be explained by distinct division rates for the different cell types, assuming a constant mutation rate. Basal epithelial cells in the esophagus have been reported to divide twice a week in mice^36^ and turn over every 11 days in humans^37^, whereas most immune cells originate from hematopoietic stem cells (HSCs) that divide once a year^38^, which would be consistent with the lower mutation burden observed in immune cells. However, tissue-resident immune cells can also self-renew and divide *in situ* upon response to injury or infection. Such responses can be highly variable across donors and may account for our observations in donor eso03, where tissue resident lymphocytes showed higher mutation burden (see more below). Overall, the epithelial mutation burdens broadly fell within the range of mutations estimated previously from a small number of single-cell epithelial colonies (n = 13 total, with 1-2 colonies per donor)^4^. Mutation burdens of immune cells, which comprise mostly T cells (**Fig. 2D**), also align with the previously reported values^39^. Our data capture cell-type resolved mutation burden from hundreds of cells per individual, and reveal the variability within donor esophageal epithelial cells that is missed when only assessing a small number of expanded clones.

### Reconstructing large, phenotypically annotated high-resolution phylogenetic trees in human somatic evolution with single-cell whole genomes and transcriptomes

To map somatic evolution over the lifespan, we next constructed phylogenies based on single-cell SNVs. The large size of our dataset presented a considerable analytical challenge, comparable to very large phylogenetic studies of hundreds to thousands of taxa across species evolution^40,41^, and with the added challenge of having to sequence the ∼3 billion bases in the human genome, instead of hundreds to thousands of phylogenetically-informative loci, in order to obtain dense SNV variation. While for some phylogenetic studies, a topological constraint or divide-and-conquer strategy can be applied based on *a priori* knowledge about species placement in major taxa, we in contrast lack information about taxa partition. As such, we innovated optimized approaches for efficient tree building that would be suitable for application to this large dataset at feasible cost and analysis time.

First, while most previous scWGS studies called SNVs independently in each single-cell genome^22,23^, to increase sensitivity of identifying variants shared between cells at sequencing coverage 15×, we combined a per-cell strategy with joint genotyping. Second, we leveraged software that classifies germline versus somatic variation without a matched bulk sample^42^, allowing us to reconstruct all cell relationships including early embryonic mutations. Next, we developed a strategy to account for single-cell sequencing errors. Standard-PTA+RNA introduces low-frequency nucleotide substitution errors during genomic amplification and is further affected by RNA-to-DNA contamination^29^, leading to RNA edits being miscalled as genomic variants. Although software to remove PTA errors has been developed^43^, this approach requires matched bulk sequencing and is not designed to remove RNA edits. Thus, in order to identify and filter these sources of noise in our analysis, we developed an *XGBoost* machine learning SMART-PTA-specific error classifier that models multiple features such as maximum variant allele frequency of the SNVs across cells or RNA expression levels (**Methods**). To obtain a set of true positive and false positive calls for training, we used linked-read information^44^ to accurately classify somatic SNVs by using read-level phasing with proximal germline heterozygous single-nucleotide polymorphisms (SNPs). We observed that our false positives used for training had trinucleotide context profiles highly similar to those seen in PTA-amplified single cells^43^, and we were able to generate features for over 37,000 sites, compared with fewer than 1,000 sites reported previously (**Supplementary Fig. 3A**). The classifier applied to all esophageal samples distinguished true positive from false positives with an AUC of 0.978 (**Supplementary Fig. 3B**), higher than the AUC achieved by the published PTA error classifier (0.79)^43^, which we speculate may be due to our significantly larger training set as well as additional features (**Supplementary Table 4**). Notably, after applying the classifier to our SMART-PTA data, we observed depletion of mutation signature SBS30 (previously linked to PTA error source^43^) (**Supplementary Fig. 3C**), indicating that our classifier effectively removes errors introduced by PTA.

In addition, we observed that before the classifier was applied, there was a significant difference in the mutation burden of lowly versus highly expressed genes, where highly expressed genes harbored a higher number of SNVs (**Supplementary Fig. 3D**), suggesting the presence of variant calls deriving from RNA edits. In contrast, after application of the classifier, there was no significant difference in the number of SNVs found in low-expression versus high-expression genes, suggesting that our classifier reduces RNA contamination of genotype calls.

We next built large single-cell phylogenies for each donor. Phylogenies from single-cell expansions have typically been reconstructed using parsimony on binary cell genotypes (mutation presence/absence)^45,46^. However, the allelic imbalance/dropout introduced during whole-genome amplification can severely distort the true genotype matrix. For this reason, we relied on a single-cell maximum likelihood model (*CellPhy*) which leverages diploid genotype likelihoods and has been shown to better accommodate for amplification error sources^47^. Indeed, direct comparison of trees built using *CellPhy* or parsimony (*PHYLIP mix*; https://phylipweb.github.io/phylip/doc/mix.html) revealed fewer instances of clonal split across cell types or esophageal regions consistent with higher accuracy of *CellPhy* versus parsimony on SMART-PTA data (**Supplementary Fig. 3E, 3F**). However, the compute time required for obtaining a maximum likelihood tree with 100 bootstrap replicates using *CellPhy* is more than 1 day for only 61 cells (eso01), rendering this approach infeasible for our large dataset comprising hundreds of cells. Since running times scale superlinearly with the number of cells, extrapolation suggests that analysis at this scale would require years of computation [*Inferring Phylogenies*, Joseph Felsenstein, 2003]. One way to mitigate this would be through parallelization, which historically in phylogenetic reconstruction efforts has been leveraged mostly through distributing computations across variant sites, rather than across taxa (because alignments often contain far more sites than taxa). However, in such analyses, multiple tree initializations and bootstrap replicates are typically executed sequentially, despite being independent and potentially parallelizable through Message Passing Interface (MPI). Given the size of our dataset, to allow for a tractable compute time, we developed *CellPhyCloud*, an asynchronous distribution of *CellPhy* that enables scalability across large computing clusters through MPI-independent parallelization across tree searches (**Supplementary Fig. 3G**; see Code Availability; **Methods**). Using *CellPhyCloud*, we were able to obtain a 608-cell phylogeny (eso02) with 100 bootstrap replicates in less than 3 days, which we estimate would have taken 3.5 years without parallelization.

Next, we optimized our strategy for time calibration. Single-cell phylogenies usually show branches that represent genetic distance in number of mutations^10,25^, but these can also be calibrated to represent time. Bayesian methods that have been developed to transform mutation burden into time^48,49^ do not scale well with increasing numbers of cells. For this reason, we leveraged *RelTime*, an approach that can transform the tree branch lengths in number of mutations into time in years in a few seconds by relaxing the assumption of a strict molecular clock, using one or more calibration constraints^50^. To obtain calibration points, we leveraged the known developmental timing of germ layer specification (∼2 weeks after fertilization), and assumed that immune and epithelial cells diverged prior to birth. Importantly, even when calibrating the root only to the donor’s age plus gestation, without providing any additional constraints, we observe nodes that generate both immune and epithelial descendants coalescing during embryonic development, as expected (**Supplementary Fig. 3H**). Relaxing the assumption of a molecular clock, also aids in accommodating differences in cell type evolutionary rates, as under the null hypothesis using *rtreefit*^49^ that assumes same evolutionary rates for all cells, developmental timings were not consistent with estimated divergence times. This suggests that *RelTime* can be appropriately applied in a somatic setting, beyond its use in the species evolution^51–53^.

Together, this pipeline allowed us to generate large, phenotypically annotated phylogenies whose depth (distance from root to terminal nodes) represents the donor’s lifespan, in order to reconstruct the somatic evolution of thousands of single cells, directly from primary donor samples (**Fig. 3A, Supplementary Fig. 4, Supplementary Fig. 5A, 5B, 5C**). To assess the robustness of phylogenies, we performed bootstrap analysis. Most of our nodes were well supported, with a median ± MAD bootstrap support of 90 ± 14.826. Although we observed very short branches with low support near the root of the tree, this is expected, as some early divisions may occur without any mutations, effectively creating a polytomy, and rare embryonic mutations may not be consistently captured by bootstrap analysis^10,25^. As expected, the number of clades detected at birth in each donor, n = 11 (eso01), n = 60 (eso02), n = 141 (eso03) and n = 191 (eso04) clades **(Fig. 3A, Supplementary Fig. 5A, 5B, 5C**) scaled with the number of cells sequenced.

**Figure 3:**
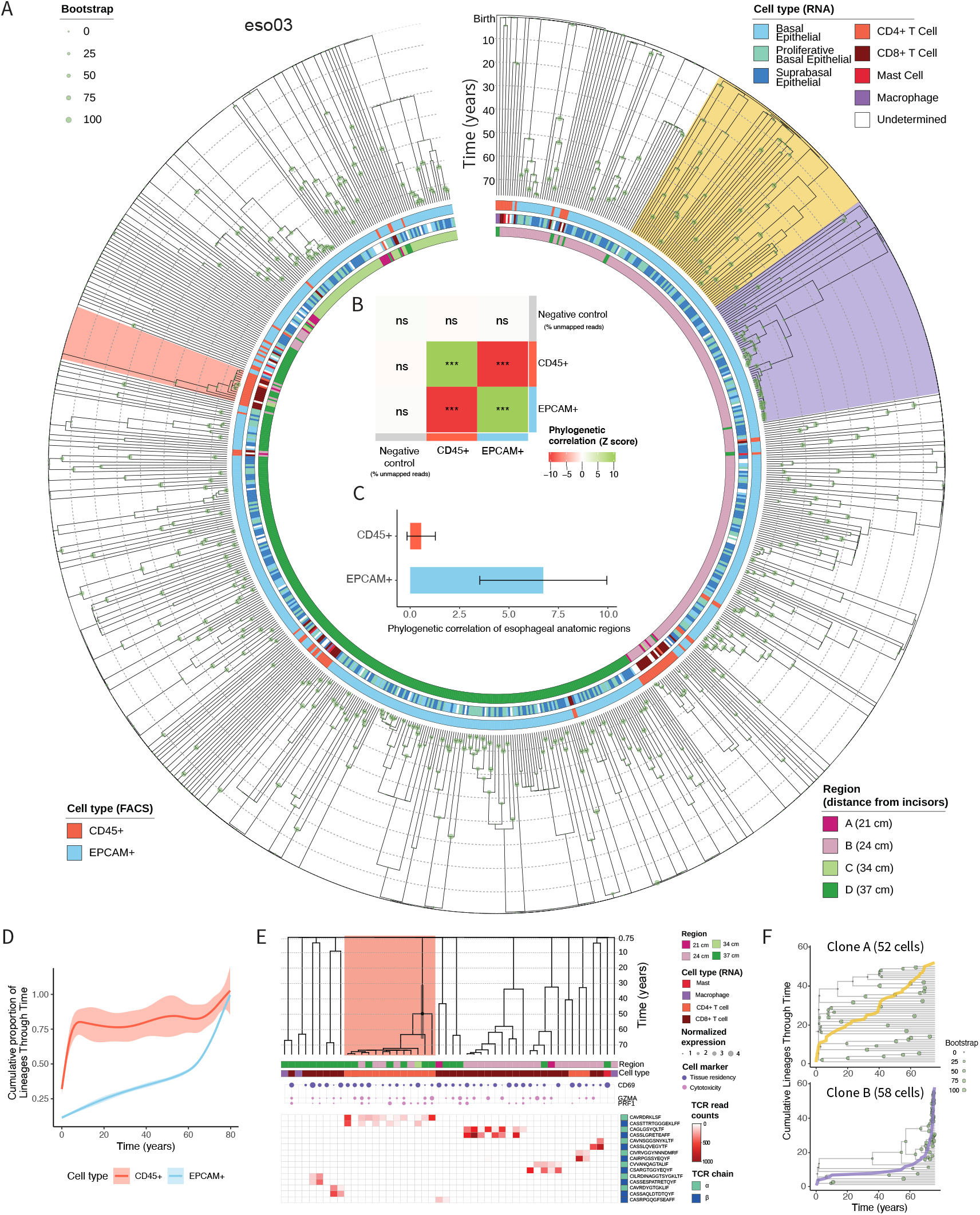
High-resolution single-cell phylogeny from esophagus tissue captures and quantifies human somatic evolution. **A**, Single-cell phylogeny of n = 758 total cells built from SNVs obtained from SMART-PTA application to esophagus samples from donor eso03. Each leaf of the tree represents a single cell, annotated as epithelial (EPCAM+) or immune (CD45+) cell based on flow cytometry data, as well as according to cell type determined from the RNA profiles. Regions corresponding to the sampling scheme in Fig. 2A are indicated and annotated for each cell. Branch lengths represent time in years. Bootstrapping values are indicated by green circles. Phylogenies for donors eso01, eso02 and eso04 are in Supplementary Fig. 5A-C. **B**, Phylogenetic correlation Z score heatmap of CD45+ (immune), EPCAM+ (epithelial) or negative control (percentage of mapped reads) cells for donor eso03. Positive correlation indicates that the cell type phenotype is shared by highly related cells. Negative correlation indicates that the cell type phenotype is absent from highly related cells. No correlation indicates that the distribution of a cell phenotype across highly related cells is random. *** One-sided *P* < 0.05. ns, not significant. Phylogenetic correlations were calculated using node distance to avoid circularity, as patristic distances were estimated using epithelial-immune divergence assumptions. Raw phylogenetic correlations are transformed into Z scores representing permutation tests using the *PATH* framework^19^. **C**, Mean phylogenetic correlation of esophageal regions for the EPCAM+ cells and CD45+ cells for donor eso03, indicating the degree to which phylogenetically related cells (immune versus epithelial cells) are dispersed in different regions of the esophagus. Phylogenetic correlation Z score of the distance of the biopsy from the incisors is calculated for EPCAM+ or CD45+ cells, respectively. Raw phylogenetic correlations are transformed into Z scores representing permutation tests using the *PATH* framework^19^. Error bars represent the standard deviation of 5 subtree replicates (Methods). **D**, Generalized additive model (GAM) fit of the cumulative proportion of lineages through time (years) across non-overlapping clones of exclusively epithelial and immune cells at birth across donors eso02, eso03 and eso04. The shaded area represents the 95% confidence interval. **E**, Analysis of immune cell clones (with ≥ 2 cells per clone) in donor eso03. The phylogeny is annotated with immune cell type assignments and anatomical region from which the cells were sampled (top). Expression of marker genes of tissue residency and cytotoxicity (circle size indicates expression level) is shown (middle). Complementarity-Determining Region 3 (CDR3) amino acid sequences from α or β TCR chains are shown (bottom). Only TCR sequences present in at least two sister cells are shown. CD4+ and CD8+ T cell types were manually assigned based on *CD4, CD8A*, and *CD8B* gene expression levels. A CD4+ clone expressing cytotoxicity and exhaustion markers is highlighted in pink. **F**, Cumulative number of lineages over donor age (lineage through time, LTT) for two clades highlighted on the phylogeny in A comprising epithelial cells with distinct topology on the tree. Top, clone A, n = 52 cells, yellow. Bottom, clone B, n = 58 cells, purple.

Finally, we annotated each individual cell on the phylogeny with phenotypic information, including esophagus sampling region, classification as epithelial or immune cells from index sorting and cell type assignments from RNA expression data (**Fig. 3A, Supplementary Fig. 4, Supplementary Fig. 5A, 5B, 5C**). To facilitate broader accessibility for the wider community, phylogenies are publicly available at https://itol.embl.de/shared/2CNE84KS0anV4, where they can be displayed next to multiple layers of phenotypic annotations for facile exploration.

### Phylogenetics of the aging human esophagus

We examined the distribution of epithelial and immune cells across the phylogenies to better understand their evolutionary patterns. Notably, we did not observe mutations uniquely defining two separate clades composed exclusively of epithelial or immune cells. Instead, we confidently identified early postzygotic mutations that are shared between exclusive sets of epithelial and immune cells (**Supplementary Fig. 5D**). These mutations likely arose before gastrulation, during any cell division after fertilization and before the second week of development, as further supported by their presence in multiple esophageal regions. This lack of strict segregation between immune and epithelial cell types is consistent with the non-monophyletic origin of germ layers, wherein certain endoderm-derived epithelial clones may share a more recent common ancestor with mesoderm-derived immune clones than with other endodermal cells^25,54,55^, such that early developmental diversification events can result in intermingled lineage relationships across distinct cell types. Despite this, the distribution of immune and epithelial cells on the tree was not random. To quantify the distribution pattern and assess how clustered or interspersed immune cells were in relation to epithelial cells, we determined phylogenetic correlations, which calculate how strongly a given cell state aligns with the phylogenetic tree structure. For this analysis, we used the Phylogenetic Analysis of Trait Heritability (*PATH*) framework^19^ that quantifies the degree of phenotypic similarity between closely related single cells on a phylogeny. We observed a strong negative cross-correlation between epithelial and immune cells (**Fig. 3B, Supplementary Fig. 5E**), indicating that overall, the sampled epithelial cells are more closely related to other epithelial cells compared to sampled immune cells, and *vice versa*. Thus, while we did not observe absolute separation between these populations, the patterns were consistent with epithelial cells differentiating from a pool of stem cells restricted to that lineage, leading to expansions of related epithelial or related immune lineages.

Our study design, which involved multi-regional sampling from distinct locations along the esophagus, allowed us to quantify the degree of cell migration for the epithelial and immune cells types. Briefly, by calculating phylogenetic correlations over anatomical regions (distance of the biopsy from the incisors), using *PATH*^19^, we estimated the degree to which spatial positioning was shared among closely related cells, for epithelial and immune populations. Here, high phylogenetic correlation would indicate that the anatomically co-localized cells are closely related, reflecting limited cell migration, whereas low phylogenetic correlation would suggest that cells within the same region are less related, consistent with higher migration. We found that location had high phylogenetic correlation in epithelial cells, while it was much lower in immune cells (**Fig. 3C, Supplementary Fig. 5F**). These results are consistent with both ontogeny and cellular niche of the distinct cell types. Regarding ontogeny, immune cells present in the esophagus travel from the yolk sac during embryonic development^56^, from the liver at fetus stage or from the bone marrow after birth^57–59^, while epithelial cells arise locally from basal progenitors within the tissue^37,60^. With respect to the niche, adherent epithelial cells reside within pits delineated by connective tissue papillae^61^, potentially limiting their spread along the lining, while motile immune cells can migrate locally within their niche in response to inflammatory signals^57,62,63^. Previous studies estimated high spatial clustering of esophageal epithelial clones with shared mutations in aged esophagus samples, with a reported median range of 2.87 mm^2^ separating related clones^4^, which is smaller than the distance between our biopsied regions. This is also consistent with the model of clonal constraints arising between neighboring mutant cells in the normal esophagus^64^.

We next performed a more detailed analysis of the clonal growth patterns. To first assess temporally-resolved clonal dynamics, we analyzed immune and epithelial cell populations across all the donors profiled with SMART-PTA, by calculating the cumulative proportion of lineages over donor age. Performing lineage-through-time (LTT) analysis^65^, we reconstructed cell-type–specific population trajectories. The scale enabled by SMART-PTA is particularly advantageous for this type of phylodynamic analysis, as LTT accuracy improves with larger numbers of sampled lineages^66^. Immune cells displayed a sharp increase in lineage diversity early in life followed by a plateau (**Fig. 3D**), reflecting the early exponential expansion that establishes the hematopoietic stem cell pool which is further maintained throughout aging^11^. We also observed a pattern consistent with secondary rise in older age in eso03 (**Supplementary Fig. 5G**). This donor had salient immune cell expansions encompassing tissue-resident lymphocytes (red clades, **Supplementary Fig. 4C**), which could potentially relate to inflammation or infection later in life^34,67^. In contrast, epithelial cells showed a slower, steady accumulation of lineages, with a notable inflection around age ∼60 (**Fig. 3D**; **Supplementary Fig. 5G**), reflecting emergence of somatic mutant clones in the esophageal epithelium that accumulate with age, particularly clones with selected mutations in cancer-associated driver genes that become more prominent after age ∼50-60^3,4^.

We next specifically focused on immune clones. To independently validate clonal expansion, we took advantage of single-cell full-length transcriptomes and analyzed T cell receptor (TCR) transcript sequences, commonly used as a molecular barcode for lineage tracing of T cells^68,69^. Consistent with our phylogenies, we identified the same full-length TCR α and β chains only among cells within expanded T cell clones (**Fig. 3E**, bottom), supporting the robustness of WGS-based clonal assignment. While previous studies have investigated immune cell phylogenies from blood precursors sampled from bone marrow^10,11^, to our knowledge, whole-genome phylogenies derived from differentiated non-malignant immune cells in solid tissues remain unexplored. We selected all immune clones containing at least two cells with RNA-defined immune cell assignment from donor eso03, who showed evidence of immune cell expansions (red clades, **Supplementary Figure 4C**), and annotated the phylogeny with anatomical region, cell type and expression level of different marker genes (**Fig. 3E**). Notably, several immune clones showed high degree of spatial dispersion, spanning all sampled anatomical regions of the esophagus, consistent with the migratory nature of T cells contributing to these expansions. The expression of *CD69* confirmed that the sampled cells are tissue resident rather than circulating lymphocytes^70^ (**Fig. 3E**, middle), consistent with the avascular nature of the esophageal epithelium from which they were extracted. Aligning with previous observations^71^, the vast majority of the clones were tissue-resident CD8+ T cells. However, we also observed some CD4+ T cell clones, and other immune cell subtypes such as macrophages or mast cells. We observed that the largest CD4+ T cell clone (orange highlight, **Fig. 3E**) expressed markers of cytotoxicity (*GZMA, PRF1*), a phenotype which is not commonly associated with this cell subtype, but that has been shown to increase during aging^72^. This CD4+ T cell clone showed a positive γ-statistic of 1.97 (*P* < 0.05), which tests for deviation from constant rate diversification, suggesting that its proliferation is not the result of physiological tissue turnover, but likely reflects clonal expansion, perhaps driven by antigen exposure. Because the γ-statistic is sensitive to incomplete sampling, we also applied the Monte Carlo Constant Rates (MCCR) test^66^, confirming that the dynamics are not caused by incomplete sampling and obtaining almost the exact same γ-statistic and a significant *P* value (*P* < 0.01).

To gain further insight into the clonal dynamics of epithelial cells, we next assessed epithelial clades on the phylogeny. A visual evaluation of the complex tree topology suggested the presence of distinct clades exhibiting different branching patterns over the donor lifespan (e.g., eso03, **Fig. 3A**). To better define these patterns, we selected the two largest epithelial clones, clone A (n = 52 cells; **Fig. 3A**, yellow) and clone B (n = 58 cells; **Fig. 3A**, purple), and performed LTT analysis^65^, allowing us to track the accumulation of lineages from the inferred time of origin (fertilization) to the donor’s age at sampling (e.g., 76 years for donor eso03). The resulting patterns revealed striking differences between clones: clone A showed a steady, linear accumulation of epithelial lineages over time, whereas clone B exhibited a delayed, rapid expansion in lineage number, especially in the donor’s final decade (**Fig. 3F**). These divergent trajectories were further supported by the γ-statistic analysis. Clone B showed a significantly positive γ-statistic (γ = 6.43, *P* = 1.3 × 10^-10^), indicating an excess of recent branching events, potentially reflecting the late acquisition of multiple somatic driver mutations in this clone (see below). In contrast, clone A exhibited a significant negative γ-statistic (γ = –3.92, *P* = 8.8 × 10^−5^, MCCR *P* < 0.01), consistent with a slowdown in diversification over time that potentially reflects a selective disadvantage of the cells in this clone compared to neighboring cells^64^. These analyses suggest that clones followed distinct evolutionary dynamics that deviate from the neutral expectation. To further confirm different evolutionary patterns exhibited by these clones, we tested whether the two clones diversify under the same birth-death process (null hypothesis) or under distinct diversification regimes (alternative hypothesis) considering 0.5% sampling (**Methods**) by fitting separate birth-death models to each clade, and comparing them to a joint model with shared parameters. This analysis strongly supported the alternative hypothesis (likelihood ratio test (LRT) = 87.76, *P* = 8.8 × 10^-20^), indicating that the two clones diversified under significantly different birth-death parameters. These behaviors reinforce the idea that somatic clones experience fundamentally distinct evolutionary trajectories driven by different mutation histories. Indeed, we speculate that the pronounced growth later in life observed in clone B coincided with the acquisition of multiple driver mutations, which we explore in the following section.

### Analysis of mutational signatures over time reveals exposure history in the aging esophagus

Motivated by our clonal dynamics analysis across clades, we next focused on characterizing the somatic clonal evolution of the esophageal epithelium, integrating the genetic, phylogenetic and transcriptomic data to assess mutant clones and their phenotypes (**Fig. 4A, 4B, Supplementary Fig. 6A, 6B**). We hypothesized that differential exposure to environmental risk factors would affect mutation patterns across donors. We stratified donors into low-, medium- and high-risk categories based on self-reported history of tobacco and alcohol consumption (**Supplementary Table 1**). These exposures are well-established risk factors for esophageal diseases, including cancer^73^. Notably, previous studies reported that heavy alcohol drinking and tobacco smoking accelerate clonal remodeling^4^ and significantly increased mutation burdens in the phenotypically normal esophagus^3^. Based on exposure history, donors eso01 and eso02 (former smokers with regular alcohol consumption) were classified as high-risk, eso03 (alcohol consumption) as medium-risk, and eso04 (no history of tobacco or alcohol use) as low-risk. To evaluate the effects of environmental exposures on global mutational processes in the human esophageal epithelium, we leveraged the extensive SNV information obtained with SMART-PTA to analyze single-base substitution (SBS) trinucleotide mutation signatures for all mutations on the phylogeny, including coding and non-coding mutations. Specifically, we examined the trinucleotide mutational context of SNVs along branches of clades for high-risk donors (eso01, eso02), medium-risk donor (eso03) and low-risk donor (eso04). This analysis revealed five SBS signatures across the four donors, including three ubiquitous/clock-like signatures (SBS1, SBS5 and SBS40a) and two signatures associated either with tobacco exposure (SBS4) or aldehyde exposure related to tobacco use/alcohol consumption (SBS16)^74,75^ (**Fig. 4C, Supplementary Fig. 6C, 6D**).

**Figure 4:**
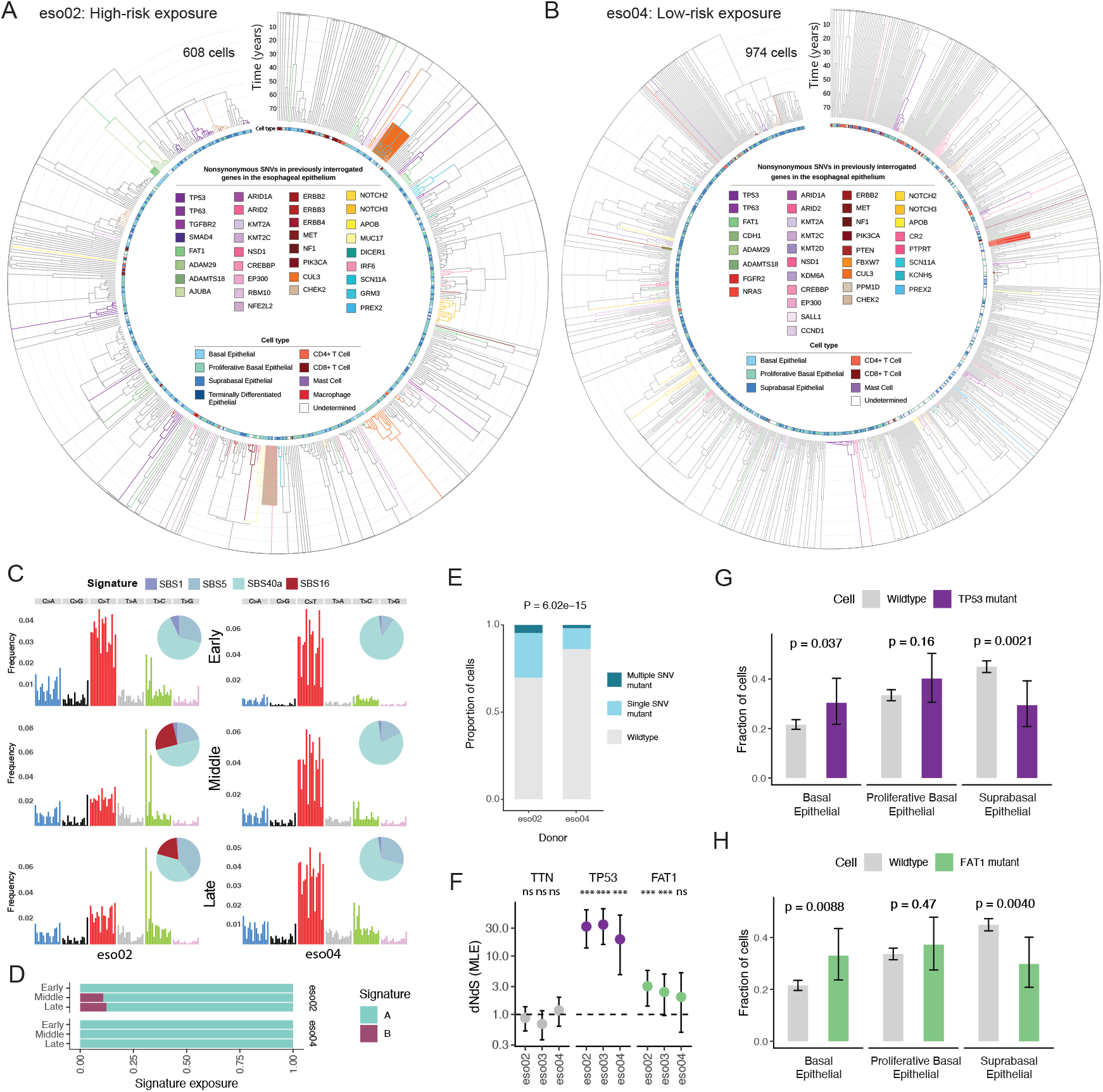
Driver mutations and signatures in aging human esophagus from donors with high-risk versus low-risk exposures. **A**, Single-cell phylogeny of n = 608 cells built from SNVs obtained from SMART-PTA application to esophagus samples from a 79-year-old donor eso02 with history of smoking and alcohol use (high-risk exposure). Each leaf of the tree represents a single cell, with cell type annotations displayed. Clones harboring nonsynonymous mutations in any of 83 genes (Supplementary Table 5) previously analyzed in aging esophagus tissue^3,4^ are indicated in different coloring of the branches. Branches with two mapped mutations are further shaded with the gene color of the second mutation. Any subsequent mutation is further indicated on the relevant subclonal branch of the double mutant clone with the corresponding gene color. **B**, Single-cell phylogeny of n = 974 cells built from SNVs obtained from SMART-PTA application to esophagus samples from a 77-year-old donor eso04 with no history of smoking or alcohol use (low-risk exposure). Annotations are the same as for (A). **C**, Analysis of contributions of trinucleotide mutation signatures across age for eso02 (left) and eso04 (right) across lifespan determined from phylogenetic branches (Early – including branches whose end node is older than 40 years, covering a time frame from embryonic development to adulthood; Middle – including branches where the end node is less than 40 years old but terminates in an internal node; Late - terminal branches whose end node is the age of the donor; Methods; Supplementary Fig. 6E). Single-base substitution (SBS) signatures were obtained by analyzing the single-cell whole genome amplification (scWGA) data and aligning to COSMIC signatures. Trinucleotide spectrum and inset with percentage of contribution of four signatures (SBS1, SBS5, SBS16 and SBS40a) are shown for each donor across the three time points. Trinucleotide mutation signature analysis for donors eso01 and eso03 are presented in Supplementary Fig. 6C-D. **D**, Contribution of de novo signatures identified from whole-genome sequencing of oral epithelium samples from donors with a history of heavy smoking/heavy drinking and donors without smoking/drinking history^76^ in eso02 and eso04 at the same time periods as in (C). The A signature shares similarity with SBS1/SBS5 (clock-like) and the B signature shares similarity with SBS16 (aldehyde exposure related to tobacco use/alcohol consumption). **E**, Proportion of mutant cells in eso02 (high-risk exposure) and eso04 (low-risk exposure) donors with somatic variants in 28 esophageal driver genes (Supplementary Table 5) previously reported to show positive selection in esophagus tissue through dN/dS analysis^3,4^, including single-gene mutants, multi-gene mutants or wild type cells. Two-sided Fisher’s exact test. Proportions of mutant cells across all donors are presented in Supplementary Fig. 6J. **F**, dN/dS maximum likelihood estimation (MLE) analysis for *TTN, TP53* and *FAT1* genes in donors eso02, eso03 and eso04. The ratio of nonsynonymous to synonymous somatic variants was determined at the gene level (Methods). Error bars represent the 95 confidence intervals. Grey dashed line represents dN/dS of 1, indicating no selection. Values >1 indicate positive selection. *** *P* value < 0.001; ns, not significant, dNdScv Likelihood-Ratio Test per donor^89^. *P* values from left to right: *TTN*-9.6x10^-2^, 3.3x10^-1^, 6.8x10^-1^; *TP53*-2.2x10^-9^, 1.1x10^-9^, 1.2x10^-4^; *FAT1*-2.3x10^-3^, 1.01x10^-3^, 1.9x10^-1^. We randomly downsampled 608 cells from each donor to avoid dNdS biases due to clonal frequency differences^90^. The low number of mutations in eso01 (55 cells) prevented calculation of dNdS in this donor. **G**, Mean fraction of cells constituting each epithelial cell type (basal, proliferative basal and suprabasal) for cells without any detected mutation in *TP53* (grey) and *TP53*-mutant cells (purple) across donors. Error bars represent 95% Clopper-Pearson confidence intervals and *P* values were calculated using a one-sided χ^2^ test. **H**, Same as (G) for *FAT1* wild-type (grey) and *FAT1*-mutant (green) cells.

We further assessed how age and exposure jointly shape mutation accumulation and signature composition over time. For this analysis, we subdivided each phylogeny into three temporal categories based on the age of the branch’s end node: early (internal branches, embryonic to 40 years prior to sampling), middle (internal branches, 40 years prior to sampling up to the latest internal node), and late (external branches, representing the most recent clonal lineages, ∼ last decade; **Supplementary Fig. 6E**). Notably, SBS16 was observed later in life exclusively in the donors with history of alcohol consumption, (eso01, eso02 and eso03), whereas it was absent in the low-risk donor (eso04; **Fig. 4C, Supplementary Fig. 6C, 6D**). In contrast, the ubiquitous clocklike signatures (SBS1, SBS5 and SBS40a) exhibited relatively constant presence across all phylogenetic time intervals, consistent with their known association with endogenous, time-dependent mutational processes. Interestingly, the SBS4 signature associated with tobacco was observed from 40 years to the most recent internal node in donor eso01, but not in the latest period, (**Supplementary Fig. 6D, 6E**), consistent with this donor’s heavy smoking history and subsequent cessation within the last 10 years (**Supplementary Table 1**).

In addition, we evaluated the presence of mutational signatures previously identified from oral epithelium whole-genome sequencing studies^76^, including signature A related to SBS1/SBS5 and signature B, which shares similarity to SBS16 and was enriched in individuals with a history of heavy tobacco and alcohol use. We saw the presence of the exposure-related signature B at later time points in donor eso02 (**Fig. 4D**), consistent with our analysis of COSMIC SBS signatures. Importantly, these insights could only be uncovered by combining high-resolution phylogenetic information to estimate the time of lineages with high-depth scWGS to accurately identify mutation signatures, enabling analysis of the effects of environmental exposures on esophageal somatic evolution across lifespans. Such a framework could be used in the future to explore mutagenic footprints throughout an individual’s age, for insights into the dynamic effects of the collective exposome^77^ on the somatic genome.

### Defining somatic evolution of the aging esophagus through analysis of clones with driver gene mutations

We next focused on evaluating potential driver mutations in functionally important genes. Previous studies analyzing somatic variants in normal esophagus tissue of elderly donors, through bulk sequencing of tissue microdissections, have identified recurrently mutated genes, including *NOTCH1, TP53*, and *FAT1*^3,4^. To assess the clonal architecture of our aged esophagus samples, we profiled SNVs occurring in 83 genes previously interrogated in the esophageal epithelium^3,4^. We identified nonsynonymous somatic variants in 61 out of the 83 genes, and annotated the phylogeny with the identified mutant genes, observing both single- and multi-mutant clones (**Fig. 4A, 4B, Supplementary Fig. 6A, 6B, Supplementary Table 5, Supplementary Table 6**). Although most genes were well covered (mean depth >10× per cell), we note that *NOTCH1*, previously reported as the most frequently mutated driver gene in the phenotypically normal esophagus, had lower coverage (2.2×) (**Supplementary Table 5, Supplementary Fig. 6F**). This is likely due to increased GC content hindering single-cell DNA amplification^78^, as *NOTCH1* has over 63% GC content, compared to 49% for *TP53*, 42% for *FAT1*, and 41% genome-wide. We observed variable coverage across the *NOTCH1* gene, where regions with high GC content were poorly covered (**Supplementary Fig. 6G**), contrasting with genes that have lower GC content, such as *NOTCH2* (40% GC content), which had high coverage (17×; **Supplementary Fig. 6H**). To examine whether low coverage of *NOTCH1* coding region is due to insufficient sequencing or GC biases in UG sequencing, we designed a targeted panel for hybridization capture across exonic regions of seven driver genes, including *NOTCH1, FAT1* and *TP53*, and applied it to 48/61 libraries from donor eso01 (**Methods, Supplementary Table 7**). After Illumina sequencing the enriched libraries to saturation (79.55 ± 7.96 % duplicate rate along targets), we observed that coverage remained sparse (**Supplementary Fig. 6I**), with an average of 25.9% of exonic bases per cell not covered on *NOTCH1*. This confirmed that the key issue is at the single-cell gDNA amplification step.

Twenty-eight genes were previously identified to be under positive selection in the normal esophagus^3,4^ (hereafter referred to as esophageal driver genes, **Supplementary Table 5**). Three of the twenty-eight genes (*NOTCH1, PLXNB2*, and *ZFP36L2*) suffered from low coverage (median read coverage per cell over 500-kb bins < 4; **Supplementary Table 5**) that likely hindered variant calling. We quantified the prevalence of somatic variants (both synonymous and nonsynonymous) within these 28 genes (single-gene mutants or multi-gene mutants) across donors, observing that high-risk donors had overall higher proportions of mutant cells compared to medium- and low-risk donors (**Fig. 4E**; **Supplementary Fig. 6J**). These results indicate that donors with higher exposure to risk factors have increased mutation burdens in esophageal driver genes^79,80^.

We next assessed the frequency of mutations in individual genes, analyzing putative high-impact variants (nonsynonymous SNVs with CADD PHRED score ≥ 20) in the 28 esophageal driver genes. Of these, *TP53* and *FAT1* were the most frequently mutated esophageal driver genes across patients (**Supplementary Table 5**), in line with mutation frequencies previously observed in whole-exome sequencing data from esophagus samples from aging donors, where these genes were observed among the top recurrently mutated esophageal driver genes^4^. We identified multi-mutant clones with more than one esophageal driver gene mutation, including those harboring mutations in different driver genes (e.g., *TP53* and *FAT1*, **Supplementary Fig. 6K**, donor eso02 cells). We also identified a double-mutant clone with distinct mutations in *FAT1*, a tumor suppressor gene in the HIPPO pathway^81^ involved in cell-adhesion^82^ and genome integrity^83^ that is recurrently mutated in the normal esophagus^3,4,84^. Interestingly, descendants of this *FAT1p*.*E4130X* ancestral clone acquired a total of three mutations affecting the key cell adhesion genes *FAT1* and *AJUBA*^85–87^ (**Supplementary Fig. 6L**).

To evaluate the intensity of selective pressure acting on the esophageal driver genes, we performed a dN/dS ratio analysis, calculating the ratio of nonsynonymous to synonymous somatic variants at the gene level (**Methods**). This analysis is used to determine whether a gene is under positive selection (ratio greater than 1), negative selection (ratio less than 1) or neutral drift/no selection (ratio of 1). We calculated dN/dS values for the most frequently mutated genes, *TP53* and *FAT1*, in donor eso02, eso03 and eso04. As a negative control, we calculated dN/dS ratio for *TTN*, which accumulates a large number of somatic variants due to its size, but should not be under positive or negative selection in the esophageal epithelium. Our analysis revealed that *TP53* and *FAT1* showed dN/dS values higher than 1, suggesting that a positive selection process is acting on clones mutated for these genes (**Fig. 4F**), consistent with results from previous bulk sequencing studies performed at the cohort level^3^. This per-patient dN/dS analysis, enabled by the high throughput of SMART-PTA, demonstrates the feasibility of detecting candidate genes under selection within a single donor. Although the number of mutations is not always sufficient to reject neutrality, we observed that the strength of positive selection on *TP53* and *FAT1* is broadly comparable between the donors regardless of risk level. Combined with our results showing that high-exposure donors have an overall higher mutation burden in selected driver genes (**Fig. 4E**; **Supplementary Fig. 6J**), these findings suggest that while exposures increase mutation rates, it is possible that selection might operate similarly on both high/low exposure samples, likely reflecting tissue-specific selection pressures that act during physiological aging, although this observation requires future exploration across a larger number of individuals.

We then analyzed the matched single-cell transcriptome data in order to obtain functional insights into the effect of these esophageal driver gene mutations on cell phenotypes, in particular for the more frequent clones harboring *TP53* and *FAT1* mutations. We evaluated the distribution of *TP53*-mutant and *FAT1*-mutant cells across basal, proliferative basal and suprabasal epithelial states (**Fig. 2C, Supplementary Fig. 2H**). We observed a significant enrichment of *TP53*-mutant cells in the basal epithelium state, accompanied by a reciprocal depletion in the fraction of *TP53*-mutant cells that were in the more differentiated suprabasal compartment (**Fig. 4G**). Previous studies using mouse models have reported that p53-mutant progenitors are biased towards basal cell differentiation^88^. Our results suggest that *TP53* mutation may bias cells towards an earlier differentiation state in humans, potentially facilitating clonal selection by expanding the pool of self-renewing basal cells. Similarly, we observed a significant decrease in the fraction of *FAT1*-mutant cells that were in the later suprabasal state compared to wild-type *FAT1* cells, and a corresponding increase of *FAT1*-mutant cells in the earlier basal state (**Fig. 4H**), suggesting that *FAT1* mutation could also be driving cells towards an earlier differentiation state. These genotype-to-phenotype mapping results reveal that *FAT1* mutations can be selected for in the phenotypically normal esophagus due to biased differentiation that skews towards the earlier basal state. This underscores the importance of this mechanism in shaping the somatic evolution of the esophagus, where mutations in multiple driver genes promote the same earlier cell state phenotype.

To orthogonally validate these findings, we reanalyzed data from an alternative single-cell genotype-to-phenotype method, scG2P^84^. This approach uses multiplexed targeted DNA and RNA sequencing, where RNA marker genes are used to calculate the differentiation score (**Methods**). These data supported our findings, as differentiation scores for *TP53*-mutant and *FAT1*-mutant cells were significantly lower than for wild-type cells (**Supplementary Fig. 6M**), indicating that the mutant cells are biased towards earlier differentiation states. These results provide further support for the phenotypic effect of somatic mutations within these esophageal driver genes on cell differentiation state.

### Clonal landscape of copy-number variants and identification aneuploidy events with driver SNVs in the same clone in the aged human esophagus

Unlike malignancies, normal tissues are considered to be largely diploid with limited numbers of structural variants or copy-number changes, which can represent the key genetic difference between benign and cancerous cells^91–93^. However, recent emerging data in normal breast tissue uncovered the presence of a low level (∼3%) of aneuploid cells^14^, promoting a rethinking of genome integrity as a specific hallmark of cancer. Having characterized the landscape of somatic SNVs in aging esophageal epithelium, we next investigated large-scale structural alterations by inferring the copy-number variations (CNV) across each individual single cell to assess levels of chromosome instability in this non-malignant solid tissue. Across all four donors, the vast majority of cells exhibited a euploid genome with no detectable copy number alterations (**Fig. 5A**), consistent with the non-malignant nature of the tissue. However, even after applying stringent filters to only retain cells with the highest coverage uniformity (1,587 /2,783 cells retained), we detected a subset of epithelial cells showing clear aneuploidy present in at least two sister cells [18/1,490 (1.21%)]. This likely represents a conservative estimate, broadly similar to findings in normal breast tissue, where 3.19% of single breast epithelial cells were reported to be aneuploid^14^. In contrast, immune cells did not show evidence of autosomal aneuploidy. The only prevalent aneuploidy event that we observed in the immune cells was chromosome X loss in female donors (**Fig. 5B**), consistent with mosaic loss of the X chromosome being the most frequent clonal somatic event reported in female leukocytes^94,95^. Notably, the presence of phylogenetic clades of more than two aneuploid cells indicates that these cells can bypass cell-cycle surveillance mechanisms and continue replicating despite harboring large CNVs. Aneuploidy events in esophagus epithelial cells included a clone with amplification of chromosome 3q and a clone with chromosome 6 amplification in donor eso03, as well as a clone with chromosome 12 amplification in eso04 (**Fig. 5A**). In addition, we identified a clone in eso03 that displayed aneuploidy across multiple chromosomes (including amplifications of chr3q, chr5, chr6, chr8, chr17 and chr21; **Fig. 5A**). Notably, recurrent whole-chromosome 3 amplifications impacting driver genes *PIK3CA, SOX2* and *TP63* are observed in ∼50% of ESCCs^96^. More infrequently, chr3q amplifications have been identified at the sample level through bulk analysis in the normal esophagus^3,4^. We orthogonally validated the presence of aneuploidy in these four clones by analyzing germline heterozygous SNPs, in order to assess allelic imbalance (**Methods**). Indeed, we observed allelic imbalances for identified aneuploidy events (**Fig. 5C**), suggesting that the chr3q amplification with an estimated copy-number of 4 in donor eso02 could represent a triplication of one allele (e.g., AAA+B gain) instead of duplication impacting both copies (AABB), where A and B represent distinct parental alleles.

**Figure 5:**
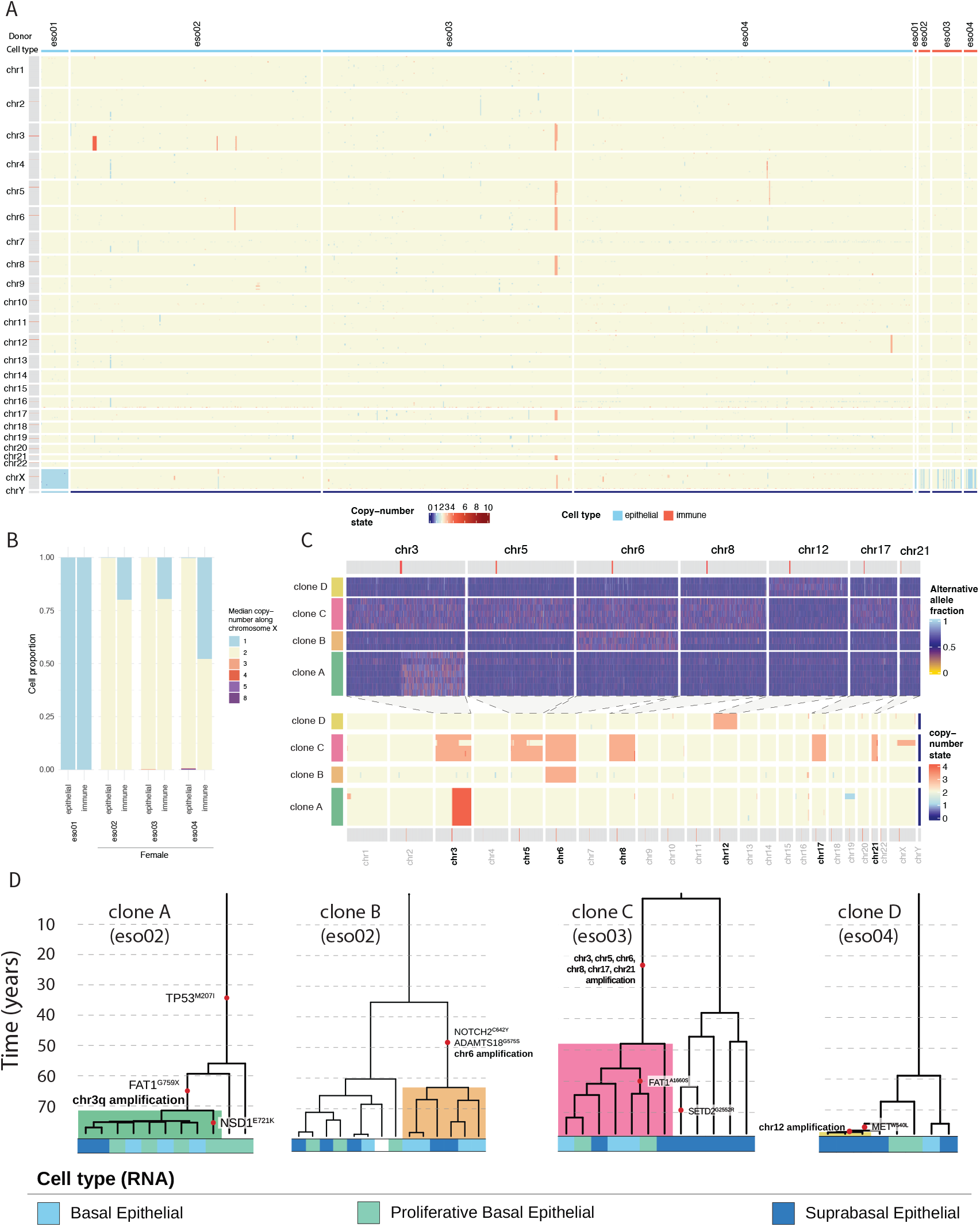
Copy-number variations and aneuploidy events in the aging human esophagus. **A**, Genome-wide copy-number profiles for epithelial and immune cells across all donors. Copy-number state is displayed for all chromosomes along ∼500 kb non-overlapping windows. **B**, Median X chromosome copy number in proportions of epithelial and immune cells for each of the four donors. **C**, Top, mean phased alternative allele frequency for the indicated chromosome regions for four clones identified with CNV events across 3 donors, with matching copy-number state (bottom). **D**, Example zoomed-in trees of clones bearing both driver SNVs as well as CNV events. Dashed lines added at 10-year intervals. Driver variants and CNV events are labeled, and cell types are indicated, for two clones in donor eso02 and one clone each for donors eso03 and eso04. Shaded colors correspond to the clone labels (clone A-clone D) in (C). We note that in the initial CNV analysis (panel A), we only included 1,587/2,783 cells with MAD score lower than 0.20, and detected 18 epithelial cells with CNVs supported by at least two sister cells. Remapping these cells onto the tree allowed for more careful assessment of CNVs in neighboring cells with lower stringency on the coverage quality. This analysis revealed 25 cells that we could annotate on the phylogeny.

To investigate whether loss of function in esophageal driver genes contributes to the emergence or selection of aneuploid clones, we intersected these aneuploidy events with driver SNV annotations. While shallow scWGS technologies have previously linked somatic SNVs and CNVs through pseudobulking approaches^13^, SMART-PTA achieves this directly with high-depth whole-genome coverage at the single-cell level, eliminating the need for aggregation of hundreds of cells (at ∼0.05× median coverage^13^). This analysis revealed that several aneuploid epithelial clones also harbored mutations in known esophageal driver genes, suggesting potential cooperation between point mutations and chromosomal alterations in early clonal expansion (**Fig. 5D**). Specifically, clone A from donor eso02 with a chr3q amplification also carried variants in two driver genes, *TP53* and *FAT1*, where the *TP53* variant was acquired prior to the CNV (clone A, **Fig. 5D**). On the other hand, clone C, which harbored multiple chromosomal amplifications events suggestive of genome instability, did not harbor a known driver co-occurring SNV (clone C, **Fig. 5D**). This could indicate that these rearrangements arise from compromised mitosis and non-disjunction events independently of driver gene point mutations.

### The *TP53*-mutant landscape in non-cancerous but high-risk exposed esophageal tissue

The identification of the *TP53*-mutant subclone that harbored a chr3q amplification event (**Fig. 5D**) prompted us to examine clonal *TP53* events in donor eso02, in which the proportion of cells carrying a *TP53* mutation was highest (>9%). This donor, with a documented history of genotoxic exposure (ex-smoker), but no clinical evidence of malignancy, represented a unique model for investigating mutational dynamics in pre-cancerous states. *TP53* is a critical tumor suppressor gene that is commonly mutated across a wide range of human neoplasias^97,98^, and has been reported as mutated in approximately ∼30% of the phenotypically normal esophagus^3,4,84^. Biallelic loss of *TP53* is a well-established driver of cancer development^99,100^, and prior studies have demonstrated that acquisition of a complete loss of *TP53* function through a “two-hit” mechanism is a critical prerequisite for malignant transformation of the esophagus^88,101^. Leveraging our clonally resolved dataset and the high coverage provided by SMART-PTA single-cell whole-genome sequencing, enabling biallelic capture, we performed a detailed characterization of the genomic events shaping the mutant *TP53* landscape, along with the phenotypic impact of these events, in the high-risk donor eso02. This approach allowed us to further expand our CNV analysis to consider CNLOH events, which can impact driver gene loci, including the *TP53* locus on chromosome 17p. Among the 614 epithelial cells from this donor, *TP53* represents the esophageal driver gene that is most frequently mutated (**Supplementary Table 5**), allowing us to investigate both the spectrum of biallelic and monoallelic events and their associated phenotypic consequences.

In addition to the chr3q amplification-bearing *TP53p. M246I* clone, we identified high impact variants of *TP53* (CADD PHRED score ≥ 20) in five multicellular clones, including one clone with a truncating variant (*TP53p. Q136X*), one clone with an in-frame deletion (*TP53p*.*R175_ H178del*) and three clones with missense variants (*TP53p. P151A, TP53p*.*C238Y* and *TP53p*.*G279V*) (**Fig. 6A, 6B**). All these variants are located in the DNA-binding domain of p53 (**Supplementary Fig. 7A, 7B, 7C**), and except *TP53p*.*M246I*, have been previously observed in phenotypically normal esophageal epithelium^3,4^.

**Figure 6:**
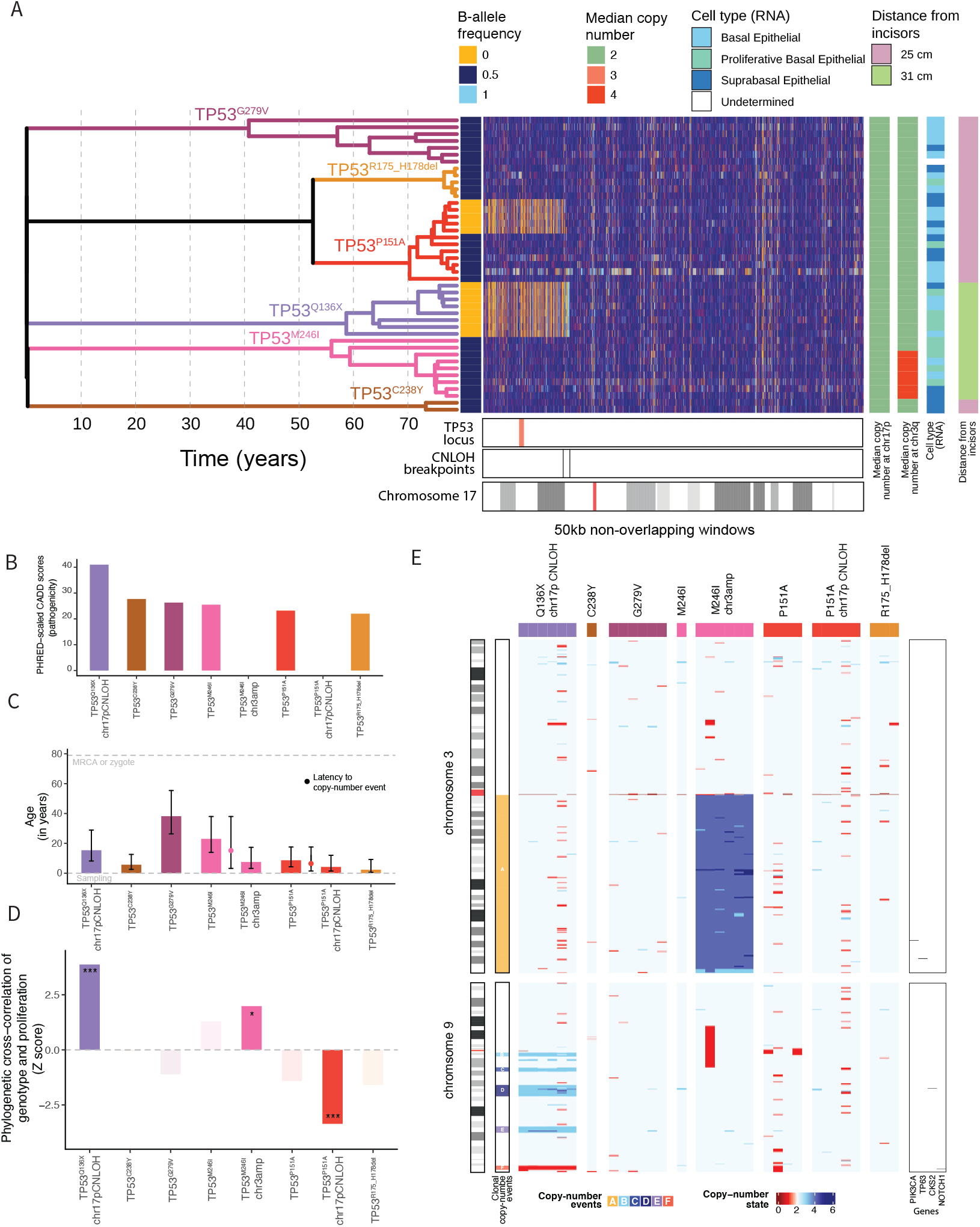
*TP53*-mutant clonal landscape of 79-year-old esophagus includes cells with monoallelic and biallelic inactivation of *TP53*. **A**, Tree displaying *TP53*-mutant clones from the donor eso02 phylogeny shown in Fig. 4a annotated with *TP53* SNVs (left) and chromosome 17p copy-number profile (right). Mean phased alternate allele frequency at the *TP53* locus, median copy number state at chromosome 17p and 3q, cell type assignments and esophagus region (biopsy distance from mouth; region B: 25 cm; region C: 31 cm) are displayed for each cell. The *TP53* locus and CNLOH breakpoints are indicated. Chromosome 17 bands are displayed with the centromere indicated in red. **B**, Combined Annotation Dependent Depletion (CADD) PHRED scores for the indicated *TP53* mutations identified in donor eso02. CADD score ≥ 20 indicates that the SNV is predicted to be in the top 1% of most deleterious out of all possible reference genome SNVs. Colors correspond to the variants on the phylogeny in (A). **C**, Estimated age of mutation event for the indicated eso02 *TP53*-mutant clones at the time of sampling. The latency from *TP53* point mutation to copy number event is indicated in red for the *TP53*.*pP151A* mutant clone, which acquired a later chromosome 17p CNLOH event, and in pink for the *TP53*.*pM246I* mutant clone, which acquired a later chromosome 3q amplification. Mutation age (in years) is estimated from the age of the ancestral node from which all descendant cells carry the mutation. Error bars represent the 95% confidence intervals. MRCA, most recent common ancestor. **D**, Phylogenetic correlation Z score of *TP53*-mutant cells with S phase score and G2M score (cell cycle score). *** = *P* value < 0.001; * = *P* value < 0.05. One-sided standard normal distribution. **E**, Copy-number variation heatmap of chromosome 3 (top) and chromosome 9 (bottom) for *TP53*-mutant cells from donor eso02. Six CNV events, one identified in a *TP53*.*pM246I* mutant subclone and five identified in the *TP53p*.*Q136X* mutant clone, are labeled A-F. Locations of the *PIK3CA, TP63, CKS2* and *NOTCH1* genes are indicated. Centromeres are indicated in red on each chromosome.

Data from mouse and organoid systems have also demonstrated that *TP53* heterozygous mutant cells in normal tissue evolve into cancer in a stereotypic manner, with subsequent *TP53* loss of heterozygosity (LOH) followed by accumulation of CNVs and aneuploidies^102,103^. We reasoned that we could leverage our single-cell lineage histories to trace the sequential acquisition of *TP53* loss events over the donor’s lifetime and assess chr17p CNLOH (**Supplementary Fig. 7D**). We identified two *TP53*-mutant clones (*TP53p*.*Q136X* and *TP53p*.*P151A*) harboring independent CNLOH events characterized by different breakpoints overlapping the *TP53* locus (**Fig. 6A**), resulting in a biallelic loss of function of the *TP53* gene. Then, we inferred the temporal origins of the *TP53* mutation events by assigning them the age (in years) of the most recent common ancestor of all the descendant cells that carried that mutation (**Fig. 6C**). Taking advantage of our ability to resolve clonal structure, we estimated that there was a multi-year latency time (∼5 years) from monoallelic to biallelic inactivation of *TP53* for the *TP53p*.*P151A*-mutant clone (**Fig. 6C**), which underwent a subsequent chr17p CNLOH event after SNV acquisition. Notably, in the other clone with biallelic *TP53* loss, the *TP53p*.*Q136X* and chr17p CNLOH events were found in all cells of the clone, and we estimated that cells existed with biallelic *TP53* loss for at least 15 years prior to sampling. These findings indicate that clones harboring complete functional loss of *TP53* can persist for ∼decades in histologically normal, non-malignant esophageal epithelium.

To gain insights into the phenotypic consequences of *TP53* mutations that we identified in eso02, we leveraged the tree to determine phylogenetic cross-correlations^19^ between *TP53* genotype and proliferation phenotypes (S phase score and G2M score) determined from the matched single-cell RNA data (**Fig. 6D**). For the proliferation phenotypes, the *TP53p*.*Q136X* mutation and *TP53p*.*M246I* mutations showed significant positive cross-correlation with proliferation (S phase and G2M score) (**Fig. 6D, Supplementary Fig. 7E**). Thus, in this donor, the cells that had a *TP53p*.*Q136X* truncating mutation accompanied by chr17p CNLOH, as well as cells that had a *TP53p*.*M246I* mutation with chromosome 3q amplification, had high cell cycle gene expression. This suggests that a proliferative advantage could be imparted through complete loss of *TP53* (two-hit mechanism). On the other hand, the effect of partial loss of *TP53* (one hit) is unclear as more than 75% of cells carrying *TP53p*.*M246I* also carry a chr3q CNV event, an amplification which has been shown to lead proliferation in ESCC^104^.

While overall the genomes of *TP53*-mutant cells were stable without prevalent aneuploidy (**Fig. 5A**), we did observe multiple amplifications and deletions along chromosome 9q in the *TP53p*.*Q136X*-mutant clone (primarily gain events) (**Fig. 6E**). These events were specific to this clone and were not observed in the other *TP53* mutants. Expression levels of genes along the impacted chromosome regions corresponded with copy number (**Supplementary Fig. 7F**). Specifically, amplified regions (3-4n copy-number states) in *TP53p*.*Q136X*-mutant cells overlapped with the *CKS2* gene (**Fig. 6E**), encoding the cyclin-dependent kinase subunit 2 that promotes cell proliferation and is an established cancer gene^105^, associated with higher expression, tumor growth and poorer clinical outcome in esophageal squamous cell carcinoma^106,107^, potentially nominating *CKS2* expression as one of the contributing factors to higher proliferation of *TP53p. Q136X*-mutant cells. Together, our results show that *TP53*-mutant clones in normal esophagus tissue are mainly euploid, consistent with a model where cancer is driven by biallelic *TP53* loss together with obligate acquisition of a growth-promoting aneuploidy event to fuel clonal expansion. Recent bulk whole-genome sequencing analysis of normal aging esophagus identified biallelic disruption of *TP53* in 3 out of 11 subsetted *TP53*-mutant samples (harboring at least one driver variant with variant allele frequency (VAF) > 0.5), with these samples exhibiting overall high genome stability^108^, consistent with our observation of mainly euploid cells that lack wild-type *TP53* (e.g., the *TP53p*.*Q136X* clone with 17p CNLOH), which persisted with biallelic *TP53* loss for multiple years.

Collectively, our analysis uncovers four distinct phenotypic patterns in *TP53*-mutant cells. We observed predicted high-impact *TP53*-mutant clones without co-occurring CNVs or impact on proliferation (e.g., *TP53p*.*R175_H178del, TP53p*.*C238Y* and *TP53p*.*G279V*), as well as a clone with biallelic *TP53* loss (*TP53p*.*P151A* clone with chr17p CNLOH) that similarly lacked aneuploidy or an associated increase in proliferation. In contrast, we identified a clone with biallelic loss of *TP53* (*TP53p*.*Q136X* with chr17p CNLOH) that harbored aneuploidy events on chromosome 9 along with higher expression of S/G2M genes, suggesting increased proliferation. Finally, we observed a *TP53*-mutant clone (*TP53p*.*M246I*) that lacked evidence for biallelic loss (no accompanying CNLOH affecting chr17p) of *TP53*, but that had acquired an aneuploidy event (amplification of chromosome 3q) and displayed higher proliferation. These results support the model that while *TP53*-mutant cells have a fitness advantage due to decreased differentiation (**Fig. 4G**), only cells that additionally harbor CNV events have a proliferative advantage. Such clones are indeed much more frequently found in cancer in comparison to normal, albeit aged and risk-exposed, tissue^3,4^.

### Prevalent and convergent chromosome 9q copy-neutral loss of heterozygosity in aging human esophagus

Due to the technical limitations in amplifying the GC-rich *NOTCH1* locus, *NOTCH1* SNVs were not detected. However, CNLOH at chromosome 9q, encompassing the *NOTCH1* locus, has previously been observed alongside *NOTCH1* SNVs in normal esophagus tissue in bulk sequencing analysis. Whole-exome sequencing of microdissections uncovered CNLOH affecting the entire chromosome 9q arm^109^, in a substantial subset of samples (49/130; 38%) of normal esophagus samples^4^. In addition, CNLOH events on chromosome 9q with distinct breakpoints were identified using bulk whole-genome sequencing data of 2mm^2^ microdissections from esophageal epithelium (12/21; 57%)^3^. Moreover, both maternal and paternal chromosomes have been seen affected in Genotype-Tissue Expression (GTEx) bulk RNA data (137/948; 14% of normal esophagus samples)^110^. While these bulk studies highlight the widespread occurrence of CNLOH, the true prevalence of these events may still be significantly underestimated, especially in context of bulk RNA data that were sampled from a range of tissue sizes^110^, where large samples are likely to contain multiple clones at low frequency, many of which can fall below detection thresholds. Furthermore, when clones with different haplotypes are affected at similar frequencies within the same sample, their opposing signals may cancel each other out, effectively masking the presence of CNLOH. SMART-PTA enables detailed characterization of chromosome 9q CNLOH events within each sample, including not only their precise breakpoints and affected haplotypes but also their co-occurrence with SNVs, the phenotype of affected cell populations and the timing of acquisition (**Fig. 7A, 7B, 7C, Supplementary Fig. 8A, 8B, 8C, 8D, Supplementary Fig. 9**).

**Figure 7:**
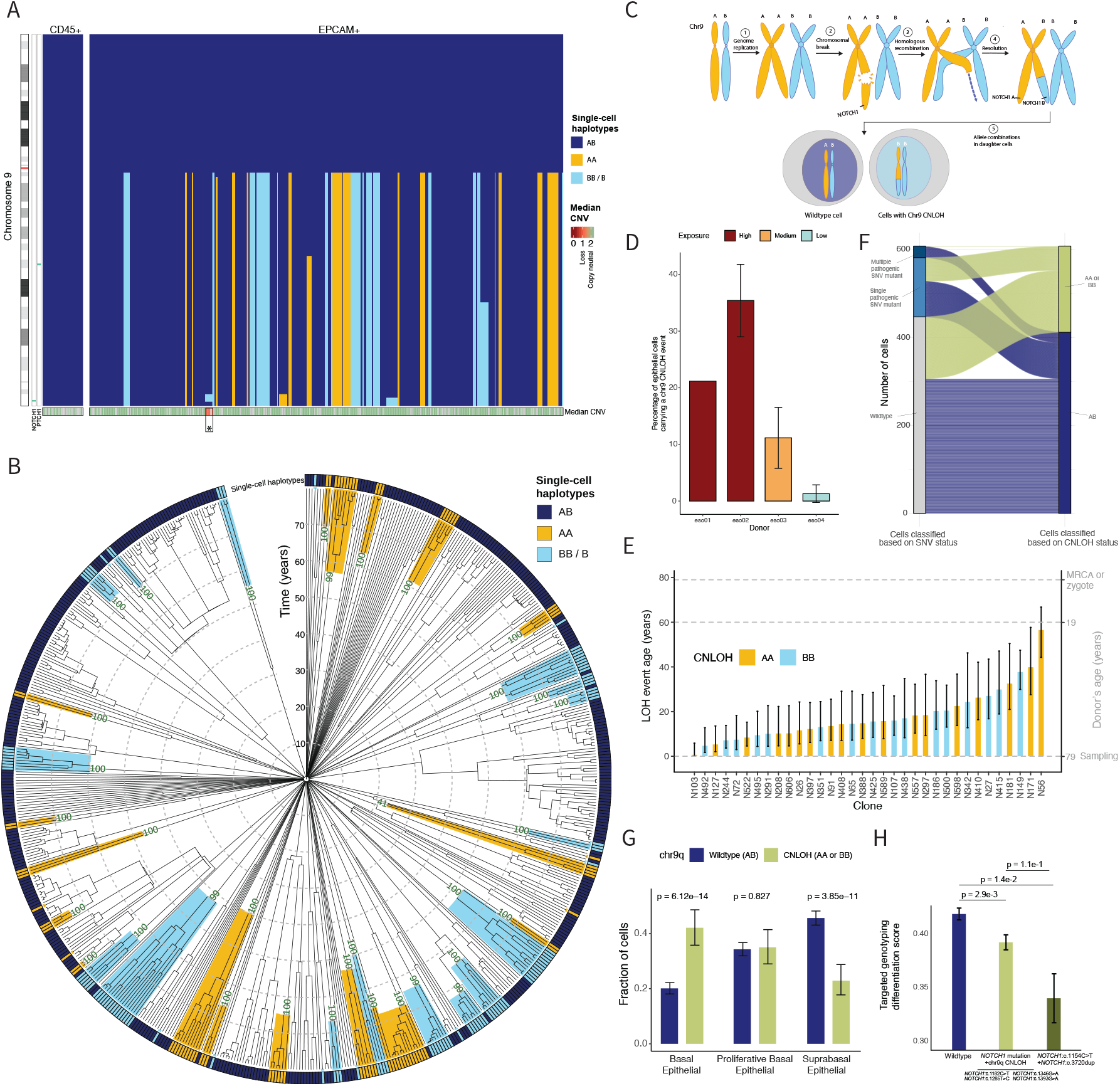
Large, independent copy-neutral loss-of-heterozygosity events affecting chromosome 9q occur frequently in the aging esophagus. **A**, CNLOH map of chromosome 9 for all immune (CD45+) and epithelial (EPCAM+) cells from donor eso02. Immune and epithelial cells are phylogenetically sorted within each group. The phylogeny was built after excluding SNVs detected in chromosome 9 to minimize the impact of residual germline variation (Methods). *NOTCH1* is located towards the telomeric end of the q arm of chromosome 9 (centromere in red on the chromosome, *NOTCH1* and *PTCH1* locations are indicated in green to the right of the chromosome). Single-cell haplotypes are assigned as AB (both parental alleles) or as having CNLOH, with AA or BB haplotypes. Median copy number across ∼50-kb non-overlapping windows is displayed in the bottom bar. Cells with LOH show median CNV for the affected region while wild-type cells show median for the whole chromosome arm. Median CNV values for cells with a coverage MAD higher than 0.2 (Supplementary Fig. 8D) are masked (grey). The asterisk indicates the *TP53*.*pQ136X* clone that had a deletion (non copy-neutral) event at the *NOTCH1* locus (as shown in Fig. 6E). Plots for the three other donors are in Supplementary Fig. 9A, 9B, 9C. **B**, Single-cell phylogeny of n = 608 cells from donor (eso02), with age estimated from branch lengths labeled by decade and indicated with dashed grey lines. Each leaf of the tree represents a single cell. Allelic status at the *NOTCH1* locus on chromosome 9q is annotated on the phylogeny as wild-type (AB) or having CNLOH, showing the retained parental haplotype (assigned as A or B). Numbers on the phylogeny indicate bootstrap values. Phylogenies for the other three donors are in Supplementary Fig. 9D, 9E, 9F. **C**, Schematic of somatic CNLOH at chromosome 9q, illustrating how after DNA damage, the homologous undamaged chromosome is used as a template for repair. A and B refer to parental haplotype at the terminal end of chromosome 9q covering the *NOTCH1* locus. **D**, Percentage of epithelial cells carrying a chr9q CNLOH event across four donors. Error bars indicate standard deviation. Donors are classified according to exposure risk: high, medium or low. **E**, Estimated timings of CNLOH events, ordered from most recent to oldest in donor’s life, with impacted parental allele, A or B, indicated. CNLOH age is estimated as the age of the MRCA of the cells carrying the CNLOH. **F**, Sankey plot showing the number of cells according to nonsynonymous (ns) SNVs filtered for CADD score (≥ 20) in driver genes and chr9q CNLOH status. Wild-type cells have no detected nsSNVs, single mutant nsSNV cells have mutations in one driver gene, and multiple nsSNV cells have mutations in more than one driver gene. AB, no chr9q CNLOH; AA or BB, chr9q CNLOH. **G**, Mean fraction of cells without chr9q CNLOH (AB) or with chr9q CNLOH (AA or BB) across three epithelial cell types (basal, proliferative basal and suprabasal). Error bars represent 95% Clopper-Pearson confidence intervals, and *P* values were calculated using a χ^2^ test. **H**, Differentiation scores of single cells with chr9q LOH + *NOTCH1* mutation (n = 334 cells), double *NOTCH1* point mutation (n = 39 cells) or wild-type cells (n = 1,258) obtained from aged esophagus samples and sequenced with targeted single-cell DNA-seq and matched RNA panel^84^. Error bars represent standard error of the mean (SEM), two-sided Student’s t-test.

At single-cell resolution, we first revealed that these CNLOH events were more prevalent than reported in bulk (14-57%), present in ∼85% (11/13) of the biopsies, and were restricted to epithelial cells (**Fig. 7A, Supplementary Fig. 8C, Supplementary Fig. 9A, 9B, 9C**), consistent with somatic pressures in the esophagus selectively acting on the tissue epithelium. The proportion of cells carrying a chr9 CNLOH was higher in high-risk donors, reaching a mean value of 35.4% across biopsies in donor eso02 with history of smoking and alcohol use and 11.2% in donor eso03 with history of alcohol use (**Fig. 7D**). This supports the idea that CNLOH events arise through mitotic recombination as a repair mechanism for DNA damage, such as double-strand breaks, which may become more frequent with increased exposure. The low coverage of *NOTCH1* (**Supplementary Fig. 6F, 6I**; **Supplementary Table 5**) restricted our ability to call somatic variants at this locus, and we only identified one chr9q CNLOH-positive cell with *NOTCH1* mutation in our targeted capture panel data (**Supplementary Fig. 10A**), indicating that *NOTCH1* mutations do occur, but are challenging to detect. In addition to these prevalent CNLOH events at chr9q, we detected clones with chr11q, chr17p (with *TP53* mutation) and chr22 (with *EP300* mutation) CNLOH (**Supplementary Fig. 8A, 8B**).

Leveraging the phylogeny, we parsimoniously determined that chr9q CNLOH events arose independently and recurrently up to 34 (eso02), 17 (eso03) and 2 (eso04) times in only 32mm^2^ of the esophagus epithelium over donor lifespan (**Fig. 7B, Supplementary Fig. 9E, 9F**), again consistent with positive selection for these alterations. The events frequently expanded beyond the *NOTCH1* locus, including CNLOH impacting the entire 9q arm in the majority of clones (**Fig. 7A, Supplementary Fig. 8C, Supplementary Fig. 9A, 9B, 9C**). We also observed clones that contained shorter CNLOH events with different breakpoints along chr9q (n = 2 clones in eso01; n = 5 clones in eso02; n = 3 clones in eso03; n = 2 in eso04; **Fig. 7A, Supplementary Fig. 8C, Supplementary Fig. 9A, 9B, 9C**), reinforcing that these are independent events. As further evidence of the convergent evolution of chr9q CNLOH, both the maternal or paternal allele were affected (**Fig. 7A, 7B**). In donor eso02 with the highest level of CNLOH, the earliest 9q CNLOH event occurred around donor’s age 20 (CNLOH age ∼60 years), with steady occurrence of repeated events over time, where the most recent event occurred within the 20 years prior tissue collection (**Fig. 7E**). The independent origin of the CNLOH events was further validated by visualizing the somatic mutations in chromosome 9 that were not leveraged for tree building to avoid circularity. With the exception of clones N65 and N72, which might carry the same ancestral CNLOH, the remaining 32 clones (containing at least two cells) harbored unique homozygous mutations likely acquired in heterozygosis prior to the CNLOH, which occurred independently within each of the lineages (**Supplementary Fig. 10B**).

To gain a broader understanding of the clonal mutant landscape, we analyzed chr9q CNLOH events together with esophageal driver gene SNVs in eso02, identifying cells harboring both CNLOH and a driver gene mutation (**Fig. 7F**). There was no significant difference between chr9q CNLOH-positive and CNLOH-negative cells in proportion of cells with a nonsynonymous SNV (CADD ≥ 20) in at least one driver gene (0.28 versus 0.26, *P* = 0.27), indicating that the CNLOH event did not increase the likelihood of acquiring a subsequent mutation in an esophageal driver gene. Notably, we observed a striking depletion of *TP53* and *FAT1* mutants in the CNLOH-containing cells (**Supplementary Fig. 10C**). Whereas *TP53*- or *FAT1*-mutant cells were commonly seen in clones without chr9q CNLOH (58% cells with at least one driver mutation), these were almost entirely absent in clones with chr9q CNLOH (0.03%). Interestingly, decreased selection of *TP53* mutants in samples with *NOTCH1* loss has been reported, where targeted sequencing of 0.05mm^2^ esophagus biopsies showed that *TP53*-mutant clones had smaller VAF in high VAF *NOTCH1*-mutant clones from >70-year-old-donors^108^. We note that this bulk analysis was restricted to samples where a mutation was present in >50% of cells, while in contrast, our approach enabled us to assess mutation co-occurrence with CNLOH at the level of the single cell. Our results suggest that cells with chr9q CNLOH events are less likely to harbor mutations in *TP53* and *FAT1*, consistent with a model where clonal competition between mutant cells within the esophageal tissue^64,111^ constrains their growth, disfavoring acquisition of *TP53* or *FAT1* mutation in CNLOH+ cells.

To assess the potential functional impact of chr9q CNLOH, we next used the transcriptome information to analyze cellular phenotypes according to CNLOH status. *NOTCH1* dysregulation has previously been associated with biasing cells to an earlier differentiation stage^84^, where clonal advantage of *NOTCH1*-mutant cells over wild-type cells derives from symmetric divisions leading to fewer differentiated mutant progeny cells^112^. In this model, wild-type basal cells are driven to differentiate by neighboring *NOTCH1*-mutant cells, leading to net expansion of *NOTCH1*-mutant basal cells through loss of their wild-type counterparts through increased differentiation, rather than increased proliferation of the *NOTCH1*-mutant cells^33,112^. Consistent with this, we observed that proportion of cells in S and G2M phase for chr9q CNLOH-positive cells did not increase compared with CNLOH-negative cells (**Supplementary Fig. 10D**), suggesting that chr9q CNLOH does not affect proliferation. To assess potential differentiation biases in CNLOH clones, we analyzed the fraction of cells with or without chr9q CNLOH across basal, proliferative basal and suprabasal epithelial cell types. Notably, we observed that the earliest basal epithelial population had a significant increase in chr9q CNLOH-positive cells, and that the later suprabasal epithelial cells had the highest fraction of CNLOH-negative cells (**Fig. 7G**), consistent with *NOTCH1* loss preferentially biasing cells toward earlier states. To validate this finding, we performed an experiment using our single-cell targeted DNA and RNA co-capture assay^84^ applied to an aged esophagus sample. Here, we leveraged germline heterozygous SNPs captured by the amplicon panel to determine CNLOH at the *NOTCH1* locus and assigned epithelial differentiation state using the matched RNA (**Methods**). We found that cells with a *NOTCH1* mutation and LOH at the chr9q *NOTCH1* locus had earlier differentiation scores compared to wild-type cells, and that these differentiation scores were comparable to those of cells with biallelic *NOTCH1* loss (two distinct SNVs; **Fig. 7H**). These results demonstrate that the CNLOH-positive cells have a phenotype that is aligned with loss of *NOTCH1*.

Together, our phylogenetically-informed analyses enabled us to show that independent chr9q CNLOH events can occur multiple times over lifespans, providing a high-resolution window into how selection shapes the somatic genome in aging esophagus epithelium. While copy-number variation is a hallmark of Barrett’s esophagus and esophageal adenocarcinoma^113^, our single-cell view into normal esophagus revealed that these genomes also bear a high prevalence of CNLOH events that can bias cells towards earlier fate.

## Discussion

Somatic evolution inexorably proceeds after fertilization and throughout life, with progeny cells acquiring new genomic variants with each mitosis, resulting in high genetic heterogeneity within an individual. To accurately catalogue this complement of somatic genomes within each individual, genomic profiling at the single-cell level is required. Recent advances in scWGS technology have empowered analyses that uncover the somatic landscape of particular tissues across aging and disease^16,21,23,114^. Beyond capturing the extant somatic variation of a tissue, scWGS can also be harnessed to reveal the temporal dimensions of clonal evolution, with SNVs acting as intrinsic barcodes that uniquely identify single cells and enable construction of somatic phylogenies. This approach provides a temporal microscope to trace the evolutionary history of cell populations, much like how germline variation is used to build phylogenetic trees in species evolution. Pioneering single-cell lineage tracing studies in humans used WGS of single-cell derived colonies of primary bone marrow cells to chart clonal hematopoiesis, implementing phylodynamic analysis to infer the timing of clonal origin and patterns of growth^10^. Such approaches have quantified the diversity of somatic clones in the blood throughout aging, estimated clonal expansion timing and measured fitness effects of clones with different clonal hematopoiesis driver mutations^11,115^. These efforts have helped to define human somatic evolution in blood, providing foundational understanding of how populations of cells mutate, grow and compete over human lifespans.

Despite these advances, single-cell cloning followed by WGS is time-consuming, requires live cells that can be cultured ex-vivo, and lacks opportunity to assess matching single-cell phenotypes. While direct scWGS methods are now available, currently these are laborious and expensive, limiting broader application of scWGS to large-scale analysis of primary human samples, including solid tissues. Recent studies have taken advantage of the improved coverage and dropout rate afforded by PTA for sequencing single cells, but these have been restricted to sequencing tens of cells per patient^21–23^. To address these limitations, we developed a framework that automates and miniaturizes PTA for the efficient generation of high-quality matched single-cell DNA and RNA libraries from up to a thousand single cells within one week. Further combining this approach with cost-efficient UG sequencing and computational advances enables scaled production of single-cell whole genomes and transcriptomes, which can be used for the construction of large, phenotypically annotated phylogenies from human samples. Overall, our SMART-PTA framework led to major reduction in production time, reagent and consumables use, sequencing cost, and analysis time, all while generating higher-quality libraries than standard-PTA+RNA. These innovations represent a major step forward in our ability to interrogate human somatic evolution across non-malignant tissues, as we are able to reconstruct cell lineages in deep evolutionary time, quantify the timing of mutation events, characterize clonal expansions and assess the mechanisms that lead to enhanced fitness of mutant clones.

With this framework, we used single-cell SNVs to build high-resolution phylogenies from >2,700 single cells from phenotypically normal esophagus biopsy samples from four older donors, further annotating these with phenotypic information from the matched single-cell transcriptomes. Constructing these phylogenies posed a challenge equivalent to recent efforts in resolving deep evolutionary histories, such as the reconstruction of a global butterfly phylogeny^116^. To overcome this, we developed new strategies for tree search and branch length time estimation. Our phylogenies revealed non-monophyletic origin of immune and epithelial cells and different dynamics of the two cell types likely due to distinct ontologies and niche. For example, we reconstructed a phylogeny of tissue-resident immune cells, uncovering the expansion of a CD4+ T cell clone. We further harnessed phylogenetic analysis to determine cell migration patterns, similar to how genomic sequencing has been used to reconstruct human migration patterns in population genetics analysis^117^. In aging esophageal tissue, epithelial cells had high phylogenetic correlation with sampling region, indicating that there was low cell migration, which is consistent with rigid tissue architecture and stereotyped progression of cells during differentiation^37^ restricting motility. restricting motility. Notably, in a more advanced pathological setting, such as Barrett’s esophagus, such constraints can be overcome by metaplastic epithelial cells, which have been observed to span large esophageal segments^118^. In contrast to epithelial cells, the immune cell population had lower phylogenetic scores with sampling region indicating higher migration, aligning with their ability to mobilize locally in response to signals^57,62,63^. Future work integrating more detailed spatial profiling with phylodynamic analyses could further reconstruct cellular migrations during somatic evolution.

The samples analyzed here were provided by donors who were age 76 or older, with some donors reporting a history of smoking or alcohol use. Our phylogenies provided a window into the effects of aging, stress and exposures on the somatic evolution of the esophagus. In one example, the ability to draw on mutations in different parts of the tree for mutational signature analysis provided a view of mutagenic exposures across the donor lifespan. Through this analysis, we revealed emergence starting in adulthood of tobacco-related/alcohol signatures in donors with exposure history, as well as evidence corresponding to smoking cessation prior to sampling. Previous bulk sequencing analyses from expanded HSC colonies have analyzed proportions of mutation signatures along branch lengths calibrated by number of mutations^119^. In our analyses, deep sampling of epithelial clones restricted in space allowed us to obtain cells that are more related than HSCs in blood, creating long internal branches which contrast with the short internal branches that have very few mutations present in most colonies^119^. Our large single-cell phylogenies additionally allow us to partition exposure effect over a donor’s lifespan, providing enough power to confidently assess genomic insults at early versus more recent time, including the most recent exposures affecting terminal branches. Thus, our work demonstrates that an accurate record of exposure is imprinted on the somatic genome, and that these patterns can be deciphered using single-cell phylogenies.

Our high-resolution phylogenies additionally allowed us to assess decades of somatic evolution in solid human tissue by identifying clones with driver gene mutations. Such driver genes have been shown to be positively selected in the esophageal epithelium^3,4^, providing a fitness advantage within the aging tissue environment. Our analysis revealed that donors with a history of exposure had an overall higher burden of total mutations compared to those donors without exposure history. Interestingly, when we assessed positive selection at the gene level, we found that *TP53* and *FAT1* showed similar degree of positive selection in donors, regardless of exposure history. This finding is limited by the small number of individuals analyzed; however, we speculate that in the specific context of the esophageal epithelium, carcinogenic exposures promote somatic evolution more due to their mutagenic effect than by altering the selection pressures (e.g., promoters^120,121^). This finding may be contrasted with selection of hematopoietic stem cells with chemotherapy^120^, or the impact of air pollution on clonal mosaicism in the lung^121^, where mutagenic agents were also shown to exert specific selection pressures aiding clonal outgrowth.

Our single-cell genome-wide profiles revealed that aneuploidy events were rare, indicating that similar to normal breast epithelial tissue^14^, the esophageal epithelium remains mostly euploid, even upon aging and risk exposure. We further harnessed our data to define CNLOH events and assess these in relation to SNVs and across time over donor age. By simultaneously analyzing SNVs and copy-number, we were able to identify clones with biallelic *TP53* loss (*TP53* mutation combined with CNLOH covering the *TP53* locus), observing instances where clones lacking intact *TP53* existed for years, without extensive aneuploidy. It is established that complete loss of *TP53* leads to clonal expansions by allowing cells that have acquired fitness-enhancing aneuploidy events to proliferate without normal genome integrity checkpoints^113,122^. Thus, loss of *TP53* itself does not immediately lead to a growth advantage per se, but allows for unchecked expansion of cells with chromosome instability and aneuploidy events, demonstrated by diploid *TP53*^*-/-*^ mutant epithelial cells being outcompeted by aneuploid *TP53*^*-/-*^ mutants *in vitro*^123^. This is consistent with our data, where the only two *TP53*-mutant clones that had evidence of increased proliferation were also the only two clones that harbored CNV events, and a clone with biallelic *TP53* loss that lacked CNVs did not show increased proliferation, further reinforcing that acquisition of a CNV is needed to bestow a growth advantage to *TP53*-mutant cells, even when *TP53* is completely lost. Previous work has associated *TP53* inactivation and accompanying genome instability with Barrett’s esophagus, early esophageal adenocarcinoma and esophageal squamous cell carcinoma, showing that *TP53* loss and genome instability initiate genome evolution^101,113,124^. Whether the *TP53-* mutant clones that we observe represent precursor lesions that will subsequently acquire more catastrophic genomic events (whole-genome doubling, chromothripsis, extensive aneuploidy) that are associated with transition to malignancy remains unknown. Within the physiologically normal tissue environment, it is possible that the growth of such clones remains in check through neighboring cells^88^, and that transformation would require a more pervasive disruption to esophageal homeostasis than exists in these donors currently.

Although overall the cells that we examined were euploid with a lack of large CNV events, we also observed a striking pattern of prevalent CNLOH at chromosome 9q. Such events have been previously observed through bulk sequencing of biopsies, with the majority of identified chr9q CNLOH impacting the entire chromosome arm^3,4^. This may limit quantification of single versus multiple events within a single sample, particularly for large biopsy regions^110^, where rarer events may be missed. Our scWGS analyses revealed that up to a third of epithelial cells harbored chr9q CNLOH in high-risk donors. Notably, these events impacted both the paternal and maternal alleles and arose independently many times. In one donor, the phylogenetic analysis revealed over 30 separate instances of chr9q CNLOH acquisition in ∼600 cells). Furthermore, we observed that these events were acquired steadily over donor lifespan, where in high-risk donor eso02 the earliest CNLOH event occurred at around 20 years old, followed by an additional event arising every ∼two years. This gradual pattern of accumulation of CNLOH events over time in the esophageal epithelium contrasts with observations from the gastric epithelium, where CNV events, including various chromosomal trisomies, arise at specific time points (punctuated evolution), potentially related to an infectious or inflammatory event^125^. Interestingly, the chr9q CNLOH burden was increased in donors with higher levels of exposure to alcohol and/or tobacco, consistent with a role for mutagen-induced DNA damage in promoting these repair events.

Our matched genome and transcriptome profiles enabled us to map genotypes to phenotypes in order to gain insights into the functional effects of driver mutations. Our analyses revealed that mutations across multiple positively selected esophageal driver genes, including *TP53* and *FAT1*, converged on an earlier differentiation state basal phenotype. We also observed this earlier differentiation skew in chr9q CNLOH-positive cells, consistent with a *NOTCH1*-mutant phenotype. Previous studies have proposed a model whereby *NOTCH1* loss disrupts normal epithelial differentiation by promoting differentiation of neighboring wild-type cells, resulting in expansion of mutant and loss of wild-type cells from the early basal layer^33^. Our findings extend this mechanism to other esophageal driver genes, indicating that in the phenotypically normal esophagus, there is high selective pressure for mutations that drive differentiation bias without disruption of normal epithelium structure. Notably, overall we did not observe enhanced proliferation across mutant cells, suggesting that the presence of driver mutations is insufficient to promote increased cell proliferation, and that other factors, such as CNV acquisition, are needed in order for a cell to gain a proliferative advantage that leads to clonal expansion.

While we have presented large single-cell, phenotypically annotated phylogenies from primary solid tissue samples, we note several limitations with our approach. While PTA offers unbiased genome-wide amplification, there are technical limitations to amplifying regions with driver genes, such as those with increased GC content. The aging esophagus mutational landscape has been shown to be dominated by *NOTCH1* mutations, yet *NOTCH1* is not amplified adequately for variant calling. Single-cell targeted DNA sequencing may serve as an alternate tool for obtaining a snapshot in time of the driver mutational landscape^84^. Technological advances will be required to overcome the current limitations of single-cell whole-genome amplification methods for a more even coverage across all genomic regions. In addition, despite our strategies to control cost through automation, miniaturization and UG sequencing, constructing phylogenies from thousands of single cell whole-genome sequences still comes with significant expense and computational cost. These challenges will need to be overcome to build even larger phylogenies, which will enable the capture of more somatic variation, allow for more powerful phylodynamic analysis and provide a more accurate map of cell ancestries. In the future, we envision that continued decreases in the cost of sequencing combined with novel protocol innovations will enable even higher throughput, to define and quantify processes during somatic evolution during human lifetimes.

Thus, these and future innovations will empower new discoveries about the mechanisms and dynamics of human somatic evolution as central to advancing our understanding of aging, chronic disease, precancerous states and malignant transformation. With this framework to directly profile matched single-cell genomes and transcriptomes from clinical samples with increased scale and resolution, we will begin to uncover the complement of genetic heterogeneity represented by the trillions of somatic genomes that exist within an individual. The scalable exploration of somatic genetics will help unravel how somatic variants differentially impact clonal expansions, cellular phenotypes, differentiation landscapes and tissue homeostasis. Phylogenetic analysis further adds the dimension of time, enabling reconstruction of somatic evolutionary histories over human lifespans. Collectively, these advances will help propel the next frontier of human somatic genomics, where it will be possible to delineate evolution of cells during aging and disease, incorporating the temporal dimension of somatic variation. Almost 50 years after Sulston and colleagues traced the development of each cell in a nematode through direct visualization of mitosis and cell fate determination^126^, we are inspired to envision how single-cell genomes with phenotypic labeling could be leveraged to analogously dissect elements of human somatic evolution.

## Acknowledgements

We thank all members of the Landau lab for critical input. We thank Jane Park for outstanding technical support and Henry Rui He for operational support. We thank Alexei Drummond and Kylie Chen for their advice on how to run Phylonco with relaxed models. We thank Iraj Eshghi for discussion about 2D growth models. We thank Iñigo Martincorena for providing processed trinucleotide profiles of buccal swab samples and extracted signatures from his study. We thank Alexey Kozlov for insightful discussions on parallelizing CellPhy. We thank Vladimir Svetlov for assistance with p53 structural modeling. This work was supported in part by the Clinical and Biospecimen and Research Core of the Columbia University Digestive and Liver Disease Research Center (P30 DK132710). D.A.L. is supported by the Burroughs Wellcome Fund Career Award for Medical Scientists, the Vallee Scholar Award, the Blood Cancer United Scholar Award and the Mark Foundation Emerging Leader Award. This work was supported by the National Cancer Institute (R33 CA267219), the National Institutes of Health Common Fund Somatic Mosaicism Across Human Tissues (UG3NS132139) and the National Human Genome Research Institute, Center of Excellence in Genomic Science (RM1HG011014). This work was made possible by the MacMillan Family Foundation and the MacMillan Center for the Study of the Non-Coding Cancer Genome at the New York Genome Center.

## Author Contributions

T.P., D.J.Y., and D.A.L. conceived the project and devised the research strategy. I.R., D.J.Y. and C.H. developed, implemented and optimized SMART-PTA protocol. D.J.Y., C.H., J.S. N.D.O., and I.R. performed experiments. T.P., J.Z. and S.K. developed the analytical pipelines for processing the phylogenetic data. T.P., D.J.Y., N.M., J.Z., S.K., J.H., F.F., R.R., N.Ru., J.S.S., N.W. and A.G performed computational analyses. S.R., A.R.D., S-H.Y., B.P., A.P.C. and N.Ro. provided analytical expertise and advice. A.Y., N.K. and S.O. provided sample eso01 and critical discussion. K.G. and J.A. provided samples and advised on biological interpretations of the data. C.P. and I.R. advised on biological interpretations of the data. T.P., D.J.Y., C.P., I.R. and D.A.L. wrote the manuscript. All authors reviewed and approved the final manuscript. T.P. and D.J.Y. contributed equally to this work.

## Code and Data Availability

Code underlying these analyses is available on GitHub: scWGA quality control: https://github.com/tamaraprieto/SingleCheck

Variant Calling/Joint Genotyping: github.com/jzinno/darkshore Distributed ML Tree Search: github.com/jzinno/cloudCellphy RNA analysis: github.com/jzinno/rsoRNA

The single-cell genomics data will be deposited in EGA. Accession codes will be made available upon publication.

## Methods

### *In vitro* DLD-1 evolutionary experiment

An *in vitro* single-cell evolutionary model was devised by systematically expanding and cloning individual DLD-1 (ATCC CCL-221) cells over a span of four months. Initially, cells from the primary culture were cultured in RPMI-1640 supplemented with 10% fetal bovine serum for a period of 30 days. A segment of the resultant parental population was cryopreserved (population A), while the remaining cells were utilized to initiate the first filial generation through single-cell cloning. Single-cell cloning involved diluting cells in media to a concentration of <1 cell per 100μL and dispensing 100μL of the suspension into cloning cylinders. Successful clones underwent expansion over a ∼30-day period, with the size of the culture dish periodically increased to accommodate population growth. A subset of expanded clones (n = 12) was cryopreserved, and the rest were used to establish a second filial generation of cell clones. This cycle of expansion (lasting 30-33 days), cryopreservation, and cloning was iterated for three filial generations, totaling ∼90 days. Following thawing, cryopreserved population A and selected clones (n = 7) from the first (B, E), second (C), and third (D, F, G, H) filial generations, were independently flow sorted into 96-well plates. The isolated cells underwent single-cell whole-genome amplification using PTA (ResolveDNA v1, BioSkryb). Subsequently, PTA products were fragmented, and libraries were prepared from the PTA amplification product using the Kapa HyperPlus workflow. Libraries resulting from two single-cell amplifications of the parental population (A1 and A2) and two single cells from each of the seven filial clones (B1, B2, E1, E2, C1, C2, D1, D2, G1, G2, F1, F2, H1, and H2) were later subjected to conversion to Ultima Genomics (UG) compatible libraries and sequenced on their platform at the New York Genome Center (NYGC) at an average of 30× per cell. Illumina sequencing data from a single-cell colony derived from the same DLD-1 cell in an independent study was downloaded to use as germline control^127^.

### Ultima Genomics (UG) DLD-1 scWGS processing

The UG reads were processed as previously reported^24^. Cram files with different read groups for the same single-cell were merged together using *samtools merge*. For calculating the original sequencing depth of the cram or bam files, we computed the number of primary alignments times the average read lengths for the first million reads and divided by the genome length. The alignment files were downsampled to 15× the genome length using *Picard DownsampleSam*. A customized version of *DeepVariant*, optimized for UG sequencing data (see below), was used to generate single-cell GVCFs and VCFs.

### Matched Illumina DLD-1 scWGS

Four libraries from clones F and H (F1, F2, H1 and H2) were sequenced using an Illumina NovaSeq6000 at NYGC, ensuring an average sequencing depth per cell exceeding 15×. Illumina reads were trimmed with *Cutadapt* (v. 2.10), mapped to human reference version hg38 using *BWA MEM* (v. 0.7.17) and further processed with *Picard* (v. 2.18.17) to mark duplicates and downsample to 15× as above. *Deepvariant* was run with default parameters (v. 4.5.0 via *parabricks*) to generate one VCF per cell.

### Benchmark UG scWGS variant calling

The F1, F2, H1 and H2 VCFs for both UG and the matched Illumina data were independently annotated with dbSNP 150 using *ANNOVAR* (June 2020 version). Those variants present in dbSNP and also detected in a single-cell colony^127^, were annotated as high-confidence germline sites. Variants with a quality of at least 30 and absent from dbSNP150 and the colony, were considered as somatic SNVs. Tables listing cell-specific variants were intersected in R to quantify how many UG-detected variants were also present in matched Illumina single-cell data. The trinucleotide context of each SNV was retrieved from the reference genome using the *scanFa* function from *Rsamtools* [doi:10.18129/B9.bioc.Rsamtools], and 96-context mutation counts were input into *deconstructSigs* to estimate the contributions of the six most common SBS signatures. Similar SBS exposures were consistently identified across SNV sets and cells.

### DLD-1 phylogenetic reconstruction and benchmark

For building the *in vitro* phylogeny, GVCFs of the 16 single cells were jointly genotyped using *GLnexus*(v1.4.1) and the resulting VCF was used as input for *CellPhy*^47^ with GT10+FO+E as model. The *in vitro* experiment supports a single phylogenetic tree containing three polytomies of sizes 3, 4, and 3, corresponding to 135 fully bifurcating resolutions. This number was calculated using the formula (2k−3)!! for each multifurcation of size *k*, and stands in stark contrast to the roughly 6.4×10^22^ possible rooted topologies that exist for 16 taxa. Using the R package *Quartet*, we assessed the similarity between our inferred tree and the true tree with polytomies, obtaining a score of 100%.

### Single-cell whole-genome amplification and reverse transcription

We optimized the Single-cell Miniaturized Automated Reverse Transcription and Primary Template-directed Amplification (SMART-PTA) workflow by integrating the Mantis Liquid Dispenser (Formulatrix) with the ResolveOME reagents (BioSkryb). For reverse transcription, PTA, fragmentation, end repair, and library amplification reagents from ResolveOME v1 assays (BioSkryb) were loaded onto a Mantis Liquid Dispenser according to manufacturer’s recommendations, using 0.25× of original reagent volumes per well. Briefly, cell membranes were lysed for reverse transcription of mRNA. Nuclear membranes were lysed to access genomic DNA for PTA, generating 1-2kb fragments of amplified genomic DNA. Generated cDNA was separated from PTA products using bead-based separation, and both cDNA and PTA products were fragmented before adapter ligation and library amplification. For all clean up steps, we utilized the G.PURE (Cytena) for automated DNA purification to retain library quality.

cDNA libraries were sequenced with NovaSeq6000 at paired-end 150bp with target of 2 million reads per cell. WGS Illumina libraries were then converted to Ultima Genomics (UG) libraries by PCR-based conversion kit (Ultima Genomics), and sequenced on UG100 sequencer at target of 15× coverage.

### Standard-PTA+RNA vs SMART-PTA benchmark

To compare the performance of standard-PTA+RNA and SMART-PTA, we analyzed cells from each protocol. Standard-PTA cells were obtained from donor eso01, and SMART-PTA cells from donor eso02. From cells with >1 million RNA-seq reads, we randomly selected 50 per protocol. For each cell, genome-wide DNA quality was assessed using *SingleCheck* (https://github.com/tamaraprieto/SingleCheck) with the N parameter on UG-processed BAM files. Transcriptome quality was evaluated using our single-cell RNA-seq pipeline (see RNA-seq Data Analysis). We computed genome metrics including coverage breadth, mappability, and coverage uniformity (defined as the inverse of the median absolute deviation [1/ MAD] of read coverage across fixed-size non-overlapping windows of 10 Mb). Transcriptomic metrics included the number of detected transcripts, and the proportion of ribosomal and mitochondrial reads. Group-wise comparisons between protocols were performed using the two-sided Wilcoxon rank-sum test.

### Regional sampling, biopsy processing, isolation and, cell sorting of esophageal tissue

For eso01, three adjacent biopsies were taken with 2mm forceps. Biopsies were rinsed in PBS and treated with Dispase for 10 minutes at 37 °C, while being transferred to a solution of 0.25% Trypsin EDTA for 10 minutes at 37 °C. Cells were strained and reaction stopped with Soy Bean Trypsin Inhibitor.

For eso(02-04), fresh esophageal biopsies were obtained from donors during upper endoscopy. Biopsies were taken from two sites spaced 3 cm apart in the proximal 6 cm of the esophagus and two sites spaced 3 cm apart in the distal 6 cm of the esophagus. From each site, two biopsies were obtained. Forceps of approximately 2mm in diameter were used for collection of biopsies and rinsed in sterile PBS prior to sampling in distinct sites to minimize contamination. Each biopsy site location was recorded as distance from the incisors (**Supplementary Table 1**).

Biopsies were mechanically disrupted and dissociated via trypsinization from the muscle layer (eso02, eso03) or with Liberase (eso04). Both reactions were stopped with 4% FBS in KSFM media.

All cell suspension was stained and epithelial (EPCAM+CD45-) and immune (EPCAM-CD45+) single cells were sorted into 384-well plates with Symphony S6 sorter (BD).

### RNA-Seq Data Analysis

Pre-processing of raw FASTQ files was conducted using *FastP* (v. 0.23.4), which included trimming of template switching oligonucleotide (TSO) sequences and homopolymer tails ≥10 bases, alongside the application of default filtering parameters. Trimmed reads were aligned to the hg38 reference genome using *STAR* (v. 2.7.1a) in two-pass mode, with the stranded parameter set to ‘no’. Gene-level quantification was performed using *HTSeq* (v. 0.12.3) with default settings on the aligned single-cell data. The resulting raw count data per cell were consolidated into a single matrix using bash. T cell receptor (TCR) reconstruction was carried out using *TRUST4* (v. 1.0.13) with hg38 reference resources on the trimmed FASTQ files. A nextflow workflow (scRNA.nf) is provided in https://github.com/jzinno/darkshore. Raw gene expression counts were then pre-processed and normalized in Python (v. 3.11) using *scanpy* (v. 1.9.8). Cells with fewer than 200K read counts were removed based on knee point. The top 1,000 most variable genes were identified using *scanpy. pp*.*highly_variable_genes* function, with the *Seurat* v3 flavor for the selection criterion. To account for batch effects, embeddings were generated using *scVI* (v. 1.1.1), which models the gene expression data while factoring out batch covariance. Donor, sequencing batch, 384-well plate and 96-well plate identity were included as categorical covariates. A nearest-neighbors graph was constructed based on the resulting embeddings (*scanpy*.*pp*.*neighbors*), and cluster analysis was performed using the Leiden algorithm (*scanpy*.*tl*.*leiden*). UMAP was applied to the nearest-neighbors graph to visualize the data in a reduced-dimensional space. Cell types were assigned to Leiden clusters by evaluating known marker genes (**Supplementary Table 3**) and gene set scores across the identified groups. (https://github.com/jzinno/rsoRNA). Pseudotime analysis was performed via diffusion pseudotime (*scanpy*.*tl*.*dpt*) on a diffusion map *(scanpy. tl*.*diffmap*) over cell embeddings.

### DNA-seq data processing and variant calling

DNA reads were demultiplexed, trimmed, and mapped to the human reference genome (hg38) using Ultima Genomics proprietary software. The resulting cram files were combined per cell and read group sample tags standardized using *samtools* (v. 1.19) functions *merge* and *reheader*, respectively. Quality control was performed using *SingleCheck* with the N option. We introduce a distributed GPU accelerated pipeline (https://github.com/jzinno/darkshore) for subsequent variant calling analysis.

The single-cell samples were concurrently duplicate marked (*GATK MarkDuplicates*) and variant called using a pre-release version of NVIDIA Parabricks *Deepvariant* on L40S GPUs based on v. 4.1.2. Per cell variant calls were then collated for joint genotyping with *GLNexus* using an UG-specific configuration. Somatic variants were annotated using *ANNOVAR* (June 2020 version, -protocol refGene, dbNSFP v4.2c, COSMIC v70, dbSNP avsnp150, ExAC v0.3, ClinVar 20220320).

### Somatic variant identification

The original *Sequoia* script (accessed April 2024), a tool for calling somatic mutations from single-cell data without matched bulk normal^42^, was modified to accept interval data and further adapted to compute the probability and statistical significance of mutation placement on artificial outgroup versus ingroup branches, incorporating code from the *treemut*^49^ (v. 1.1) R package. The resulting VCF with Sequoia annotations was normalized and filtered using *BCFtools* (v. 1.15.1) to exclude low-quality variants (QUAL < 20) and low-quality cells. Cells were excluded if they had <50% genome breadth and either a median absolute deviation (MAD) > 0.5 or sequencing depth < 3x; > 5% unmapped reads; or <50% breadth and were annotated as CD45^+^ immune cells. Variants present in dbSNP build 150 (avsnp150), mapped to the outgroup, and with Sequoia germline q-values > 0.00005 were removed as putative germline. From the remaining variants, those recurrent in the first three donors or with a single-cell mean depth >30 were excluded. Variants not found in dbSNP and with a Sequoia rho > 0.5, or any variant with rho > 0.65, were retained as candidate somatic mutations.

### Germline variant selection and recall calculation

Germline variants were filtered using *BCFtools* (v. 1.9). Variants present in dbSNP, with a germline quality score greater than 0.00005 and a mean sequencing depth greater than 3 were retained as high-confidence germline sites. From this filtered set, heterozygous germline SNPs were selected by keeping only sites tagged with ‘GT=het’ and an allele frequency between 0.25 and 0.75.

For each cell, we extracted read counts for the reference and alternative alleles at heterozygous germline sites from the VCF using *BCFtools*. To assess recall, the proportion of heterozygous sites with at least one alternative allele read was computed per cell.

### Phylogenetic analysis

Phylogenies were built using a HPC distributed version of *CellPhy*^47^ (0.9.3git *CellPhy* branch) accessible at https://github.com/jzinno/cloudCellphy with a VCF containing the somatic SNVs as input and choosing GT10+FO+E as model. Somatic variants were excluded from phylogenetic reconstruction if they were present in fewer than two cells, had missing genotypes in over 50% of the samples, or showed a mean depth per cell less than or equal to 9. These filters were applied to reduce computational complexity while ensuring that only variants with sufficient data quality and informative value contributed to the analysis. We further excluded somatic mutations on chromosome 9 to prevent biases introduced by CNLOH events. In such cases, multiple germline heterozygous sites within the region can be misclassified as independent somatic mutations (due to genotype changes from 0/1 to 1/1), potentially inflating support for the tree despite stemming from a single somatic event. Importantly, all these excluded mutations were subsequently rescued and reintegrated into the phylogenetic backbone during mutation mapping.

### SMART-PTA error classifier

We built a PTA-error classifier that does not rely on a healthy matched sample. We obtained the following values per somatic SNV site: alternative and total read counts within the merged BAM of RNA libraries, presence in REDI RNA editing database (http://srv00.recas.ba.infn.it/atlas/), lowest coverage MAD score (coverage dispersion) among cells, highest variant allele frequency among cells and highest alternative allele count among cells. We further adapted code from *PTATO*^43^ to generate upstream and downstream 10-bp sequence context including the positive strand reference base in the middle, distance to the closest gene (0 if within a gene), DNA strand transcribed in the closest gene, distance to the closest simple repeat, replication time of the region, 200-kb window index, residual of the Loess regression of variant allele frequencies of heterozygous germline SNPs, and *P* value of a binomial test for the alternative. Putative true positives and false positives were identified by classifying variants under the assumption that each cell contains only two haplotypes. Therefore, reads that span both a somatic SNV (heterozygous) and a nearby germline heterozygous SNP (hetSNP), referred to as linked reads, should support no more than two consistent haplotypes^44^. In order to obtain linked reads, BED files containing somatic variants were intersected with VCFs containing high-confidence germline SNPs using *bedtools* (v. 1.9). Haplotype counts for each variant-hetSNP pair were calculated from the single cell BAM files using a custom python script after filtering out reads by edit distance (<= 5), base quality (>= 15), and mapping quality (>= 20). Multiple metrics were calculated from the linked-read information in R including the genomic position of the closest heterozygous germline SNP, median Shannon diversity index of linked-read counts across cells, number of linked-read counts supporting any of the four expected single-cell haplotypes, number of linked-read counts supporting other alternative haplotypes, list of cell identifiers in which the linked-read approach was applied, proportion of in which the site was classified as a true or false positive, classification based on number of haplotypes supported by the reads (basic classification), and a stringent classification that incorporates additional criteria to classify a site (stringent classification). We classified variants per cell as true positives when supported by linked reads from exactly two haplotypes, and as false positives otherwise, except when a third haplotype was supported by only a single read, which was still considered a true positive. Variants supported by just one haplotype were filtered out. We then calculated the proportion of true and false positives per variant across cells, and annotated variants with a ratio of true positives and false positives different from one and zero, respectively, as uncertain. For the stringent classification, some true positive variants of the basic classification were also reclassified as uncertain if they failed to meet one or more quality criteria. Specifically, true positives were labeled uncertain if they had median linked-read haplotype Shannon diversity across cells higher or equal to 0.7, a true positive ratio different from one, or were supported in only one cell. In addition, false positives with haplotype diversity less or equal than 0.75 were also marked as uncertain. We then trained a model on the stringent classification of variants across all donors using *XGBoost*. We performed 5-fold cross-validation to optimize model performance and assess accuracy. The generated probabilities of SMART-PTA error for each site were annotated into the VCFs using *bedtools* (v. 1.9). Variants detected in only a single cell were discarded if their classifier-predicted probability of being an error exceeded 0.5, whereas a stricter threshold of 0.9 was applied to remove variants present in two or more cells. In addition, autosomal variants observed in only one cell were excluded if they exhibited extreme variant allele frequencies (>0.9 or <0.25), in combination with the classifier-based filters.

### Calculation of phylogenetic branch lengths

By incorporating mutation mapping using the *treemut*^49^ R package (v. 1.1) for branch length recalculation in number of mutations, we aimed to reduce the false negative rate and filter out recurrent PTA-induced errors that appear across multiple unrelated branches. We first mapped mutations after tree building and then we remapped the filtered set after applying the classifier. Cells with a proportion of singleton avsnp150 sites higher than 0.25 were removed prior to the second mutation mapping round. Further cell filtering was manually performed when a negative correlation was observed between the cell breadth and branch length likely caused by exogenous contamination.

### Phylogenetic time-calibration analysis

Tree calibration was performed with the command-line application from MEGA^128^ (v.11) running the *RelTime* method with calibration information on the ancestral nodes generating both epithelial and immune cells. We took advantage of the fact that we analyzed cells from two different germ layers: endoderm-derived epithelial (EPCAM+) cells and mesoderm-derived immune (CD45+) cells, whose multipotent common ancestors would have existed no later than gastrulation^129^. We searched for shared mutations on the branches that would identify multipotent ancestral cells, and we only observed a very low (or null) number supporting the early splits further supporting the early divergence. As we have sampled immune and epithelial cells from distal esophageal regions, we can only a single read, which was still considered a true positive. Variants supported by just one haplotype were filtered out. We then calculated the proportion of true and false positives per variant across cells, and annotated variants with a ratio of true positives and false positives different from one and zero, respectively, as uncertain. For the stringent classification, some true positive variants of the basic classification were also reclassified as uncertain if they failed to meet one or more quality criteria. Specifically, true positives were labeled uncertain if they had median linked-read haplotype Shannon diversity across cells higher or equal to 0.7, a true positive ratio different from one, or were supported in only one cell. In addition, false positives with haplotype diversity less or equal than 0.75 were also marked as uncertain. We then trained a model on the stringent classification of variants across all donors using *XGBoost*. We performed 5-fold cross-validation to optimize model performance and assess accuracy. The generated probabilities of SMART-PTA error for each site were annotated into the VCFs using *bedtools* (v. 1.9). Variants detected in only a single cell were discarded if their classifier-predicted probability of being an error exceeded 0.5, whereas a stricter threshold of 0.9 was applied to remove variants present in two or more cells. In addition, autosomal variants observed in only one cell were excluded if they exhibited extreme variant allele frequencies (>0.9 or <0.25), in combination with the classifier-based filters.

### Calculation of phylogenetic branch lengths

By incorporating mutation mapping using the *treemut*^49^ R package (v. 1.1) for branch length recalculation in number of mutations, we aimed to reduce the false negative rate and filter out recurrent PTA-induced errors that appear across multiple unrelated branches. We first mapped mutations after tree building and then we remapped the filtered set after applying the classifier. Cells with a proportion of singleton avsnp150 sites higher than 0.25 were removed prior to the second mutation mapping round. Further cell filtering was manually performed when a negative correlation was observed between the cell breadth and branch length likely caused by exogenous contamination.

### Phylogenetic time-calibration analysis

Tree calibration was performed with the command-line application from MEGA^128^ (v.11) running the *RelTime* method with calibration information on the ancestral nodes generating both epithelial and immune cells. We took advantage of the fact that we analyzed cells from two different germ layers: endoderm-derived epithelial (EPCAM+) cells and mesoderm-derived immune (CD45+) cells, whose multipotent common ancestors would have existed no later than gastrulation^129^. We searched for shared mutations on the branches that would identify multipotent ancestral cells, and we only observed a very low (or null) number supporting the early splits further supporting the early divergence. As we have sampled immune and epithelial cells from distal esophageal regions, we can assume that most likely the most recent common ancestor of all our cells in each patient is the zygote or one of the embryo blastomeres. As such, we created a calibration file to ensure that the age of nodes from these branches with shared mutations was constrained to be less than the donor birth age. For each of these nodes, the patient age plus 9 months (fertilization age) was used as *minTime*, and the patient age as *maxTime*. The *time* parameter for the root node of the tree was set up to the fertilization age.

### Somatic SNV mutation burden

Branch lengths derived from mutation mapping, measured in number of mutations, were added from the root to each of the terminal nodes using the *get_all_distances_to_root* from *castor* R package, and used as a measurement of somatic SNV burden per cell. Resulting counts per cell were corrected by considering per-cell variant calling recall percentages:

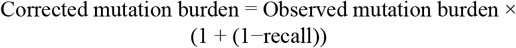

### Phylogenetic clade (clone) count

To identify and quantify clones within somatic phylogenies, we defined a temporal cut-off at birth (0.75 time units from the root) and retained all the nodes branching beyond this point. Using the R packages *dplyr*(v. 1.1.3), *tidyr*(v. 1.3.1) and *treeio*^130^(v.1.14.4), we converted the tree into a data. frame object and selected nodes branching immediately after the cut-off as candidate clones. We then filtered out nested internal nodes to avoid double-counting descendant lineages, retaining only terminal tips and the most basal internal nodes. For each retained basal node, we then enumerated its tip descendants and annotated their correspondent cell types in an array.

### Lineage through time (LTT) analysis

LTT curves were computed for each clone phylogeny of interest using the *ltt*.*plot*.*coords* function from the ape package. For immune versus epithelial LTT comparisons, region- and cell type–specific phylogenies were obtained by pruning non-target tips from the complete donor phylogenies with the *drop*.*tip* function from the *treeio* R package (v. 1.14.4). If fewer than three tips were available for a given cell type and region, the LTT was not computed to avoid artifacts due to low sampling. Proportional lineage counts were calculated for each tree. These proportions were independently fitted for immune and epithelial compartments across donors and regions using a generalized additive model (*method = “gam”*) to capture smooth trends in lineage accumulation.

We computed the γ-statistic^66^ using the *gammaStat* function from the *ape* R package. The corresponding *P* values were obtained from a two-sided normal approximation. We further calculated the γ-statistic and *P* value using the Monte Carlo Constant Rates (MCCR) function implemented in the R package phytools with parameters rho=0.005 and nsim=100. Rho was estimated to be 0.005 as our clones are defined by the epithelial stem cell population, estimated to be ∼5,000 for a biopsy of 1mm^2 131^ while sampling around 50 cells. The presence of thousands of stem cells per 1mm^2^ aligns with the expected number of cells in the basal layer, given an estimated 100,000 cells per 2mm^2^ of a skin biopsy^132^ which has a similar structure to the esophageal epithelium and assuming that it is composed of 25–40 layers. We are assuming similar sampling rates for immune cells as we are likely sampling a higher proportion of the immune population than the epithelial while expecting a lower number of descendants originating from multiple hematopoietic stem cells present in the biopsy. Immune cells are reported to range between 5-15% of the cells in the esophageal epithelium^34,35^.

### Phylogenetic correlations between immune and epithelial cells

For each donor, we calculated a phylogenetic weight matrix using the *one_node_tree_dist* function from *PATH*^19^. Cell types (EPCAM+ and CD45+) were converted into a presence/absence matrix with cell names as rows using the *catMat* function. Percent unmapped reads per cell were included as an additional column to serve as a negative control. Phylogenetic correlation Z scores were then computed using *xcor* (*PATH*) with the phylogenetic weight matrix and the cell metadata matrix as inputs.

### Phylogenetic correlation of anatomic esophageal regions

For each donor, we separated the full phylogeny into two compartment-specific subtrees: one containing only epithelial cells (EPCAM+) and another containing only immune cells (CD45+). For each compartment, we computed a pairwise phylogenetic inverse distance weight matrix using the *inv_tree_dist* function from *PATH*^19^. We then constructed a vector containing the anatomical location of each cell, expressed as the distance (in cm) from the frontal teeth, and calculated phylogenetic correlations Z score between anatomical distance using *xcor*, including the weight matrix previously calculated. For donor eso02, all immune cells were sampled from the same anatomical region, preventing estimation of phylogenetic correlations, which require at least two sampling locations. To estimate variability in the phylogenetic correlation Z score statistic, we generated five tree subtrees per compartment by randomly partitioning the tips and calculated phylogenetic correlations on them, using the standard error of these replicates to generate error bars.

### Epithelial signature analysis

Immune branches were removed from the phylogenetic tree using *treeio* (v.1.14.4) and *tibble* (v3.2.1) R packages, SNVs obtained from Sequoia output were read with *vroom* (v1.6.5) and merged with branch metadata. Divergence times (backwards, from present to past) for branches were used to classify mutations into three groups: early, middle, and late, respective to donor’s life periods. The branches were assigned to the three different categories based on branch type and the branch terminal node age in divergence time: internal branches with ≥40 years old (early period of donor’s life), internal branches with <40 years old (middle period of donor’s life), and external branches with terminal node exactly matching the sampling point, zero years old). Importantly, mutations can occur at any point in the branch and even earlier under incomplete sampling, any time before the divergence of the most recent common cell carrying it. For that reason, we calculated the branch midpoint age in forward time by subtracting half of the branch length from the branch terminal node age in forward time. Finally, trinucleotide mutation contexts for mutations assigned to each branch category were extracted relative to the GRCh38 reference genome and converted into the 96 mutational profiles using the *MutationalPatterns* function *mut*.*to*.*sigs*.*input*. To further decompose these trinucleotide profiles, we used Sigprofiler with the COSMIC (hg38, v3.4) SBS reference signature set, to fit to aging (SBS1, SBS5, SBS40a), tobacco/alcohol exposure (SBS4, SBS92, SBS16), and clear cell renal cell carcinoma signatures (SBS40b and SBS40c, these serving as a negative control).

### Proportion of mutated vs wild-type cells

Somatic SNVs in 83 genes previously analyzed in phenotypically normal esophagus studies (**Supplementary Table 5**) were annotated with Combined Annotation Dependent Depletion (CADD) PHRED scores^133^ in R. From the original CADD database (hg38, v1.7), we extracted site-specific SNV tables for each gene using a bash script. Only exonic SNVs annotated as synonymous or non-synonymous by *ANNOVAR* were kept.

Twenty-eight genes that were shown to be under positive selection from both studies^3,4^ were used for driver mutation analysis (**Supplementary Table 5**). The branch identifier for each mutation on these driver genes (encoded as zeros and ones according to the tree topology) was then used to calculate the number of mutations per cell. Cells with zero SNVs were annotated as wild type.

### dN/dS analysis

The ratio of synonymous to non-synonymous variants was computed using the R package *dndscv*^89^ (v. 0.0.1.0) with hg38 covariates. Per-patient gene dN/dS rates were obtained by supplying a *gene_list* parameter with *TTN, FAT1* and *TP53*. Confidence intervals for the genes were calculated using *dndscv* the *geneci* function with the same *gene_list*. For all donors, the presence-absence matrix of gene mutations was randomly downsampled to 608 cells (minimum cell number across donors), and mutations with all-zero entries were removed. This ensured that differences in sampling rates, which can produce distinct VAF profiles and thereby influence dN/dS estimates, did not confound comparisons across donors. Each donor’s mutations were included only once.

### Loss of heterozygosity (LOH) calling

Germline variants were phased using *SHAPEIT5* (v5.1.0). For each donor, variants with minor allele frequency (MAF) ≥ 0.1% were phased in predefined chunks, corresponding to large genomic regions (∼25 cM) provided with the program resources. Each chunk was phased independently using the *phase_common_static* function with a genetic map and a high-coverage 1,000 Genomes reference panel. Phased chunks were subsequently ligated using *ligate_static* to generate chromosome-wide haplotypes, which were indexed with *BCFtools* (v1.15.0). LOH events were detected using the *HMMCopy* R package, specifically the function *HMMSegment* run with non-default parameters to improve sensitivity to allelic imbalance. Specifically, mu was set to the 50th and 99th quantiles of the observed VAF distribution, while kappa and eta were adjusted to (0.7, 0.3) × 1000 and (50, 5) × 10,000, respectively.

### CNV calling

Copy number variants (CNVs) were called using *Ginkgo*^134^. For each single cell, BAM files were filtered to retain reads with mapping quality ≥ 40 using *samtools* (v1.21), and converted to BED format with *bedtools* (v2.31.0). The resulting BED files were used as input to the analyze.sh script provided with the software. Copy number profiles were inferred using default parameters, including GC correction and segmentation, except for the reference genome (hg38 instead of hg19). Required files for running *Ginkgo* with hg38 were constructed guided by the structure of the precalculated files for hg19. CNVs were called at both 500-kb and 50-kb bin sizes.

### Targeted deep sequencing of driver genes

Targeted deep sequencing of a driver gene panel of *NOTCH1, NOTCH2, NOTCH3, TP53, FAT1, PPM1D*, and *ZFP36L2* was performed according to manufacturer’s protocol using a custom bait panel of probes (xGen Custom Hybridization Capture Panel, IDT). The panel was designed to capture exons of driver genes (**Supplementary Table 7**). All genes were designed with 2× tiling, with the exception of *NOTCH2* and *FAT1*, which had 1× tiling due to their increased coverage in the scWGS libraries. Final Illumina libraries prepared from PTA-amplified products were pooled for hybridization capture. After hybridization capture and library amplification, enriched fragments were sequenced using paired-end 150 bp on a Novaseq6000.

For each cell, adapter sequences were first removed using *CutAdapt*. Reads were then aligned to the reference genome (hg38) using *BWA MEM*, and the resulting alignments were sorted with *Picard SortSam*. Duplicate reads were identified and marked during merging using Picard *MergeWithMarkDuplicates*. Finally, base quality scores were recalibrated in two steps with *GATK*. Initial variant calling was performed per donor using *GATK HaplotypeCaller* in GVCF mode. The resulting GVCF files were then jointly genotyped using *GATK GenotypeGVCFs* to produce a single VCF.

### Single-cell targeted genotype-to-phenotype assay

Single-cell targeted genotype-to-phenotype assay (scG2P^84^) was performed according to the authors’ methods. Briefly, a DNA targeted panel was designed to capture hotspot mutations along six driver genes with high mutation frequencies in normal esophageal epithelium. A targeted RNA panel was designed to capture 58 transcripts encompassing epithelial cell types, epithelial differentiation, and cell cycling. Four biopsies from a donor were dissociated enzymatically and sorted for EPCAM+ populations to be used as input. 100,000 EPCAM+ cells were loaded onto the Mission Bio Tapestri device, and single-cell encapsulation and library preparation was performed according to the authors’ methods. Final libraries were sequenced on NovaSeq6000 with paired-end 150 bp. DNA FASTQs were input to the Mission Bio Mosaic pipeline for adapter trimming and cell calling. Variant calling was performed using *GATK HaplotypeCaller*; the set of GVCFs were merged into a Mission Bio object. Genotypes were assigned to single cells, then filtered at the variant level for the quality of the variant and how frequently it is genotyped, followed by filtering at the cell level. Genotyping cells required the following criteria: total read counts >= 10 per cell, alternate allele count >= 3, and genotyping quality score (*GATK*) >30. We further removed variants where genotyping calls were present in less than 50% of cells and removed cells with <50% informative genotypes. Genotypes were converted to wild-type, heterozygous, and homozygous. Status of loss of heterozygosity of *NOTCH1* was determined using the five SNPs detected in intronic regions and verified on GnomAD. RNA FASTQs were processed using a custom script (https://github.com/Theob0t/PRIMR). Cell by transcript matrices were generated and analyzed using *Seurat* (v. 4.0.0). Cells were filtered for a minimum of 50 reads per cell. Count matrices were normalized using Center-Log Ratio, followed by dimension reduction and clustering. Cell cycling and differentiation scores were calculated using *AddModuleScore* function using gene lists previously described.

### Esophagus donor sample

For samples eso02-04, donors were recruited from among adult patients who were scheduled for elective upper endoscopy for clinical indications at Columbia University Irving Medical Center. Signed informed consent was obtained prior to or on the day of endoscopy from all patients. The protocol was approved by the Columbia University institutional review board (AAAS4603).

Sample eso01 was from enrolled patients who underwent therapeutic or diagnostic endoscopy for upper gastrointestinal symptoms at Kyoto University Hospital. Informed consent was obtained. A history of heavy alcohol drinking (HIGH indicated by ≥396 g alcohol per week) and tobacco smoking (HIGH indicated by ≥30 pack-years) was reported in this individual, who was considered to have positive lifestyle ESCC risks. Biomaterials, including esophageal tissues, were newly collected from this individual using endoscopic biopsy, according to procedures approved by the Internal Review Board at Kyoto University (G0645).

## Supplementary Figures

**Supplementary Figure 1:**
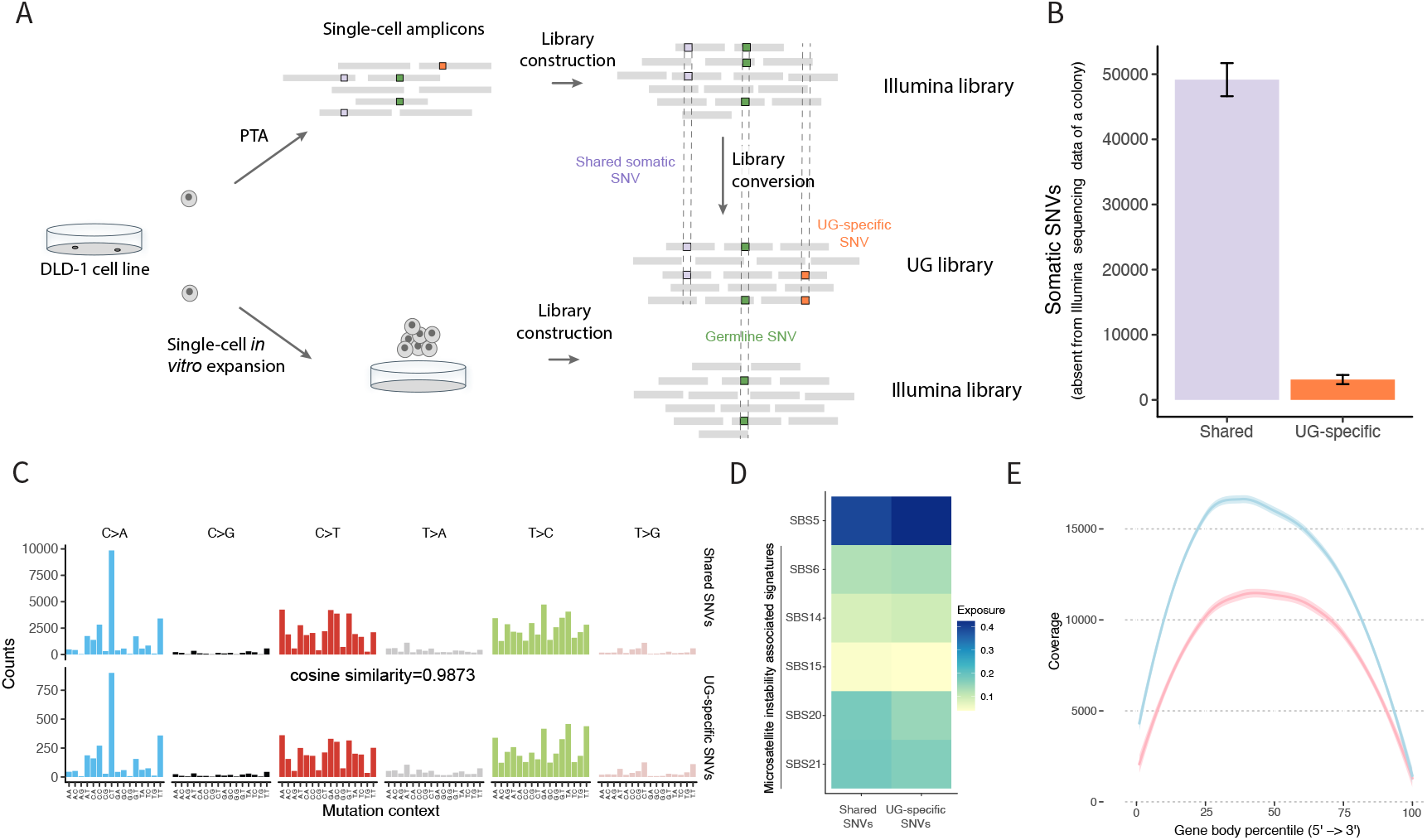
SMART-PTA quality control metrics. **A**, Schematic of library preparation to compare Illumina to Ultima Genomics (UG) sequencing for PTA. Illumina sequencing data from a DLD-1 colony (bottom) were downloaded from SRA to use as a germline control^127^. **B**, Mean number of SNVs called across 4 UG libraries (F1, F2, H1 and H2) that are also detected in the matched Illumina library (shared), or that are present only in the UG-converted libraries (UG-specific). Only cells from clones F and H were selected as they represent independent lineages that evolved for the same duration and had >15× coverage for both Illumina and UG libraries. Error bars represent the standard deviation. **C**, Mutation signature analysis of the trinucleotide context of SNVs identified in both Illumina and Ultima libraries (shared) and SNVs that were only called from Ultima libraries (UG-specific). Cosine similarity between these two SNV sets is 0.9873. **D**, Heatmap showing single base substitution mutation signatures (SBS) identified from both Illumina and UG libraries (shared SNVs) and from UG libraries (UG-specific SNVs). SBS signatures were obtained from the COSMIC database. The scale bar represents the level of signature enrichment. SBS5 is a ubiquitous, clock-like signature, while SBS6, SBS14, SBS15, SBS20 and SBS21 are microsatellite instability-associated signatures that are expected to be observed in DLD-1 cells, which have microsatellite instability. **E**, Average number of cDNA reads (coverage) across the gene body (represented as gene body percentile from the 5’ to 3’ ends) using standard-PTA (pink) versus SMART-PTA (blue).

**Supplementary Figure 2:**
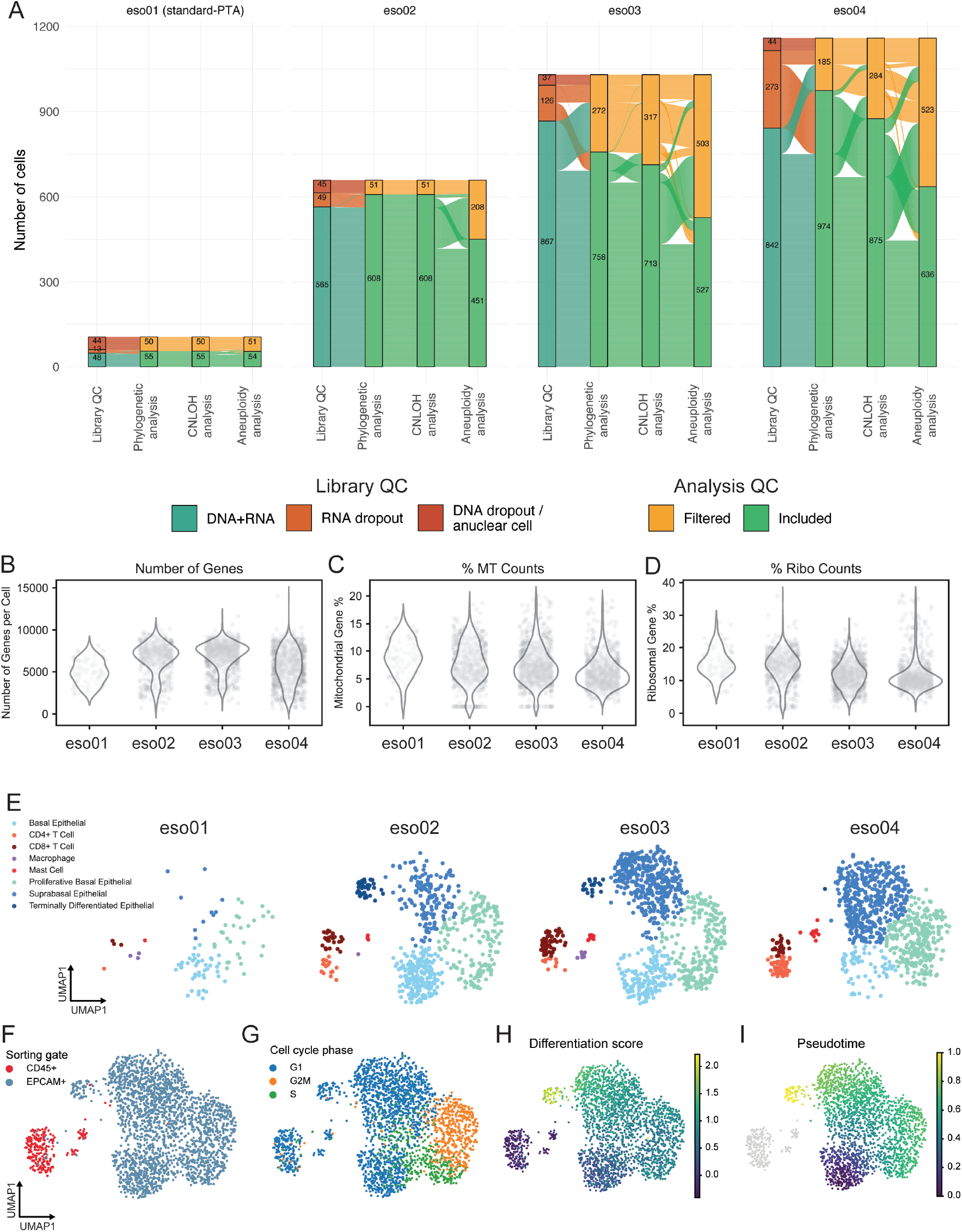
Single-cell RNA profile metrics and cell type analysis across donors. **A**, Sankey plot of loss of DNA and RNA libraries during standard-PTA (eso01) and SMART-PTA (eso02, eso03, eso04) library preparation and number of cells filtered out during phylogenetic, CNLOH, and aneuploidy analyses. CNLOH = copy neutral loss of heterozygosity. Cells with <200K read counts were considered RNA dropouts. For phylogenetic reconstruction, we only considered high quality cells with breadth > 50%, depth of coverage >= 3x, unmapped reads below 5% and MAD lower than 0.5. Additionally, cells with >25% singletons overlapping with dbSNP and those with low genome breadth and long external branch lengths were removed from the tree backbone after mutation mapping, to minimize potential risk of exogenous DNA contamination. From the remaining cells, those that showed noisy heterozygous germline site profiles were excluded from CNLOH analysis. For aneuploidy analysis, we only included cells in the phylogeny with high coverage uniformity (MAD<0.2). **B**, Quality control metrics for RNA libraries generated from each of the four donors, showing the number of genes per cell. **C**, Percentage of mitochondrial (MT) gene reads per cell displayed for each donor. **D**, Percentage of ribosomal RNA gene reads per cell displayed for each donor. **E**, RNA expression UMAP for each donor (eso01 = 92 cells; eso02 = 610 cells; eso03 = 904 cells; eso04 = 886 cells). Cells are colored by annotated cell type, including immune cells and epithelial cells. **F**, UMAP showing sorting gate assignment into CD45+ and EPCAM+ populations from FACS cell sorting for all donors (n = 2,491 total cells). **G**, UMAP showing cell cycle phase (G1, G2M and S) from the single-cell RNA data for all donors (n = 2,491 total cells). **H**, UMAP showing differentiation score for the EPCAM+ population from the single-cell RNA data for all donors (n = 2,491 total cells).

**Supplementary Figure 3:**
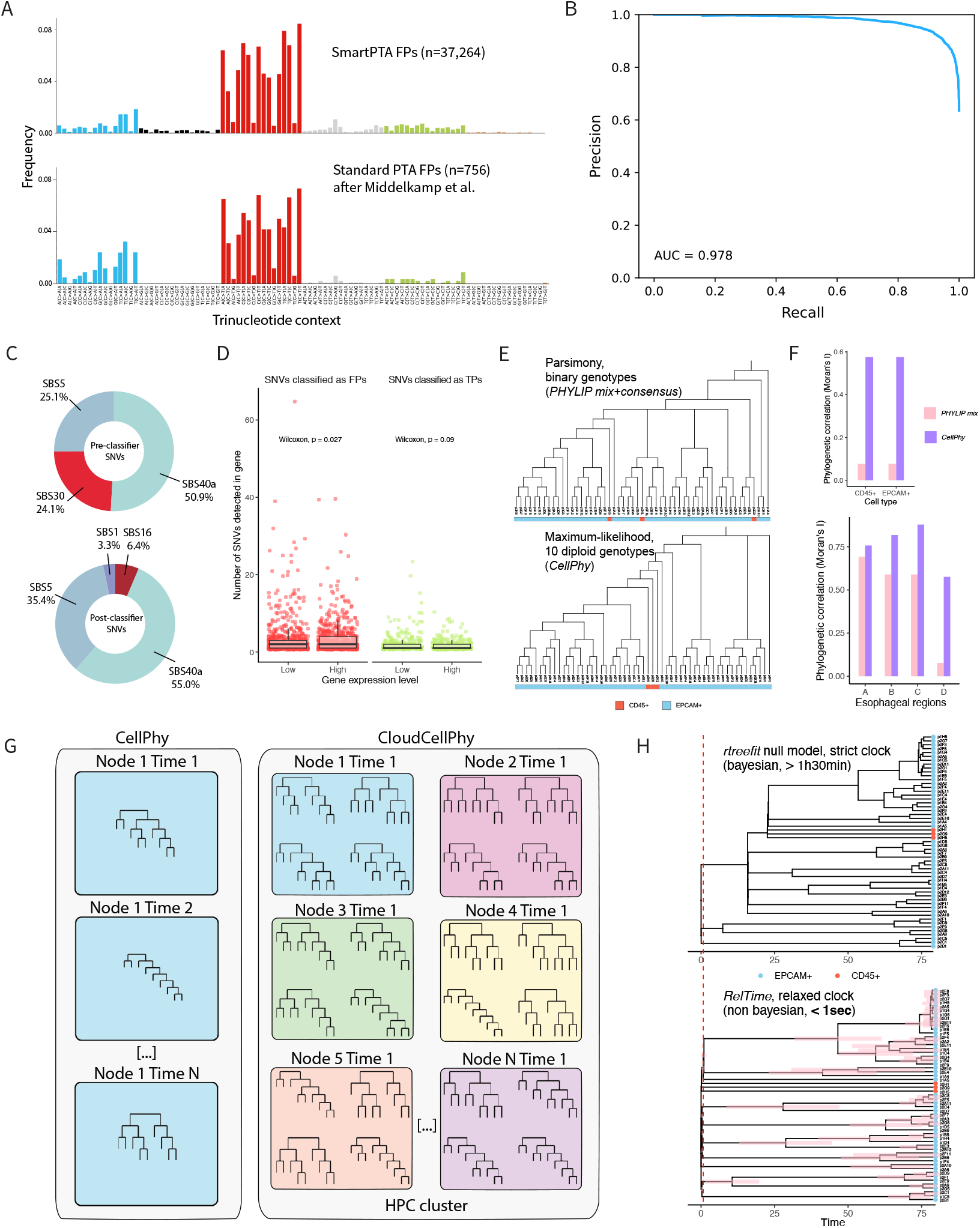
Overview of classifier performance, SMART-PTA error profiles, RNA-editing filtering, and phylogenetic inference and calibration approaches. **A**, Top, Trinucleotide context of the false positives (FPs) used to train the classifier. Bottom, Approximated trinucleotide context of the SNVs classified as FPs in Middelkamp et al.^43^, adapted from the figure in the publication (Middelkamp et al., *Cell Genomics*, 2023). **B**, Precision-Recall curve of the SMART-PTA classifier trained on 37,264 FPs and 21,349 true positives (TPs) obtained through a linked-read analysis (Methods). **C**, Signatures of the SNVs before (top) and after (bottom) applying the classifier (error probability > 0.5 for FPs and <= 0.5 for TPs) across all donors. SBS30, the signature previously reported as more similar to the PTA error profile^43^, present at 24.1% of the variants in prefiltering, is not detected after applying the classifier. Remaining signatures are explored in more detail in Fig. 4C and Supplementary Fig. 6D [SBS1 (ubiquitous clock-like), SBS5 (ubiquitous clock-like), SBS16 (aldehyde exposure-related) and SBS40a (ubiquitous clock-like)]. **D**, Number of SNVs detected per high or low expression gene (including exonic, intronic, UTR and up/downstream mutations) for the FP SNV set (left) and TP SNV set (right). Genes were classified as showing low or high expression level based on the normalized mean expression level in the RNA analysis with the threshold selected to create two groups with a similar number of genes using the *ntile* function from the *dplyr* R package. Box plots represent the median, bottom and upper quartiles; whiskers correspond to 1.5 times the interquartile range. Two-sided Wilcoxon test. **E**, Maximum parsimony phylogeny (built with *PHYLIP mix* followed by *consensus*, top) and maximum likelihood phylogeny (built with *CellPhy*, bottom) from donor eso01 after removal of poor-quality cells (Methods). In the CellPhy tree, the immune cells cluster under a direct common ancestor (one-node away distance among them) rather than being distributed across multiple clades (multiple node away distance among them), as observed in the maximum parsimony tree. This suggests that the *CellPhy* tree is more accurate. **F**, Phylogenetic correlation (Moran’s I) of esophageal regions (A, B, C, D) and cell types (CD45+, EPCAM+) for a phylogeny of 60 eso02 cells. Fifteen cells per biopsy region were randomly selected, and their somatic variant sites were downsampled from the VCF to reconstruct phylogenies. Trees were inferred either with *Phylip mix* followed by *consensus* (runtime: 130,385s) or with *CellPhy* under the GT10+FO+E model (runtime: 95,951s). The phylogenetic distance matrix used for phylogenetic correlation estimation was based on node distances, since parsimony does not provide branch lengths. Higher phylogenetic correlation indicates less phenotypic splitting on the tree, which is consistent with biological expectations. Region D, CD45+ and EPCAM+ phylogenetic correlations for the Phylip tree are not significant, indicating completely random segregation on the tree. **G**, Schematic of the computational parallelization implemented by *CloudCellPhy*. Left, *CellPhy* in default mode sequentially performs maximum likelihood tree searches. Right, *CloudCellPhy* uses multiple nodes, with multiple processes per individual node, within a high-performance computing (HPC) cluster to simultaneously perform maximum likelihood tree searches. **H**, Time calibrated phylogenies of eso01. Calibration performed with a bayesian method (*rtreefit*^49^, (top) or with a non-bayesian method (*RelTime*^50^). Although it is possible to run *rtreefit* specifying different rates for different branches, potentially providing a more consistent estimate of divergence time for the MRCA of epithelial and immune cells (which should be 2-3 weeks after fertilization, before birth), the method is time consuming and doesn’t escalate well with number of cells. In contrast, *RelTime* estimates divergence times that are broadly consistent with developmental timings for the different germ layers when only constraining the age of the root to the donor age plus the gestation period.

**Supplementary Figure 4:**
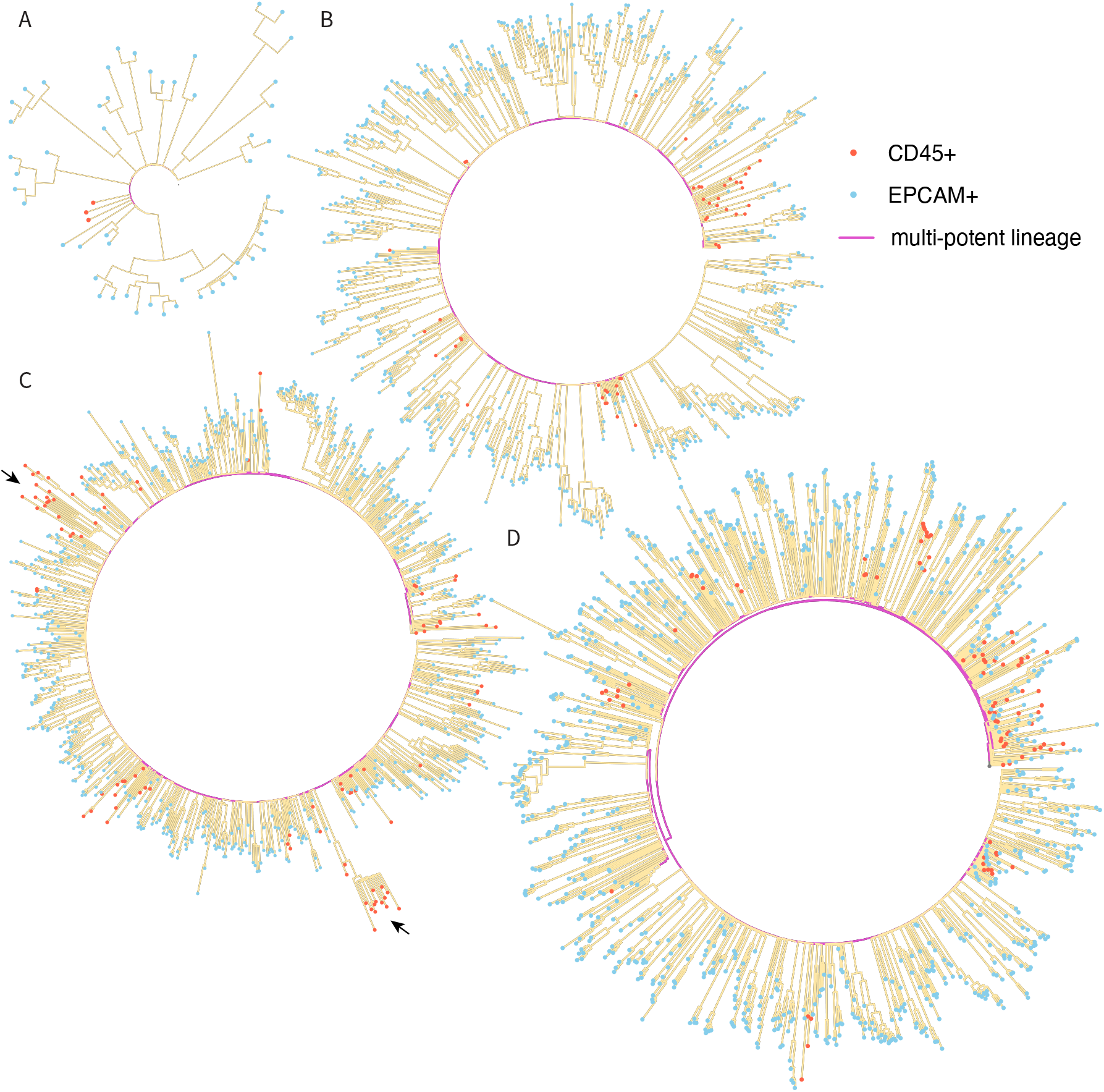
Phylograms displaying epithelial and immune lineages for each of the four donors. **A**, Branch lengths represent genetic distance in number of mutations. Multi-potent lineages that lead to both EPCAM+ and CD45+ cell descendants are shown in magenta. The remaining lineages whose descendants consist of a single cell type (EPCAM^+^ or CD45^+^) are shown in light yellow. Donor eso01 (n = 55 cells, maximum distance from root to terminal node = 3,888 mutations). **B**, Donor eso02 (n = 608 cells, maximum distance from root to terminal node = 3,987 mutations). **C**, Donor eso03 (n = 758 cells, maximum distance from root to terminal node = 3,081 mutations). Arrows indicate immune clones with high mutation burden. **D**, Donor eso04 (n = 974 cells, maximum distance from root to terminal node = 3,928 mutations).

**Supplementary Figure 5:**
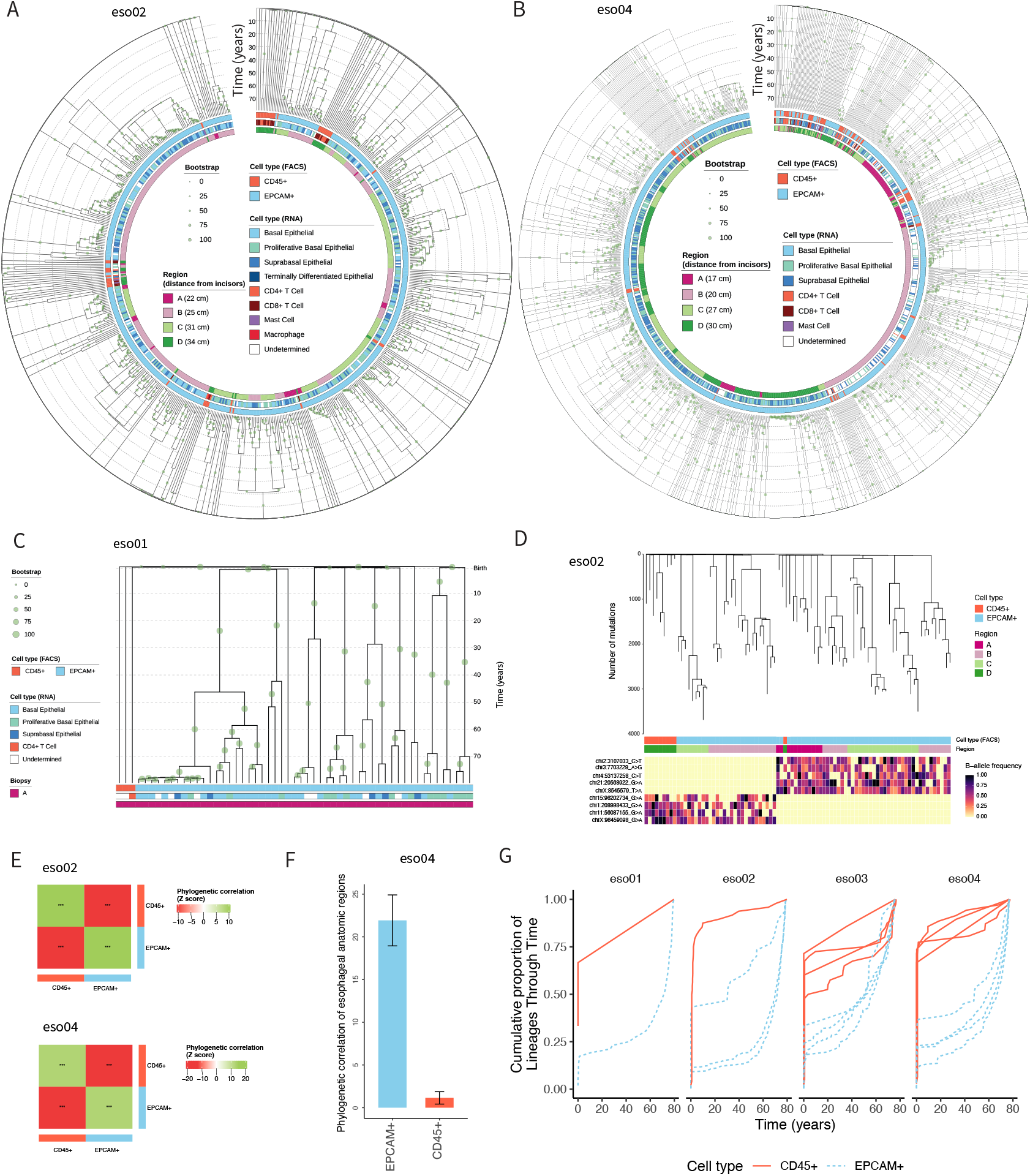
Single-cell phylogenies and lineage analysis of immune and epithelial cells in the aging esophagus. **A**, Single-cell phylogeny of n = 608 total cells built from SNVs obtained from SMART-PTA application to esophagus samples from donor eso02. Each leaf of the tree represents a single cell, annotated as epithelial (EPCAM+) or immune (CD45+) cell based on flow cytometry data, or according to cell type determined from the RNA profiles. Regions corresponding to the sampling scheme in Fig. 2A are indicated and annotated for each cell. Branch lengths represent time in years. Bootstrapping values are indicated by green circles. Because of the large size of the eso04 phylogeny, we performed only 10 bootstrap replicates instead of 100 and scaled the results. **B**, Same as (A) for donor eso04 (n = 974 total cells). **C**, Same as (A) for donor eso01 (n = 61 total cells), presented as a linear rather than circular phylogeny due to the lower number of cells. Note that cells from this donor underwent sequencing with the standard-PTA protocol. **D**, Subset phylogeny for donor eso02 showing two nodes for which there are both immune and epithelial descendant cells. Branch lengths represent the number of mutations. Cell types (CD45+ immune or EPCAM+ epithelial), sampling region and shared embryonic somatic variants are displayed. **E**, Phylogenetic correlation Z score heatmap of CD45+ (immune) or EPCAM+ (epithelial) cells for donor eso02 (top) and donor eso04 (bottom). Positive correlation indicates that the cell type phenotype is shared by highly related cells. Negative correlation indicates that the cell type phenotype is absent from highly related cells. *** One-sided *P* < 0.05. Phylogenetic correlations were calculated using node distance to avoid circularity, as patristic distances were estimated using epithelial-immune divergence assumptions. Raw phylogenetic correlations are transformed into Z scores representing permutation tests using the *PATH* framework^19^. **F**, Mean phylogenetic correlation of esophageal regions for the EPCAM+ cells and CD45+ cells for donor eso04, indicating the degree to which phylogenetically related cells (immune versus epithelial cells) are dispersed in different regions of the esophagus. Phylogenetic correlation Z score of the distance of the biopsy from the incisors is calculated for EPCAM+ or CD45+ cells, respectively. Raw phylogenetic correlations are transformed into Z scores representing permutation tests using the *PATH* framework^19^. Error bars represent the standard error of 5 subtree replicates. This analysis was not carried out on eso01 or eso02 as only one sampled region per donor included immune cells. **G**, Generalized additive model (GAM) fit of the cumulative proportion of lineages through time (years) across non-overlapping clones of exclusively epithelial and immune cells at birth for each donor. Each line represents an esophageal sampling region.

**Supplementary Figure 6:**
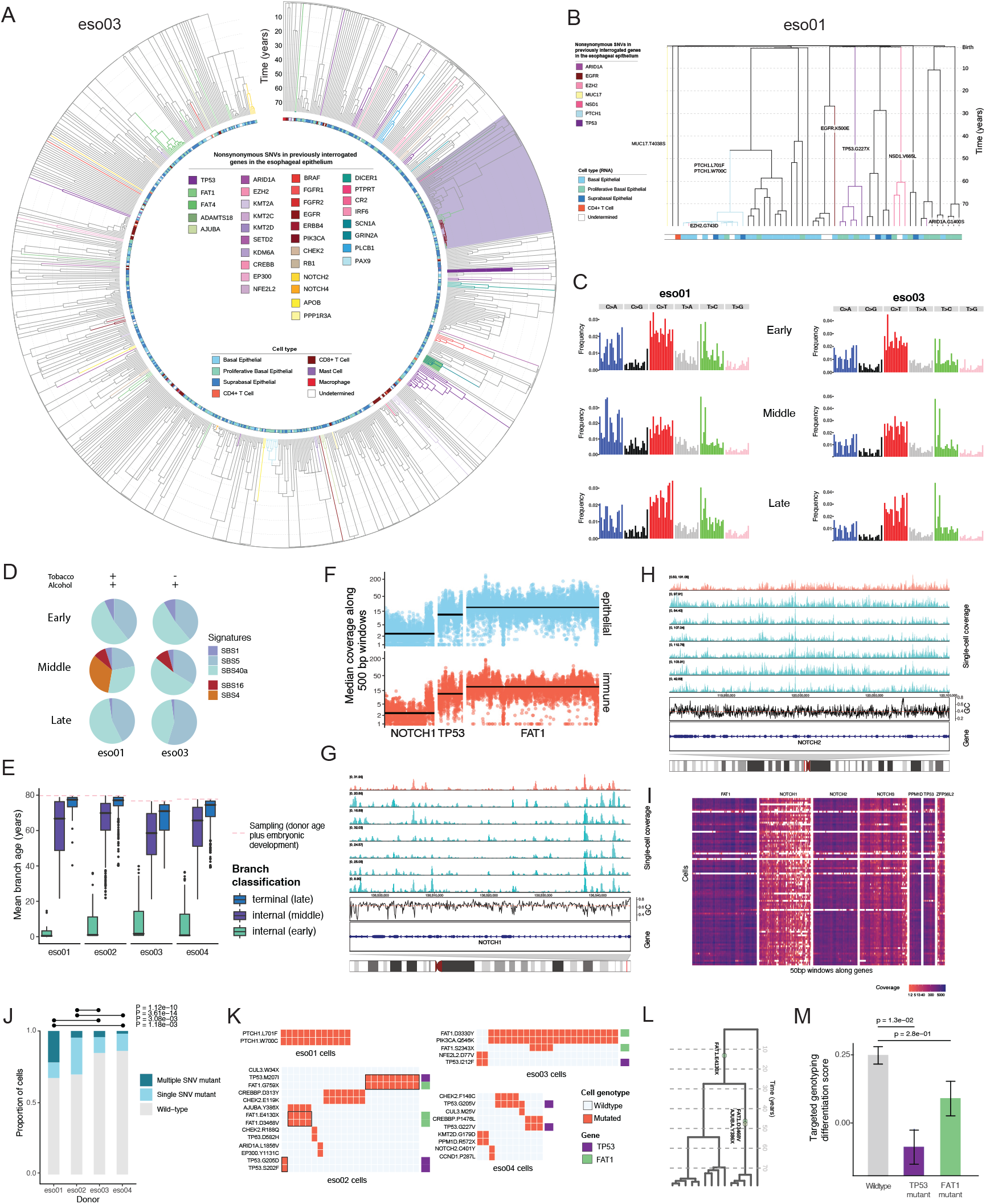
Driver mutations and signatures in the aging human esophagus. **A**, Single-cell phylogeny of n = 758 cells built from SNVs obtained from SMART-PTA application to esophagus samples from donor eso03 with no history of smoking but history of alcohol use. Each leaf of the tree represents a single cell, with cell type annotations displayed. Clones harboring nonsynonymous mutations in any of 83 genes (Supplementary Table 5) previously analyzed in aging esophagus tissue^3,4^ are indicated in different coloring of the branches. Branches with two mapped mutations are further shaded with the gene color of the second mutation. The clone labeled in purple corresponds to Clone B from Fig. 3F. **B**, Single-cell phylogeny of n = 61 cells built from SNVs obtained from standard-PTA+RNA for donor eso01 with history of smoking and alcohol use, presented as a linear rather than circular phylogeny due to the lower number of cells. Annotations are the same as in (A). **C**, Analysis of contributions of trinucleotide mutation signatures across age for eso01(left) and eso03 (right) across lifespan determined from phylogenetic branches (Early – including branches whose end node is older than 40 years, covering a time frame from embryonic development to adulthood; Middle – including branches where the end node is less than 40 years old but terminates in an internal node; Late - terminal branches whose end node is the age of the donor; Methods). Trinucleotide spectra are shown for each donor across the three time points. **D**, Percentage of contribution of four signatures [SBS1 (ubiquitous clock-like), SBS5 (ubiquitous clock-like), SBS16 (aldehyde exposure-related) and SBS40a (ubiquitous clock-like)] are shown for each donor across the three time points. Single-base substitution (SBS) signatures were obtained by analyzing the scWGA data and aligning to COSMIC signatures. **E**, Mean age for different branch classes across all donors. Internal branches whose end diverged earlier than 40 years of donor’s life (including embryonic development) are classified as early. Internal branches whose end diverged after 40 years and prior to sampling, are classified as middle Terminal branch ends match terminal nodes that have been sampled at the same time within each donor and are classified as late. Y axis represents the mean branch age, calculated as the age of the branch end minus half of the branch length. **F**, Average SMART-PTA coverage of *NOTCH1* (3.43×), *TP53* (15.15×) and *FAT1* (23.75×) along 500-bp windows in eso02. Average genome-wide coverage across all donors is 14.8x. **G**, Average coverage along 100-bp windows on the *NOTCH1* gene for seven single cells, epithelial (blue) and immune (orange) showing non uniform whole-genome amplification. The position of the gene is indicated in the chromosome 9 ideogram. Average GC content per window is displayed under the read count histograms of the single cells. **H**, Average coverage along 100-bp windows on the *NOTCH2* gene for seven single cells, epithelial (blue) and immune (orange). The position of the gene is indicated in the chromosome 1 ideogram. Average GC content per window is displayed under the read count histograms of the single cells. **I**, Coverage along 50-bp windows for exonic regions of the genes included in a targeted panel (Supplementary Table 7) applied to PTA libraries generated from donor eso01 (Methods). Driver genes are labeled on top and each row represents a single cell. Duplicate-marked reads were retained in the coverage calculation. **J**, Proportion of mutant cells with variants in any of the 28 esophageal driver genes (Supplementary Table 5) previously reported to be positively selected in the esophageal epithelium^3,4^ across donors. The plot compares the proportion of cells that carry single or multiple synonymous SNVs or that are wild type (no detected synonymous SNVs) across donors. Significant differences in proportion of mutant cells between donors are shown with *P* values. One-sided 𝒳^2^ test or Fisher if <5 counts). **K**, Plot of all multi-mutant single cells with nonsynonymous variants in any of the 28 esophageal driver genes (Supplementary Table 5) previously reported to be positively selected in the esophageal epithelium^3,4^ that are present across cells for all donors, where cells in the same clone are adjacent. *TP53* double mutant, *FAT1* double mutant and *TP53* + *FAT1* double mutant cells are outlined in black. **L**, Zoomed-in region of the eso02 phylogeny showing multi-mutant clones harboring sequentially acquired driver mutations over time representing the donor’s age in years. A clone with a *FAT1p*.*E4130X* mutation split into subclones, where one acquired an *ADAM29p*.*R311C* mutation and the other acquired an additional *FAT1p*.*D3468V* mutation together with an *AJUBAp*.*Y386X* mutation. **M**, Differentiation scores of *TP53*- and *FAT1*-mutant cells obtained from aged esophagus samples and sequenced with targeted single-cell DNA-seq and matched RNA panel^84^. Error bars represent the standard error of the mean (SEM).

**Supplementary Figure 7:**
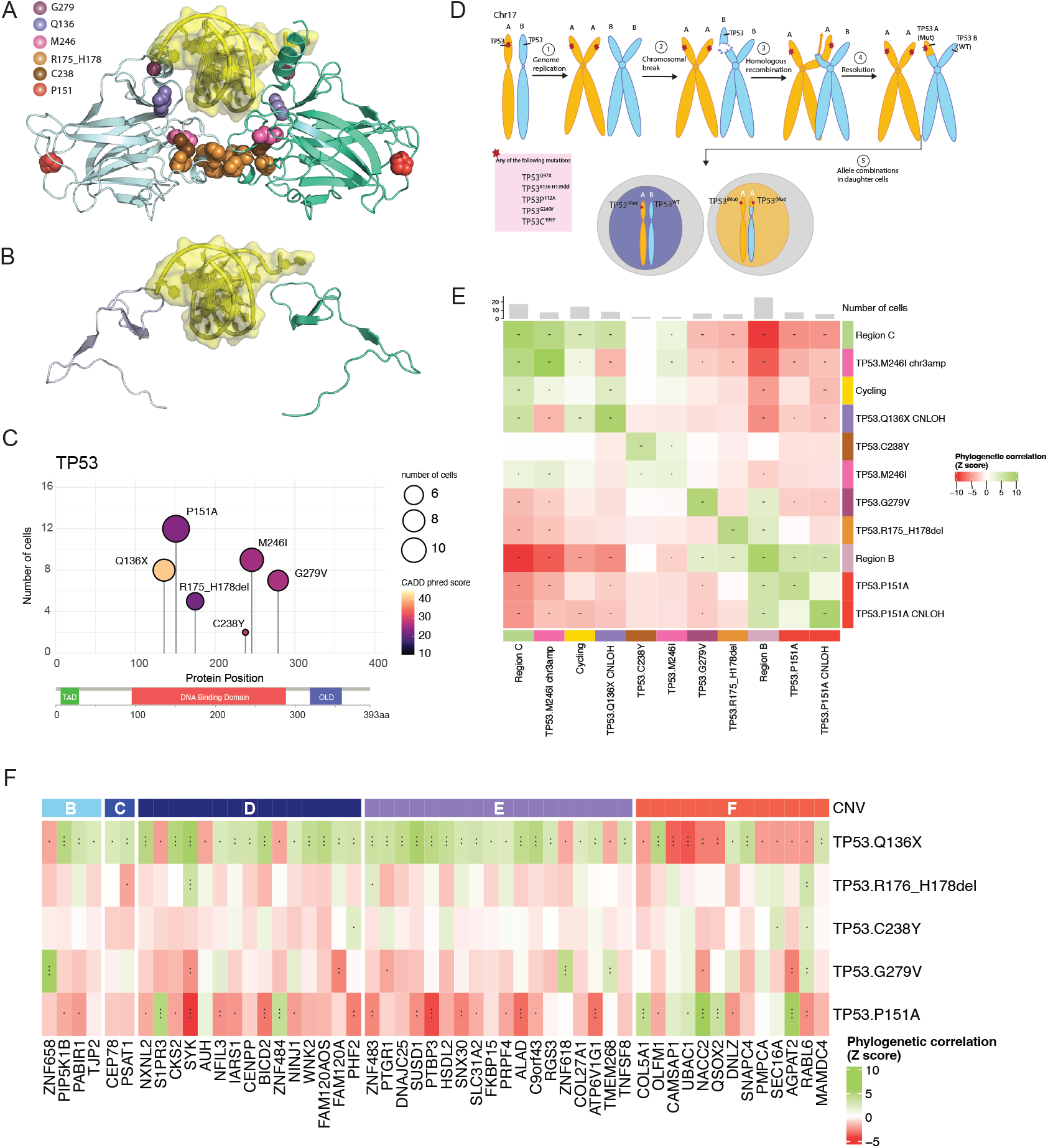
Landscape and phenotypic analysis of TP53-mutant clones with high functional impact prediction scores in eso02. **A**, Illustration of p53 structure^135^ (PDB 2ADY) annotated with the amino acid residues affected by SNVs in different colors. The protein is depicted as a dimer bound to DNA (yellow), in which the impacted amino acids are colored in both subunits. Illustration was prepared using PyMOL. The structure encompasses the whole DNA binding domain: G279 contacts DNA; Q136 is truncated; M246 and R175-H178 are at the dimerization interface, C238Y contacts DNA; disruption to P151 (proline residue) may impact folding. **B**, Illustration of a p53 aberrant structure^135^ (PDB 2ADY) depicting the effect of a truncating variant on residue 136 where originally there was a glutamine (Q136X). The DNA and dimerization interfaces are completely lost. Illustration was prepared using PyMOL. **C**, Illustration of p53 domains with annotations from UniProt (TAD = transactivation domain, OLD = oligomerization domain) for *TP53*-mutant clones identified in donor eso02. CADD PHRED scores and clone cell number are indicated. All variants fall within the DNA binding domain. **D**, Schematic of somatic CNLOH at chromosome 17p, illustrating how after DNA damage, the homologous undamaged chromosome is used as a template for repair. A and B refer to parental haplotype at the terminal end of chromosome 17p covering the *TP53* locus. **E**, Phylogenetic correlation Z score heatmap of indicated *TP53*-mutant cells, proliferation phenotype (S phase and G2M scores) and esophageal biopsy regions for donor eso02. The number of cells in grey is shown at the top. *TP53p*.*Q136X* and *TP53p*.*M246I* mutant clones are exclusively found in the C region. *** = *P* value < 0.001; ** = *P* value < 0.01 * = *P* value < 0.05. One-sided standard normal distribution. ns, not significant. **F**, Phylogenetic correlation Z score heatmap of indicated *TP53*-mutant cells expression of genes along chromosome 9q. The five annotated CNV regions correspond to the identified CNVs found in *TP53*.*pQ136X* mutant cells shown in Fig. 6E.

**Supplementary Figure 8:**
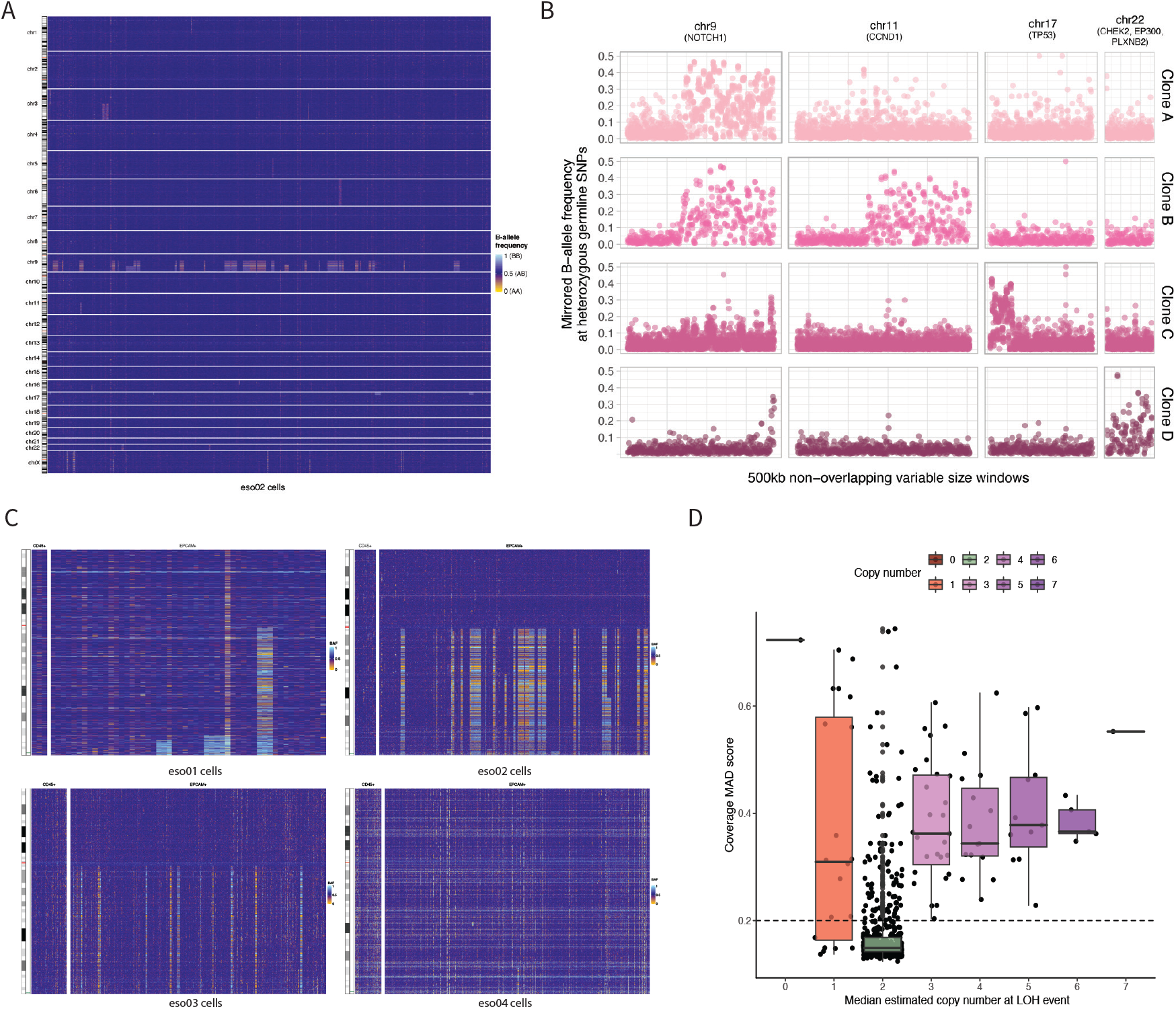
Copy-neutral loss of heterozygosity at chromosome 9q impacts the *NOTCH1* locus and occurs repeatedly. **A**, Genome-wide CNLOH map for all cells from donor eso02. **B**, Single cells from four distinct clones carrying CNLOH events from donor eso02, as indicated by the phased B-allele frequency (BAF) of germline heterozygous sites. Clone A (n = 5 cells) shows CNLOH at chromosome 9q. Clone B (n = 2 cells) harbors two CNLOH events, at chromosome 9q and 11q. Clone C (n = 8 cells) has CNLOH at chromosome 17p. Clone D (n = 2 cells) show CNLOH of chromosome 22. **C**, CNLOH (phased alternate-allele frequency) map of chromosome 9 for all immune (CD45+) and epithelial (EPCAM+) cells from donor eso01 (top left), eso02 (top right), eso03 (bottom left) and eso04 (bottom right). *NOTCH1* is located towards the telomeric end of the q arm of chromosome 9 (centromere in red on the chromosome, *NOTCH1* location indicated in blue to the right of the chromosome). **D**, Mean absolute deviation (MAD) coverage score for cells according to estimated copy number at chr9q arm or LOH region for donor eso02. Grey dashed line indicates a MAD score of 0.2. To ensure confident copy-number state assignment within the chr9q LOH region, we used this as a quality threshold such that cells with a genome MAD score >0.2, indicating higher technical noise, were masked on the CNLOH plot in Fig. 7A. Cells estimated to have higher ploidy (copy number >2) tended to have higher MAD scores, suggesting that this is technical noise. Box plots represent the median, bottom and upper quartiles; whiskers correspond to 1.5 times the interquartile range.

**Supplementary Figure 9:**
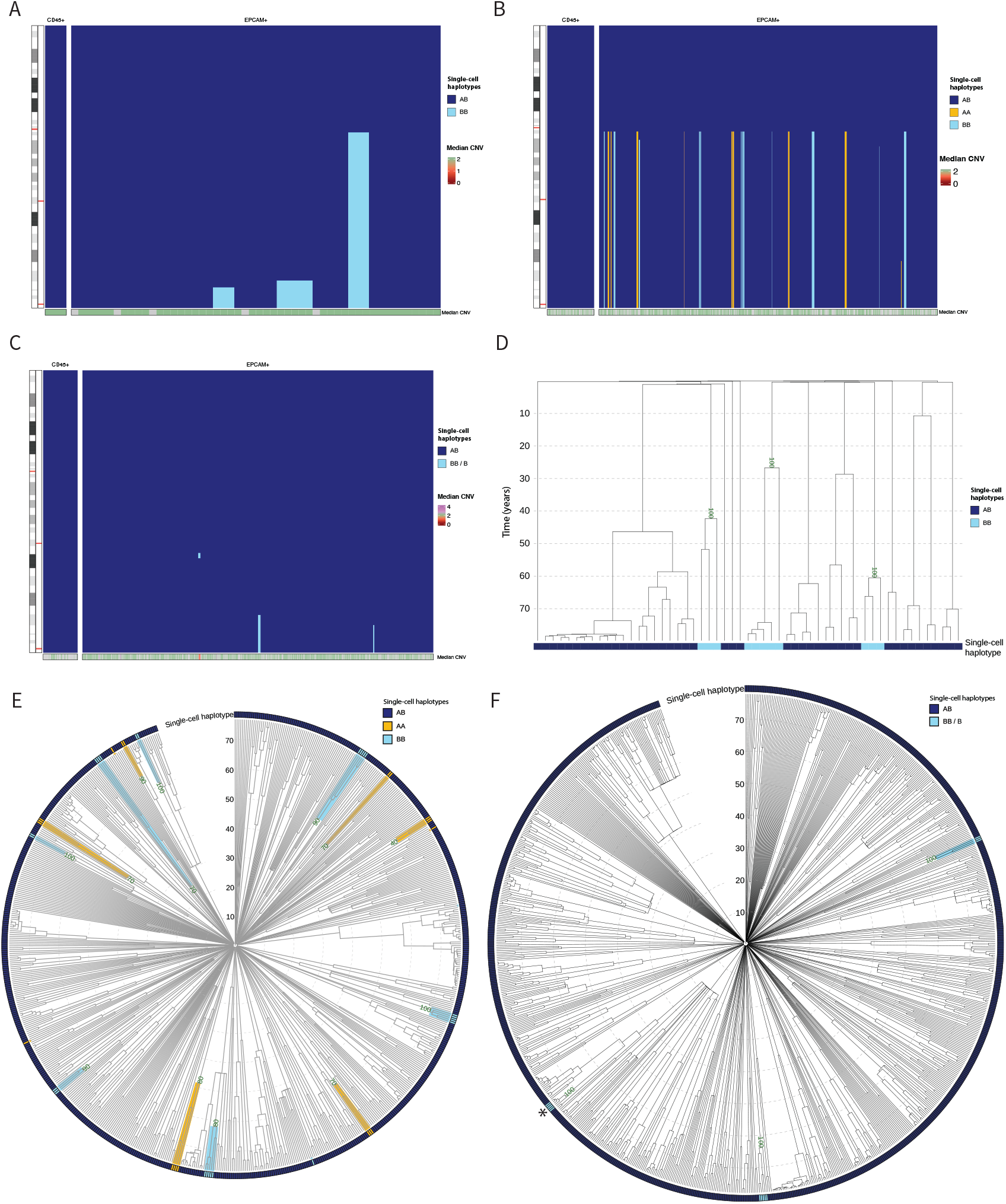
Loss of heterozygosity at chromosome 9q in donors eso01, eso03 and eso04. **A**, LOH map of chromosome 9 for all immune (CD45+) and epithelial (EPCAM+) cells from donor eso01. LOH segments were identified from the B-allele frequencies of germline heterozygous SNPs using a hidden markov model (Methods) and further manually correcting the interval exact breakpoints through visualization of the raw BAF profiles. Immune and epithelial cells are phylogenetically sorted within each group. The phylogeny was built after excluding SNVs detected in chromosome 9 to minimize the impact of residual germline variation (Methods). *NOTCH1* is located towards the telomeric end of the q arm of chromosome 9 (left, centromere in red on the chromosome, *NOTCH1* and *PTCH1* locations are indicated in red to the right of the chromosome). Single-cell haplotypes are assigned as AB (both parental alleles) or as having CNLOH, with AA or BB haplotypes. Median copy number across ∼50-kb non-overlapping windows is displayed in the bottom bar. Cells with LOH show median CNV for the affected region while wild-type cells show median for the whole chromosome arm. Median CNV values for cells with a coverage MAD higher than 0.2 (Supplementary Fig. 8D) are masked in grey. **B**, Same as in A for donor eso03. **C**, Same as in A for donor eso04. Three cells of this donor carry a small non-copy neutral LOH (B) not involving the *NOTCH1* gene in the middle of the large chromosome arm. **D**, Single-cell time-calibrated phylogeny of n = 55 cells from donor eso0, labeled by decade, indicated with dashed grey lines. Each leaf of the tree represents a single cell. Single cells showing LOH at any region of chromosome 9q implying a loss of the A haplotype (B haplotype) are annotated in light blue. Wild-type cells are represented in dark blue (AB). Bootstrap support of the MRCA for the independent LOH events is shown in dark green. **E**, Single-cell time-calibrated phylogeny of n = 699 cells from donor eso03, labeled by decade, indicated with dashed grey lines. Each leaf of the tree represents a single cell. Cells showing LOH at any chromosome 9q region are annotated in yellow (AA) or light blue (BB). Wild-type cells are represented in dark blue (AB). A circular layout was chosen to improve visualization of hundreds of cells. 59 cells with noisy BAF profiles were filtered out from phylogeny after visual inspection. **F**, Same as in E for donor eso04 (n = 874). * Marks a clone with loss of heterozygosity which is non neutral (B) instead of copy neutral (BB). A circular layout was chosen to improve visualization of hundreds of cells. 100 cells with noisy BAF profiles were filtered out after visual inspection.

**Supplementary Figure 10.**
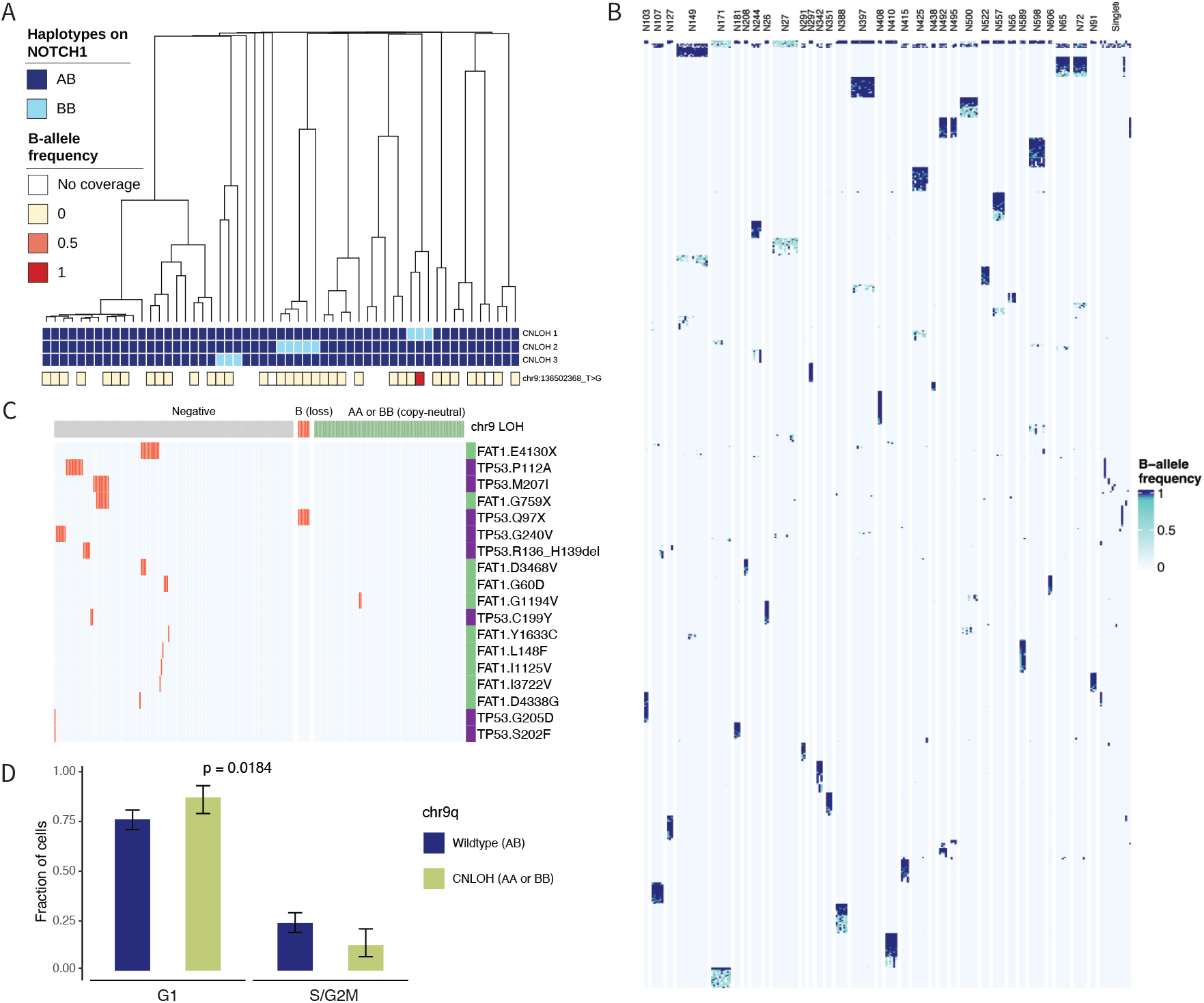
*NOTCH1* coverage and CNLOH replication phenotype. **A**, Phylogeny of eso01 including the three chr9q CNLOH events identified across cells (BB, indicated in light blue) and one somatic *NOTCH1* SNV detected in one cell carrying chr9q CNLOH, with B-allele frequency of 0.92 (bottom row). This suggests that the SNV may have occurred after acquisition of CNLOH on one of the chromosomes, although there is allelic imbalance. Targeted libraries were generated only for cells with sufficient amount of Illumina scWGS library left (highlighted with an outlined square, where BAF frequency is indicated). **B**, Heatmap of B-allele frequencies (BAFs) for somatic mutations on the long arm of chromosome 9 across cells in CNLOH clones, illustrating that CNLOH events arise independently. Each column represents a chr9q CNLOH-positive cell, grouped by clone as in Figure 7E. Only somatic mutations on chr9q that were not used for phylogenetic reconstruction and that show at least one cell with a BAF > 0.9 were included and clustered. SNVs with a BAF of 1 that are unique to each clone serve as validation that the clones arose independently: these mutations had to be present in heterozygosity in the most recent common ancestor of the cells in each clone before the CNLOH event, which subsequently doubled the BAF and are detected in homozygosis. **C**, Analysis of mutation status of *TP53* and *FAT1* together with copy-number status at chr9q in donor eso02. Left, plot of individual cells with at least one mutation in a driver gene but without chr9q CNLOH, with *TP53* or *FAT1* mutations indicated in red. Middle, all cells with a chr9q LOH event (deletion). Right, all cells with chr9q CNLOH (right). Red indicates cells with *TP53* or *FAT1* mutation, with the variant labeled on the right in purple or green, respectively. **D**, Proportion of epithelial cells in different cell cycle state (G1, S/G2M) for wild-type cells without chr9q CNLOH (AB) or cells with chr9q CNLOH (either the AA or BB haplotypes). *P* value obtained from a χ^2^ test.

